# Socially transmitted innovations in dynamic predator-prey systems

**DOI:** 10.1101/2022.09.13.507786

**Authors:** David W. Kikuchi, Margaret W. Simon

## Abstract

Individual behavioral variation is common, yet often we do not know how it is maintained. A potential explanation is that some behaviors must be acquired rather than genetically inherited. We investigate the social transmission of behavioral innovations, which can be key for the success of predator species, especially in contexts where environmental changes take place. We examine innovation in two classic predator-prey models. We assume that innovations increase predator attack rates or conversion efficiencies, or that innovations reduce predator mortality or prey handling time. We find that a common outcome of innovations is the destabilization of the system. Destabilizing effects include increasing oscillations or limit cycles. If either of these outcomes increases the risk of extinction, innovations that benefit individual predators may not have positive long-term effects on predator populations. Furthermore, as populations cycle, innovative individuals can be nearly eliminated, maintaining temporal behavioral variability. The destabilizing effects of behavioral innovations on predator-prey dynamics could have implications for biological invasions, urban populations, endangered species, and, more broadly, the maintenance of behavioral polymorphisms.

---

Individual variation in behavior is widespread among animal populations (Bolnick et al. 2003; Dall et al. 2004, 2012; Bell et al. 2009; Wolf and Weissing 2010; Araújo et al. 2011), yet we struggle to explain the mechanisms by which it persists (Dall et al. 2012). In some cases, evolution may maintain individual differences (Sinervo and Lively 1996; Martin and Pfennig 2009). In other cases, non-selective processes may maintain variation. These include variation in preferences learned during early development for food or mates (Darmaillacq et al. 2004; Hernandez et al. 2009). They also include behaviors that can be learned at any point, but which depend on finding another individual to learn from (Cavalli-Sforza and Feldman 1981; van der Post and Hogeweg 2006; Thornton and Clutton-Brock 2011). For example, bats learn socially about acoustic cues to locate anuran prey (Page and Ryan 2006) and capuchin monkeys learn from each other to crack open nuts using rocks as hammers (Ottoni and Izar 2008; Coelho et al. 2015). In this study, we examine such socially-mediated learning. Specifically, we focus on behavioral innovation, in which an animal learns a novel behavior that allows it to exploit a novel resource, or exploit an existing resource in a novel way (Greenberg 2003).

The aim of our study is to contribute to our basic understanding of reciprocal effects between individual variation and population dynamics. This question has applications to how populations respond to invasive species, such as predators that learn how to devour toxic, invasive cane toads in Australia (Beckmann and Shine 2011; Parrott et al. 2019). It also has relevance for species undergoing range expansion. For example, Common Mynas on their invasion front in Israel are more innovative and also more accepting of novel foods than in their native range (Cohen et al. 2020). Finally, this question speaks to how species come to occupy new niches. Great tits in Hungary, for example, have adapted themselves to consuming bats when their traditional food resources are scarce (Estók et al. 2010).

Innovations that spread from individual to individual cannot be studied in the same frameworks as genetically based innovations, or those that depend only upon how a single individual interacts with its environment (i.e. reaction norms). Instead, we need to explicitly consider the mechanisms that underlie information spread. There are many mechanisms by which an innovation might pass between individuals. They include various forms of vertical (parent to offspring), horizontal (e.g., within a stage/age class), and oblique transmission (between non-parent adults and juveniles) (Cavalli-Sforza and Feldman 1981; Denton et al. 2020). The spread of social information can be further determined by the specific structure of social networks (Farine et al. 2015; Shultz et al. 2017), which may themselves respond to the fluctuating value of social information (Cantor et al. 2021; Romano et al. in press). In this study, we focus on innovations that follow horizontal transmission in populations of randomly mixing individuals. This assumption relates innovation spread to some well-studied models for the transmission of disease.

The metaphor of disease yields a simple explanation for how horizontally transmitted innovations could result in a behaviorally variable population (Cavalli-Sforza and Feldman 1981). In disease ecology, individual variation in infection status is maintained when the disease spreads on the same timescale as demographic rates (births and deaths) (Anderson and May 1980). In the same way, behavioral variability could be maintained when social learning is slow enough that an individual has a chance of not acquiring a behavior during its lifetime (Cavalli-Sforza and Feldman 1981). For example, it may take a significant portion of a capuchin monkey’s life to learn how to crack a nut with a stone hammer; thus, not all monkeys will learn (Ottoni and Izar 2008; Coelho et al. 2015). However, in their most basic (and familiar) formulations, these models do not consider how a disease affects an individual’s reproductive rate or mortality risk. When disease affects fitness, it can create feedback with population size (see Ashby et al. 2019 for discussion in an evolutionary context). Similarly, social information might not only spread on the same timescale as demographic rates, but could also *change* those rates (Ihara and Feldman 2004; Thorogood et al. 2018; Whitehead et al. 2019). Furthermore, as host populations grow or shrink, the proportion of infected and uninfected individuals can vary too because disease spread depends on population size (Anderson and May 1980). Behavioral innovations could simultaneously influence the number of individuals in a population, and its behavioral variability.

Innovations in foraging are by far the most widely documented types of novel behaviors in the literature (e.g. Lefebvre et al. 1997; Overington et al. 2009). There is evidence for their social transmission in several systems (Hämäläinen et al. in press). These include humpback whales using a novel lobtail feeding technique (Allen et al. 2013), and wild populations of birds learning about novel prey phenotypes, where they are predicted to impact prey evolution (Thorogood et al. 2018). Innovations have also been connected to extinction risk (Ducatez et al. 2020). This makes them a logical target for theoretical investigation.

Earlier theoretical studies have examined how social information transmitted on short timescales (e.g. alarm calling) affects populations in predator-prey systems (Gil et al. 2018, 2019; Tóth 2021). Our study examines a longer timescale where predators acquire an innovation that positively affects some component of their fitness for the remainder of their lifespan. Recently, Borofsky and Feldman (2022) examined how such innovations among predators impact populations of prey; however, in their study, predator populations remained constant. Here, we allow both predator and prey populations to vary dynamically. Throughout our study, we build on basic equations from population biology and epidemiology. We begin by examining the spread of innovations in a simplistic, abstract system. Then we extend our exploration to more realistic, complex models.

## Conceptual framework

We study a single species of predator that feeds upon a single species of prey. We assume a single innovation spreads within the population of predators, but that all predators are otherwise identical in their demographic rates. All predators are born without knowledge of the innovation (we call such individuals “naïve”), and must acquire it from another predator that has already learned the behavior (we call such individuals “innovators”). Both naïve and innovative individuals consume resources and give birth only to naïve individuals. The innovation spreads according to the law of mass action, where encounters between naïve and innovative individuals lead to the spread of the innovation in proportion to their population densities (i.e. horizontal transmission, Cavalli-Sforza and Feldman 1981). Figure 1 provides a conceptual overview of the models we develop here (Fig. 1A), and highlights our main results (Fig. 1B).

**Figure 1.**
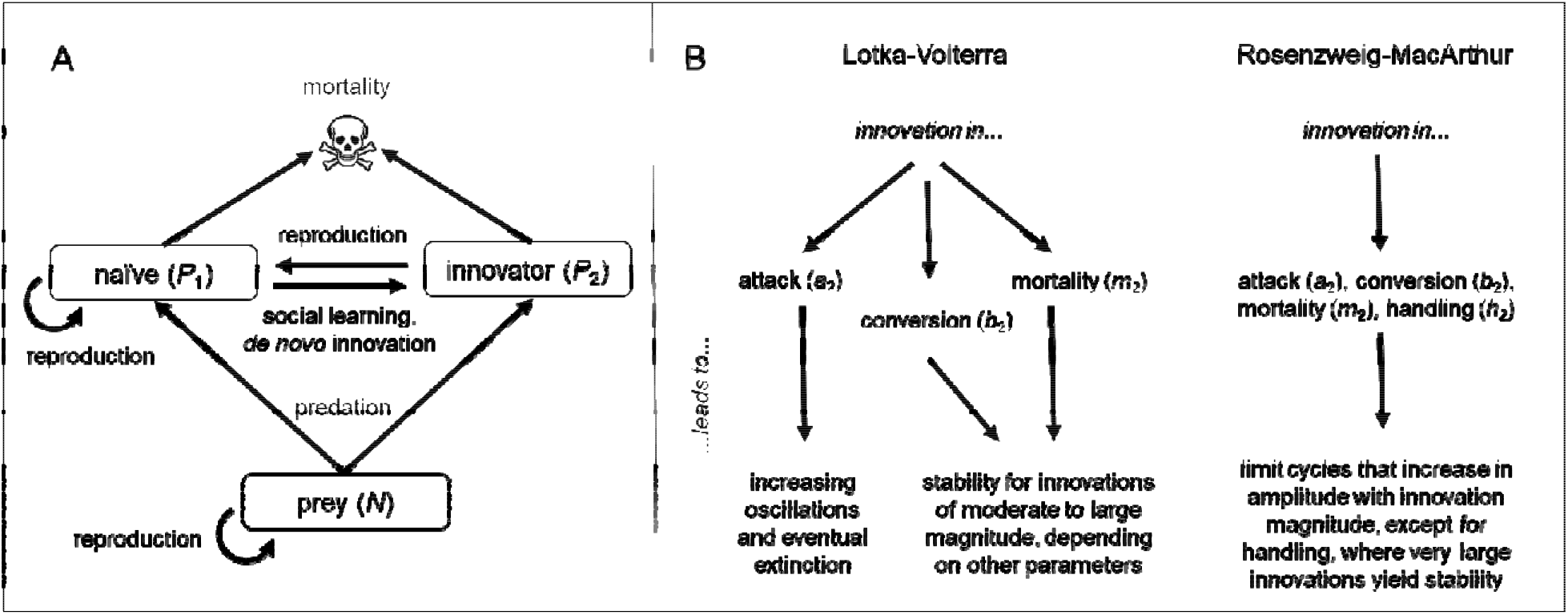
A) Structure of the predator-prey community assumed for all the models developed in this study. Common features include: prey that occupy the bottom trophic level, predators that reproduce by converting their prey into new naïve predators, social learning and *de novo* innovation that transforms naïve predators into innovators, and constant per capita predator mortality. B) Summary of results generated when innovations take place under two alternative assumptions about prey demography and predator foraging efficiency (Lotka-Volterra and Rosenzweig-MacArthur). In Lotka-Volterra systems, innovations eliminate the neutral stability exhibited by the classic model. Instead, attack rate innovations cause eventual extinction. Innovations in conversion and mortality cause either stability or limit cycles, depending on their magnitude and other parameter values. In Rosenzweig-MacArthur systems, innovations above a certain magnitude cause limit cycles, and larger-magnitude innovations tend to create more extreme limit cycles. The exception to this is handling time, where stability can be restored with innovations of very large magnitude.

To develop intuition for innovations that spread by mass action, we first describe how innovations spread without explicit consideration of the prey. Ignoring prey may not be realistic in the context of predation, but it lets us quickly see how innovations might spread in general. It might also be a reasonable representation of a system with a constant amount of prey that do not respond to changes in predator population size, such as milk bottles that are delivered daily (which tits have learned to open Lefebvre 1995). This model formulation is consistent with Cavalli-Sforza and Feldman’s (1981) description of the spread of a novel behavior in an isolated population. If we assume fast information spread relative to demographic rates, and no effect of innovation on demographic rates, then the rate of change in the proportion of innovators can be described by *sQ*_1_*Q*_2_, where *s* is the rate at which a naïve individual acquires the innovation if it contacts an individual bearing the innovation (we assume *s* < 1). *Q*_1_ and *Q*_2_ are the proportion of naïve and innovative individuals, respectively. The resulting time series (proportion of innovators in the population versus time) is a sigmoidal curve that has an asymptote at *Q*_2_ = 1 (i.e. the entire population becomes innovative). This is also predicted by epidemiological models of the same form.

Cavalli-Sforza and Feldman (1981) also describe novel behaviors that spread slowly relative to demographic rates. Assuming population size is constant, with individuals dying (and being born) at per capita rate *g*, there is a stable equilibrium where the proportion of innovators is (*s – g*)/*s*. As the rate of social learning increases, the proportion of innovators increases; as the rate of population turnover accelerates, the proportion of innovators decreases because naïve individuals die before they can learn. Cavalli-Sforza and Feldman (1981) additionally examine cases where individuals can reject an innovation and become resistant to it – there may be biological analogues of this in animal populations, but we do not explore this possibility here.

### A simple dynamic system of social learning

The epidemiological model with population turnover makes a good departure point for exploring innovations that affect demographic rates. Here we assume that an innovation has a beneficial effect on predator demographic rates, although as seen above, neutral (or even deleterious) innovations can spread by the same mechanisms. Models of disease in natural populations have found that hosts can be limited by a pathogen (e.g. Anderson and May 1980). However, our assumption that innovations have positive effects on demography creates the slight problem that a population with an innovation could grow without bound. A natural way to limit a population in which an innovation spreads is to let the population be limited by the availability of prey. Instead of a single parameter *g*, we divide predator population turnover into different components that represent the rate of encounter with prey *a*, the efficiency with which encountered prey are converted into new predators *b*, and predator mortality *m*. Additionally, we relax the assumption of a constant population size so that instead of dealing with proportions of the population, we use densities. We assume that the density of prey *N* increases according to the function *R*(*N*). We also assume prey are captured according to *F*(*P*_*i*_, *N*), which describes the functional response. We use *P*_*i*_ ∈ (*P*_1_,*P*_2_) to represent the densities (rather than proportions) of naïve (*P*_1_) and innovative (*P*_2_) individuals. Prey dynamics are therefore described by the equation:

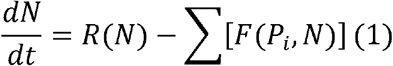

All predators are born naïve, so both naïve and innovative predators contribute to the density of naïve predators, whereas innovators are only produced through contact with naïve predators. Naïve and innovative predators convert captured prey at rates *b*_1_ and *b*_2_, respectively (*b*_2_ > *b*_1_). We consider the possibility that naïve predators may innovate via non-social learning, i.e. *de novo* innovation. This is represented by the function *D*(*P*_1_) in equations (2) and (3) below. The function *s*(*P*_1_,*P*_2_) describes the rate at which naïve individuals become innovative upon contact with an innovator, i.e. learn socially. We assume that *s*(*P*_1_,*P*_2_) is independent of the magnitude of the innovation. We define the magnitude of an innovation as the difference between parameter values for innovative and naïve individuals, e.g. *b*_2_ – *b*_1_. Finally, some types of innovations may reduce predator mortality, so we assume separate death rates *m*_1_ and *m*_2_ (*m*_2_ < *m*_1_). Therefore, the predator population dynamics are described by:

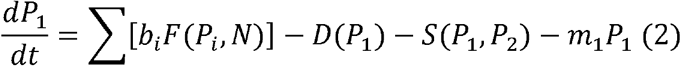

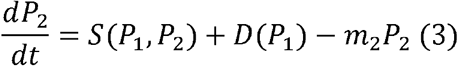

Equations 1 – 3 assume that all parameters are positive.

First, we analyze the model under Lotka-Volterra assumptions (Table 1). Prey are not self-limiting, such that *R*(*N*) = *rN*, where *r* is the intrinsic rate of increase. We also assume that predators follow a Type I functional response (Holling 1959, 1965) where *F*(*P*_*i*_,*N*) = *a*_*i*_ *P*_*i*_ *N*. Under this assumption, prey are captured at rate *a*_1_ when they are encountered by naïve predators, but at rate *a*_2_ when they are encountered by innovators (*a*_2_ > *a*_1_), which means they take a negligible amount of time to handle their prey. We assume social learning takes the form *s*(*P*_1_,*P*_2_)= *sP*_1_*P*_2_. Additionally, we assume *de novo* innovation does not occur, i.e. *D*(*P*_1_) = 0.

**Table 1.** Forms of prey growth *R*(*N*), predator functional response *F*(*P*_*i*_, *N*), and social learning *s*(*P*_1_,*P*_2_) used in this study.

| Function | Form | Interpretation |
| --- | --- | --- |
| $R(N)$ | $rN$ | Prey are not self-limiting, growing at intrinsic rate $r$ (Lotka-Volterra assumption). |
| | $\frac{rN(k - N)}{k}$ | Prey are self-limiting with carrying capacity $k$ (Rosenzweig-MacArthur assumption). |
| $F(P_i, N)$ | $a_i P_i N$ | Predators exhibit a Type I functional response (Lotka-Volterra assumption). |
| | $\frac{a_i P_i N}{1 + a_i h_i N}$ | Predators exhibit a Type II functional response (Rosenzweig-MacArthur assumption). |
| $D(P_1)$ | 0 | No *de novo* innovation. |
| | $dP_1$ | *de novo* innovation occurs in proportion to the density of naïve predators. |
| | $dP_1 N$ | individuals are only likely to innovate *de novo* when encountering a prey item. |
| $S(P_1, P_2)$ | $sP_1 P_2$ | Social learning depends only upon a naïve individual observing an innovative individual. |
| | $sNP_1 P_2$ | Social learning depends on a naïve individual observing an innovator interact with a prey. |
| | $\frac{sa_2 NP_1 P_2}{1 + a_2 h_2 N}$ | Social learning depends on a naïve individual observing an innovator handle prey. |

Although the assumptions of the Lotka-Volterra equations may lack realism for many empirical systems, they let us arrive at some analytical conclusions. Subsequently, we relax these assumptions to consider alternative forms of *R*(*N*), *F*(*P*_*i*_, *N*), *D*(*P*_1_), and *s*(*P*_1_,*P*_2_) (Table 1).

We seek conditions under which an innovation can initially spread in (invade) a population. In epidemiology, the number of susceptible individuals infected by a diseased individual is called *R*_0_ (see e.g. Cavalli-Sforza and Feldman 1981). This threshold must exceed 1 for the disease to increase in the population, and depends on the number of susceptible individuals available to be infected (Anderson and May 1986). Analogously, there are thresholds that the population density of naïve individuals must achieve for innovations to spread 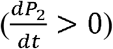. If we rearrange equation (3) (with *s*(*P*_1_, *P*_2_)= *sP*_1_ *P*_2_ and *D*(*P* _1_) = 0), we see that^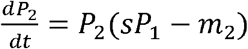^. For a positive rate of change in innovator abundance, the condition 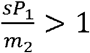 must be met (i.e.,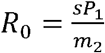). Innovators must both encounter enough naïve individuals, and pass the innovation on to them, before dying. Whether they achieve this depends on *s*. Additionally, innovations that decrease mortality rate will spread more easily because they lower the minimum number of naïve individuals required for sustained innovation spread, much like a less virulent disease. From equation (1), we can also see that at equilibrium (denoted by a “*” superscript) in the absence of innovation (i.e.,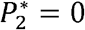), that 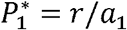 and *N* ^∗^ = *m*_1_/(*a*_1_*b*_1_). For an innovation to innovation to invade a population of solely naïve predators, 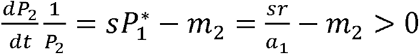(from equation 3), which means that 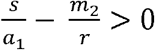 must be true for successful innovation spread (i.e., *P*_2_ > 0). This is also a necessary condition for an equilibrium where both types of predators coexist with their prey (at such an equilibrium 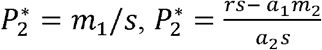, and 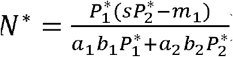). Biologically, this means that for an innovation to spread and be maintained in the population, prey reproduction and predator social learning must be fast relative to the attack rate of naïve predators and the death rate of innovative predators. If this condition is not met, the population of innovators cannot spread nor be sustained.

The equilibria of this system are a marked departure from both the epidemiological models and the Lotka-Volterra equations upon which it is built. The trivial equilibrium where all species are extinct is unstable, as is the equilibrium where only the prey and naïve predators exist – this is simply the neutrally stable Lotka-Volterra model. It is difficult to make meaningful general statements about the stability of the equilibrium point where all three types coexist due to the large number of parameters involved. However, in this full system, it is assumed that the innovation beneficially affects all demographic parameters. This may not be realistic for most innovations. We therefore examine specific scenarios in which an innovation causes only a single pair of parameters to differ between naïve and innovative individuals. First, we consider innovations that increase attack rate such that *a*_2_ > *a*_1_, but *b*_1_ = *b*_2_ and *m*_1_ = *m*_2_. This could represent an innovation that elevates foraging success, such as imitating a successful search strategy or habitat choice. We find that when an innovator’s attack rate *a*_2_ is greater than the naïve attack rate *a*_1_, it generates oscillations of increasing amplitude, eventually leading to extinction. This is a consequence of the innovation acting directly on both predator and prey populations, resulting in positive feedback between the two. The larger the innovation is, the more quickly either the predator or the prey reaches extinction.

Next we examine the case where an innovation increases conversion efficiency (*b*_2_ > *b*_1_; *a*_1_ = *a*_2_, *m*_1_ = *m*_2_), as might happen when predators learn how to more efficiently exploit prey tissues. For example, some species of rodents cut and store toxic plants, and only return to consume them after the plant’s toxicity has decreased (Glendinning 2007). This behavior presumably increases the net value of toxic plants as a resource. Innovations to conversion rate cause the system to reach a stable equilibrium, or to exhibit limit cycles. Dropping subscripts on *a* and *m*, a necessary condition for stability at equilibrium is 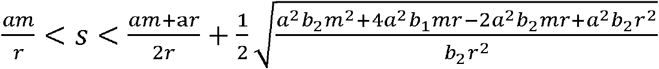. In other words, the system is stable when social learning is moderate in speed; predator populations are gradually supplemented by reproductive output of innovators back into the naïve population to such a degree that the whole predator population does not collapse at lower prey abundances (Fig. 2A). Innovative predators can be thought of as storing energy for release back into the naïve predator population, as though a segment of the population were in a temporal refuge buffered from starvation. However, when social learning occurs too slowly or too quickly, the predator population either does not receive a critical level of supplement, or receives too much, such that it overshoots prey abundances. Both of these cause limit cycles (Fig. 2B). This condition can also be framed in terms of *b*_2_. If *a* > *s*, only *b*_1_ < *b*_2_ is required for stability (i.e. any innovation). When *a* < *s*, stability requires 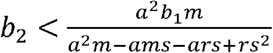 (i.e. there is an upper bound on innovation magnitude beyond which the system is not stable).

**Figure 2.**
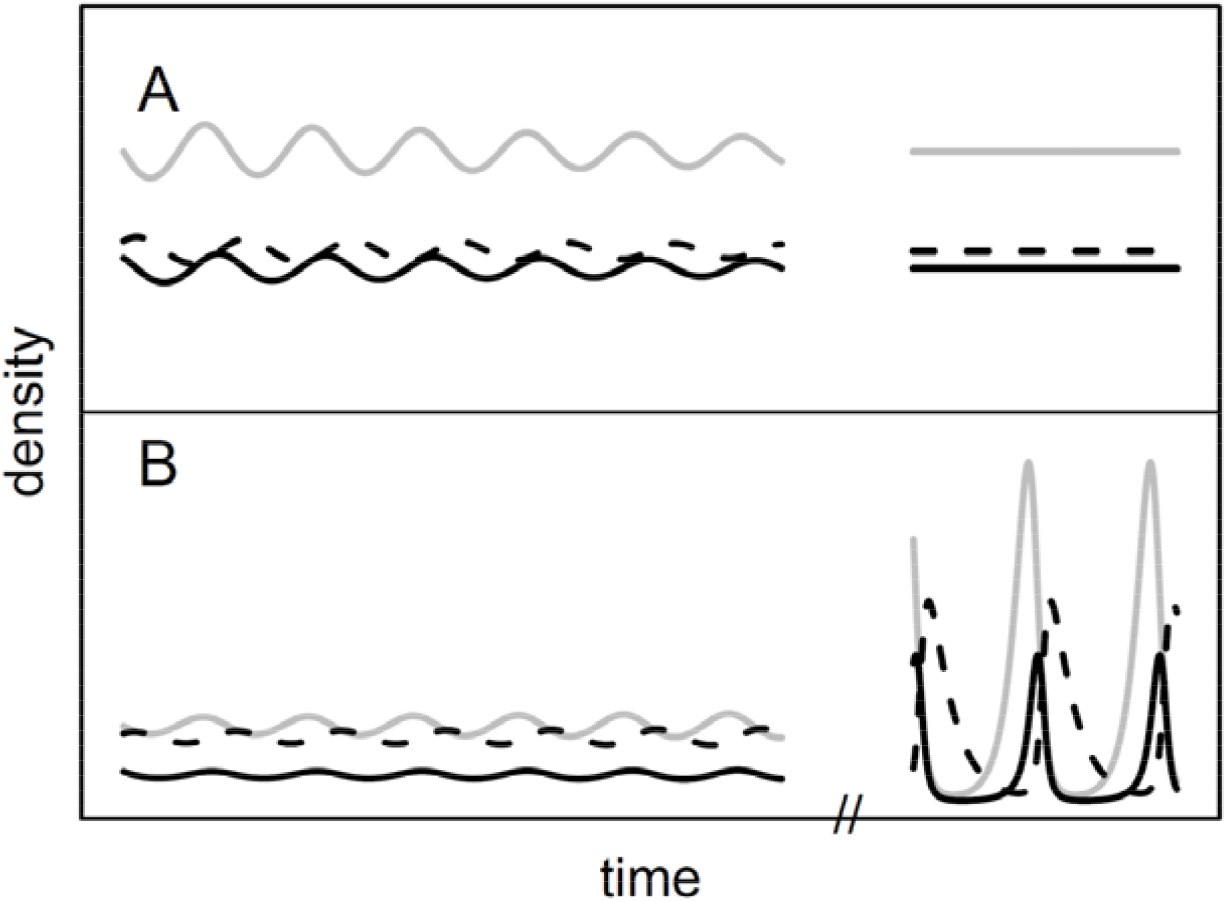
Lotka-Volterra outcomes of conversion rate innovations depend on social learning. A) moderate social learning speed yields a stable equilibrium over time (*s* = 0.16). B) Social learning speed exceeds a critical threshold, destabilizing the system into limit cycles (*s* = 0.25). Gray = prey; solid black = naïve predators; dashed = innovative predators. In both panels, other parameters are *a* = 0.15, *b*_1_ = 0.4, *b*_2_ = 0.7, *m* = 0.2, r = 0.4.

For completeness, we also consider innovations that decrease mortality (*m*_2_ < *m*_1_; *a*_1_ = *a*_2_, *b*_1_ = *b*_2_). These are not directly related to foraging, but still constitute innovations when they arise from individuals learning to utilize their environments in a new way (resource use in a broad sense). This could include new shelter-use behaviors like hiding in cars (Cauchard and Borderie 2016) or nesting in artificial structures (Lowry et al. 2013; Dias et al. 2017). Predator survival could also increase in response to social learning to avoid dangerous or toxic prey (other than the focal prey modeled here) (e.g. Thorogood et al. 2018). Under the assumption that *m*_2_ < *m*_1_, dropping other subscripts, one condition for stability is that 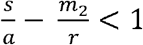. From our analysis of *R*_0_ and requirement for a biologically feasible equilibrium (above), we can also place a lower bound on this expression: 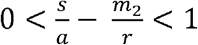.

This means that for stability, *m*_2_, *a*, and *s* have both upper and lower bounds (in some cases 0 for *m*_2_). Additionally, *r* may have both an upper and lower bound, or only a lower bound depending on the relationships between the other parameters (see Supplementary Mathematica notebook for details).

Generally, the spread of innovations in conversion rate and mortality reduces the tendency of predator populations to crash during periods of low prey abundance, resulting in a weaker prey boom at the start of the subsequent cycle. However, when social information spreads too quickly, this damping effect is limited, and instead a stable limit cycle results. This is similar to the effect on the Lotka-Volterra model when a *prey* species uses multiple habitat patches: as long as predators disperse randomly between the two patches, stability results (Holt 1984). The system only reverts to neutral stability when predators disperse with infinite speed (Holt 1984). Likewise, in our system, if social information spreads at such a fast rate that demographic parameters are irrelevant, this separation of timescales returns the system to the original Lotka-Volterra equations.

The relationship between resource abundance and information spread means that, rather than a single predictable polymorphism in behavior (as with the traditional SI epidemiological model in which total population size is assumed constant), the proportion of innovative individuals depends on resource productivity (*r*). This predicts that across a productivity gradient, the proportion of innovative predators will increase (Fig. 3). In general, because of the influence of resource growth rate on population dynamics at higher trophic levels, we should expect social learning to be impacted by the productivity of the system in which it is embedded.

**Figure 3.**
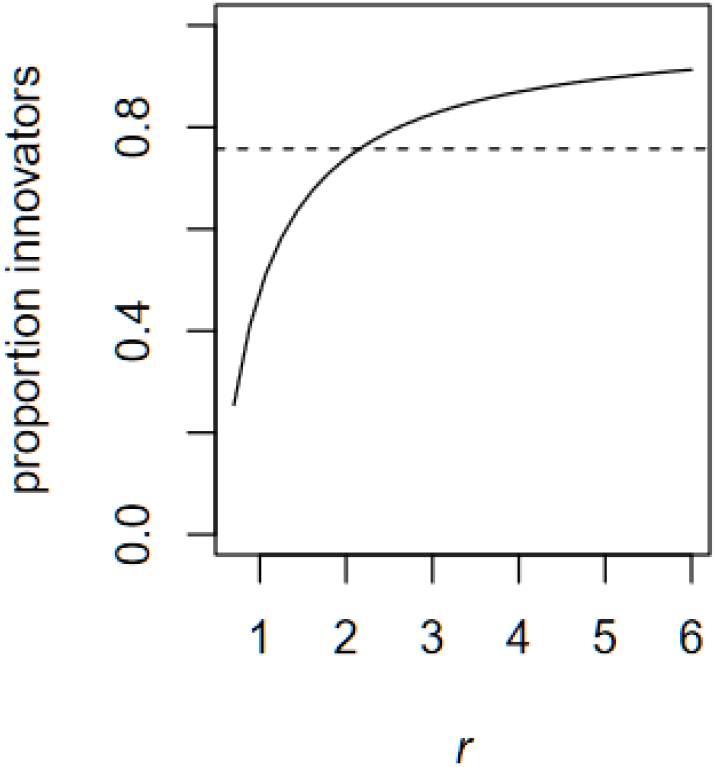
The proportion of innovative individuals in a population. Solid line = result from the Lotka-Volterra system that includes conversion rate innovators (*b*_2_ > *b*_1_). Dashed line = population with social information transmission and vital dynamics, but no consequences for population densities (i.e. a vital rate SI model with *d* = 0.5 and *s* = 0.16). Here *a* = 0.167, *b*_1_ = 0.5, *b*_2_ = 0.55, *m* = 0.5, *s* = 0.16.

To examine the influence of *de novo* innovation on our system’s dynamics, we investigated two modes of non-social learning (Table 1). Initially, we let *s*(*P*_1_,*P*_2_)= 0 in equation (3), so that social learning does not occur. First, we examine *de novo* innovations that occur in proportion to the density of naïve predators, such that *D*(*P*_1_) = *dP*_1_, where *d* is a constant. This could represent the adoption of a novel foraging strategy independent of encounters with prey, such as foraging in hedges rather than fields. Under this assumption, stability analyses reveal that innovations to attack rate result in instability, whereas innovations in conversion rate and mortality always yield stability at the lone non-trivial equilibrium (see Supplementary Mathematica notebook for details). Second, we assume that *de novo* innovation requires that a predator encounter a prey, which we represent by letting *D*(*P*_1_) = *dP*_1_*N*. In this case, at the nontrivial and biologically feasible equilibrium, instability results from innovations in attack rate and conversion rate; mortality innovations yield stability (see Supplementary Mathematica notebook for details). Next, we examine the system’s behavior when predators can either innovate *de novo* (*D*(*P*_1_) ≠ *d0*) or use social learning *s*(*P*_1_,*P*_2_)= *sP*_1_*P*_2_. Analytically, such systems are difficult to interpret. However, numerical analysis reveals that when *D*(*P*_1_) « *s*(*P*_1_,*P*_2_), the system behaves primarily like the one where social learning takes place alone (Fig. S1). Therefore, the ecological effects of *de novo* innovation are likely to be swamped by social learning in systems that meet Lotka-Volterra assumptions.

The above analyses illustrate the feedback between dynamic populations and social learning, and they give us some intuition into how different quantities influence one another. Yet the assumptions of unbounded prey growth, and linear relationship between prey abundance and predator consumption rate, are poorly supported for many systems. Below, we add more details to allow our model to apply to a far larger proportion of predator-prey ecologies.

### Innovation spread in the Rosenzweig-MacArthur model

Two classic assumptions in consumer-resource models are that resources (when alone) exhibit sigmoidal population dynamics (reaching a carrying capacity at high abundance), and that individual predators exhibit saturating functional responses to prey abundance (e.g. Holling Type II) (Rosenzweig and MacArthur 1963; Rosenzweig 1969; for an overview, see McPeek 2022). With self-limiting prey, we assume that 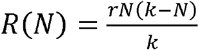, where *k* is the carrying capacity of the resource. In a Type II functional response,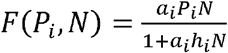, where *h*_i_ is the time to handle prey (Table 1).

Under these assumptions, (1) – (3) do not yield analytical equilibria, nor can their stability be treated with analytical methods. We can, however, predict how innovations are likely to affect the system by comparing them to the nullclines of a system without innovation (Rosenzweig and MacArthur 1963). Systems with a single self-limiting prey and a single class of predator with Type II functional response have a stable equilibrium when the predator nullcline is to the right side of a “hump” in the prey nullcline (Fig. 4A). If the predator nullcline falls on the left side of a hump, stable limit cycles result (Rosenzweig 1969) (Fig. 4B).

**Figure 4.**
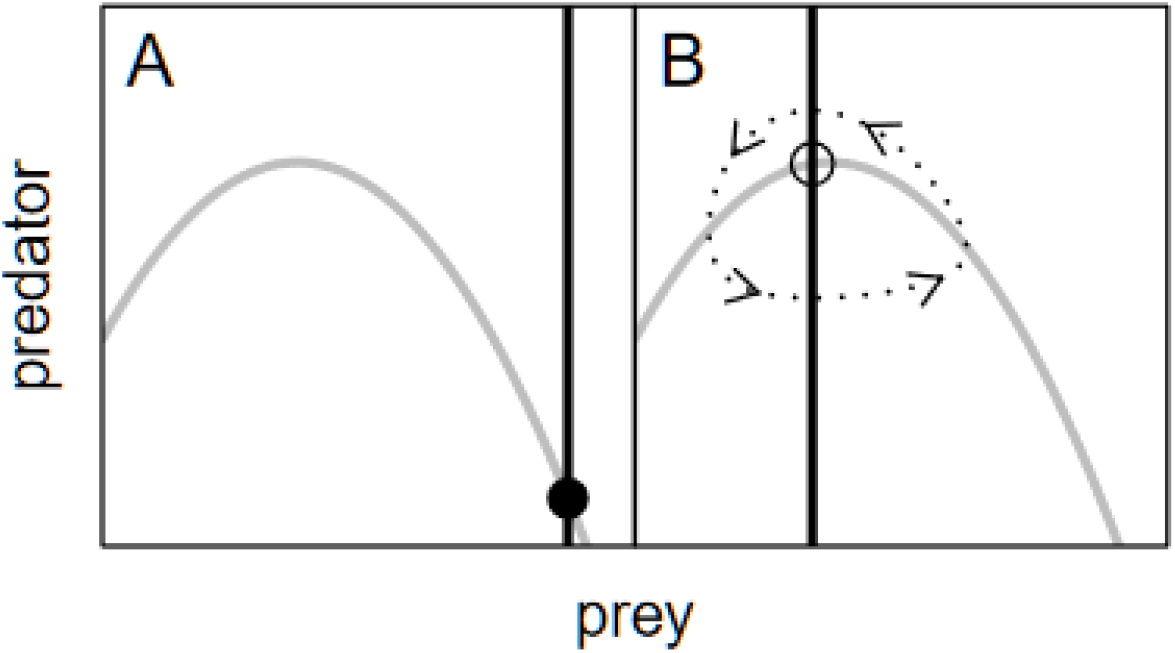
Prey nullclines (gray curves) and predator nullclines (black vertical lines) in systems where prey are self-limiting and predators use Type II functional responses. A) stable equilibrium shown by the black dot (e.g. without innovation). B) Limit cycles following the dotted line around an unstable equilibrium shown by the open dot (e.g. where all individuals have acquired an innovation).

Because they can be sustained by fewer prey, predators that are more efficient at exploiting their prey will have nullclines further to the left than less efficient predators. Beneficial innovations may therefore create segments of the predator population that tend to push the system away from stability, towards limit cycles.

To test this hypothesis, we examine systems where populations of only naïve individuals and their prey have stable equilibria. For an innovation to initially invade a system at stable equilibrium, the naïve predator population must be above the threshold density 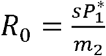, where again 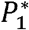 denotes equilibrium density. This suggests that predator populations living a marginal existence may be the least likely to foster innovations that would help them succeed (i.e., if the vertical line in Fig. 4A were even further to the right). An innovation that reduces mortality (*m*_2_) would increase *R*_0_ at a given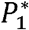, and so might be more likely to invade a small predator population than other innovations. It is easy to see why mortality innovations are special by considering extreme cases. If a predator discovers an innovation that makes it immortal, it will eventually encounter another predator to share the discovery with no matter how long it takes. By contrast, if a predator discovers an innovation to massively increase its rate of converting prey, it may produce a flood of naïve offspring but die before any of its children can learn its secret.

Under parameter values where the innovation can invade, we use bifurcation analyses to see whether larger-magnitude innovations shift the system from stable equilibria to limit cycles. Across innovation types, the bifurcation analyses show that stable systems are destabilized by the spread of innovations that have large positive effects on innovator performance (Fig. 5). Innovations that confer only a small advantage do not destabilize the system. All innovations large enough to destabilize the system shift the trophic structure of the community, such that predator populations are higher relative to prey (shown on a log scale in Fig. 5). With innovations in handling time, the amplitude of cycles increases, then decreases as innovations bring handling time close to zero (Fig. 4). This occurs because the shape of the prey isocline loses its “hump” as handling time approaches 0. In other words, the predator becomes “stable” in its foraging behavior because its functional response becomes approximately linear (sensu Abrams and Holt 2002). With other innovations, populations remain unstable as innovation magnitude increases (Fig. 5).

**Figure 5.**
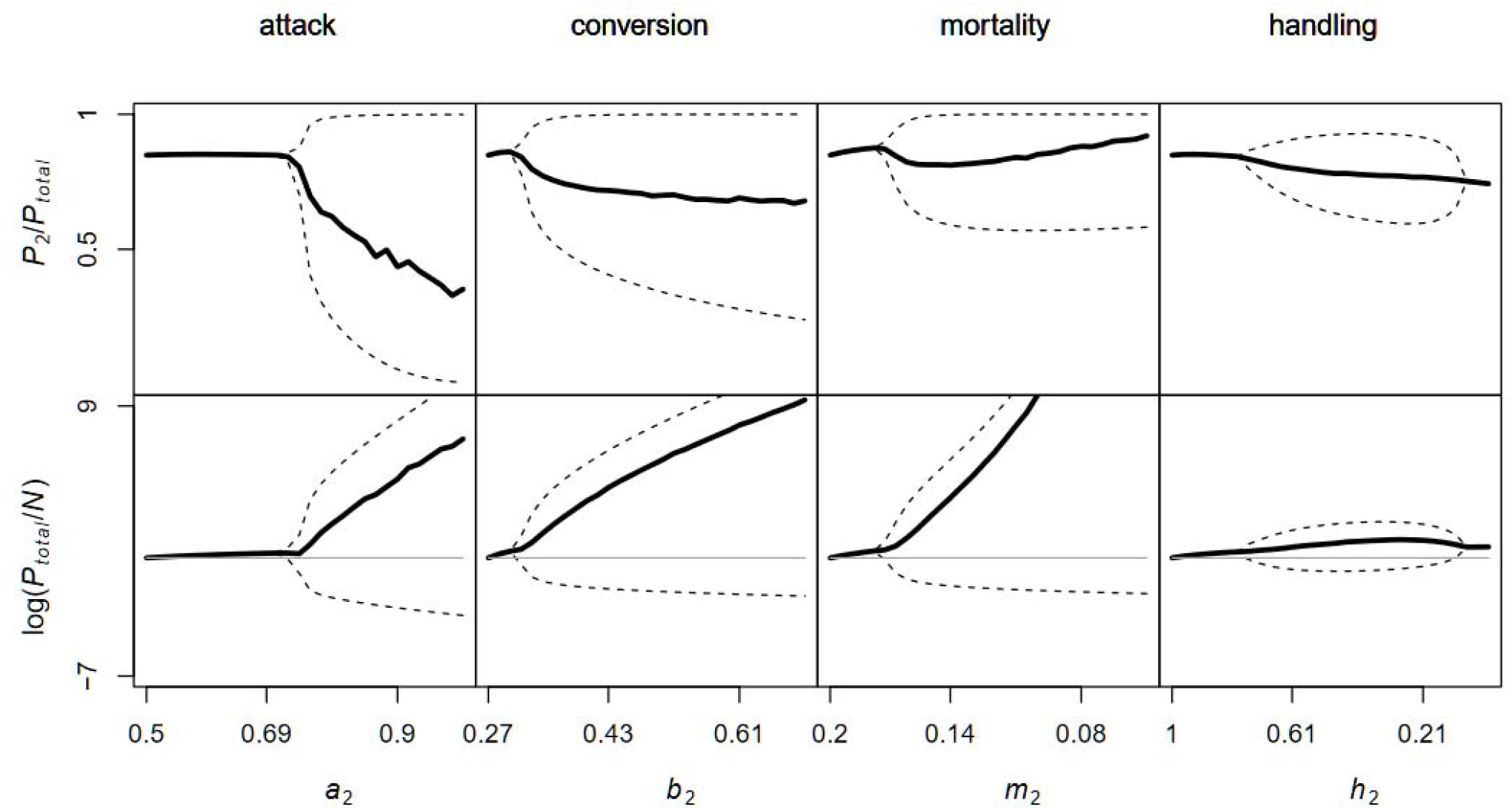
The effect of increasing innovation magnitude on the proportion of innovative individuals in the population (top) and the log ratio of predator:prey (bottom) in the Rosenzweig-MacArthur system. Black line = time-averaged mean. Dotted line = range of limit cycles. Gray line = population equilibrium in absence of innovation. Except for the varying parameter on the abscissa, naïve and innovative predators shared the same parameter values: *a* = 0.5, *b* = 0.27, *m* = 0.2, *h* = 1, *s* = 0.2, *r* = 2, *k* = 10.

However, perhaps surprisingly, the more advantageous an innovation in attack rate or conversion efficiency is, the lower the average proportion of innovators in the predator population. This happens because there is a critical density of naïve individuals required for *R*_0_ to exceed 1, allowing an innovation to spread. The innovation has difficulty spreading during periods of low predator density, so during these times most predators are naïve. A fluctuating proportion of innovators is predicted in a population with cyclic dynamics because the cycles include periods of abundance where *R*_0_ < 1. Indeed, a parallel exists in wildlife disease, where the population cycles of great gerbils drive periodic outbreaks of the bacterium that causes bubonic plague (Davis et al. 2004). Mortality innovations produce slightly different patterns, with the proportion of innovators increasing at very large magnitude innovations (because innovators live a very long time).

As with the simple Lotka-Volterra system above, it is also helpful to have predictions about how a single type of innovation affects population dynamics and behavioral variability across an ecological gradient, e.g., resource productivity or general proclivity towards social learning. We examine the influences of *r* and *s* on how a population is impacted by the spread of an innovation.

Stable equilibria only exist at low values of *r* and *s*, which have qualitatively similar effects on stability. Once limit cycles emerge, their amplitudes typically increase with increases in *r* or *s* (Fig. 6A). Trends in limit cycles are similar to those displayed in Fig. 6A regardless of innovation type (i.e. *a*_2_, *b*_2_, *m*_2,_ or *h*_2_,), or whether we examine predator and prey populations separately, predator:prey ratios, or the proportion of innovative:naïve predators. The mean proportion of innovators in the population also increases with *r* and *s* (Fig. 6B). This happens because higher prey populations support more predators, and larger *s* increase the rate of social information spread. Both of these increase *R*_0_, so the innovation is adopted by more predators.

**Figure 6.**
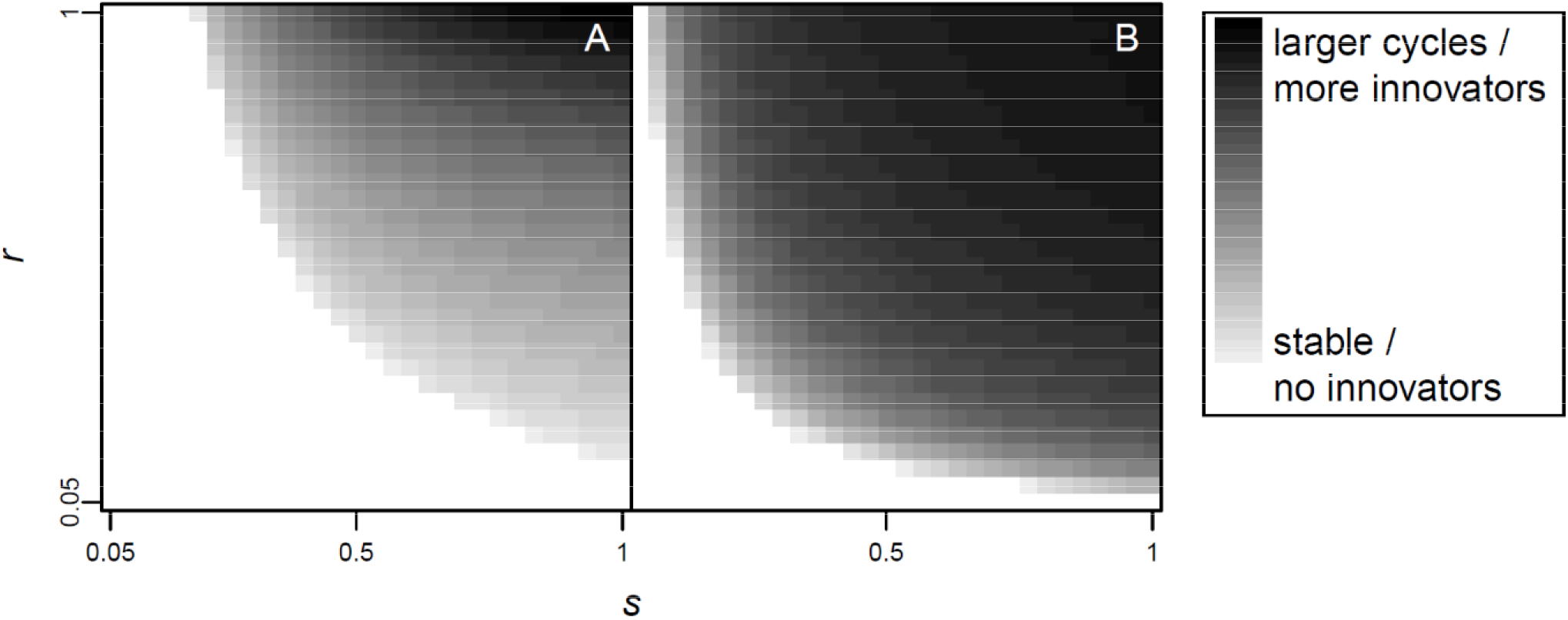
A) Bifurcation analyses of the total predator population size across a range of *r* and *s* values for attack rate innovations in the Rozensweig-MacArthur system. Increasing either parameter shifts the system from stability towards increasingly high-amplitude limit cycles. B) Corresponding plot showing the mean proportion of the population composed of innovators increases with *r* and *s*. In both panels, *a*_1_ = 0.5 and *a*_2_ = 0.75; all other parameter values the same as in Fig. 5.

### When social learning depends on observing an innovator interact with prey

So far, we have considered only mass action spread of innovations between two predators. Many other mechanisms of spread exist (Cavalli-Sforza and Feldman 1981; McCallum et al. 2001; Galef and Laland 2005; Denton et al. 2020). A mechanism likely to be especially relevant in nature is one where the transmission of foraging innovations depends on not only interactions between naïve and innovative individuals, but also with prey. There are many circumstances where naïve individuals need to observe an innovator interact with an object to acquire the relevant skill. For example, capuchin monkeys need to see another monkey interact with a nut to learn how to crack it open (Coelho et al. 2015), and a bird needs to observe another bird interact with a warning-colored prey item to learn avoidance (Hämäläinen et al. 2021). Unlike the models that we explored above, such models of information transfer are not as directly analogous to epidemiological transfer functions.

We explore this scenario by letting *s*(*P*_1_,*P*_2_) include resource density. We do this in two ways (Table 1). First, we extend the mass-action assumption to require three agents: *s*(*P*_1_,*P*_2_) = *sNP*_1_*P*_2_. This means social learning requires that naïve individuals catch a glimpse of an innovator interacting with a prey. Second, we add the rate at which innovators process prey, meaning that we reduce the rate of social learning by prey handling time and encounter rate: 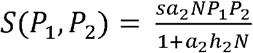. This means social learning requires that a naïve individual observes the entire process of prey capture and handling by an innovator. It is based on the Type II functional response that governs predator foraging. Using bifurcation analyses as above, we examine the effect of these functions on the system across a range of values for *a*_2_, *b*_2_, and *h*_2_ (the spread of mortality-reducing innovations should not depend upon prey density, so we ignored them).

If *s*(*P*_1_,*P*_2_) = *sNP*_1_*P*_2_, then the proportion of innovators in the predator population increases. It also reduces amplitude of the cycles in *P*_2_*/P*_total_, so the population is less variable in the proportion of innovators (Fig. S2). Why does this happen? After a prey population crash, naïve and innovative predators both decline steeply; often, it is difficult for an innovation to spread. However, the *R*_0_ value for information spread is 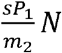. The addition of the *N* term to *R*_0_ causes prey populations to increase the rate of social learning within the predator population as the *prey population* recovers following a crash. This prevents the proportion of innovators from falling as low as when *s*(*P*_1_,*P*_2_) = *sP*_1_*P*_2_. Including *N* in the transmission function also increases the prey abundance at which *R*_0_ is highest in populations of entirely naïve predators, so innovations are more likely to spread at lower predator abundances (i.e. Fig. 4A).

When 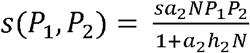, the proportion of innovators in the population decreases rather than increasing (Fig. S3). This pattern is driven by a decrease in the upper bound on the cycles in the proportion of innovators, while the lower bound remains similar to the case where *s*(*P*_1_,*P*_2_) = *sP*_1_*P*_2_. Overall, the amplitude of the cycles decreases. The *R*_0_ for innovation spread when 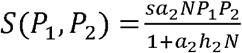 helps us see why: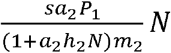. The denominator limits the speed of social information spread with respect to *N*. This causes innovative and naïve predator populations to be more synchronous than when *s*(*P*_1_,*P*_2_) = *sP*_1_*P*_2_, so the proportion of innovators does not grow as extreme. See Fig. S4 for a comparison of the three transmission functions using time series.

### *Influence of* de novo *innovation on Rozensweig-MacArthur systems*

How does the capacity of individuals to learn non-socially impact their dynamics? We examine two cases to address this question. In the first, we let *D*(*P*_1_) = *dP*_1_ and *s*(*P*_1_,*P*_2_) = *sP*_1_*P*_2_. This assumes that the presence of the prey is unnecessary for both *de novo* innovation and social learning. In the second, we let *D*(*P*_1_) = *dP*_1_N and *S*(*P*_1_,*P*_2_) = *sP*_1_*P*_2_*N*. In this case, both mechanisms for acquiring the innovation require a prey item. Again assuming that *D*(*P*_1_) « *s*(*P*_1_,*P*_2_), numerical analyses reveal that regardless of the nature of the innovation (attack, conversion, etc.), systems are dominated by the influence of social learning, and that *de novo* innovation has little impact on their dynamics (Fig. S5). This is intuitive; adding a *de novo* innovation essentially performs a perturbation analysis, nudging the system away from an equilibrium point or cycle. Rosenzweig-MacArthur systems (Edelstein-Keshet 2005) have stable equilibria or limit cycles as attractors, so it is unsurprising that perturbations caused by *de novo* innovation have little impact upon system behavior.

## Discussion

Our models show that behavioral innovations among predators in dynamic populations have the potential to alter the stability of a system, shift the distribution of energy to higher trophic levels, and maintain behavioral variability within predator species. Furthermore, larger innovations tend to be more destabilizing in the more realistic Rosenzeweig-MacArthur system. Although we do not explicitly consider extinction probability as a function of population size, other work shows that periods of low population size can increase the likelihood of this outcome due to stochastic processes (Bartlett 1960; Lande 1993) or Allee effects (Gil et al. 2019; Aubier 2020). Thus, innovations in stable systems could act as catalysts of change that qualitatively alter dynamics by pushing them towards cyclic behavior or local extinction when predators depend on a single species of prey.

Within the Rosenzweig-MacArthur systems, when social learning requires interaction between only naïve and innovative predators, the *R*_0_ for innovation spread 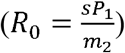 could preclude the innovation from taking hold in small populations that subsist on prey populations near their carrying capacity. Such innovations, like foraging in particular microhabitats or using a particular movement style, might be most likely to spread in established populations. Innovations that spread through a three-way interaction of naïve predator, innovator, and prey, have a maximum *R*_0_ at lower predator abundances (and higher prey abundances) (Fig. S6). These innovations might be most likely to spread in less established predator populations – for example, among introduced species (i.e., a species starting at low abundance) that have yet to become locally invasive. In newly colonized pine forests, black rats *Rattus rattus* learn to open pinecones by observing conspecifics (Aisner and Terkel 1992). Conversely, marginalized native species responding to an invader might be likely to learn to forage on the invader via the same three-way mechanism. This appears to be occurring when some Australian predators encounter cane toads (Beckmann and Shine 2011; Parrott et al. 2019). Innovations that reduce mortality independently of prey can spread more easily than other types of innovations, and so their spread might not be reliably correlated with population size or mode of information transmission. Once an innovation is established in a predator population, the temporal pattern of innovation prevalence is also somewhat sensitive to the type of transmission function (Fig. 4; Figs. S2-S4).

In a spatially structured system, even a single predator and prey might persist if social learning is heterogeneous across the landscape, as populations might cycle asynchronously with the temporal and spatial spread of the innovation (a phenomenon seen in other, non-behaviorally labile systems, and termed the “inflationary effect”; Roy et al. 2005; Kortessis et al. 2020). Thorogood (2018) examined social learning about warning-signaling prey among predators in a metapopulation. The target of their analysis was evolution, rather than population abundances, but their work highlights the importance of considering spatial structure. Another potentially interesting factor we do not explicitly consider is the possibility of stochastic extinction of the innovation (even if the predator species persists). This might be quite important if the innovation is discovered very rarely relative to demographic rates. Stochastic extinction is a realistic component of many aspects of cultural evolution, where cultural entities are often envisioned as having their own evolutionary histories of origination and extinction (Dawkins 1976; Cavalli-Sforza and Feldman 1981; Pagel 2009). It also appears to be quite common with innovation in animal societies. For example, predatory behavior appears to have been acquired several times by Great Tits (*Parus major*), with isolated reports of attacks on other birds in the literature (Saunders 1889; Caris 1958; Barnes 1975) and in the popular news (Jokinen 2013). A population of Great Tits in Hungary apparently also learned to find and kill bats as they emerged from their hibernation caves (Estók et al. 2010). However, these behaviors are not commonly observed in the species across its very large geographic range. Thus, they may represent an instance of spontaneous emergence and transient local persistence of innovations (although more research is needed to determine any genetic component). The probability of extinction and re-emergence of an innovation would add an additional layer of nuance to our predictions for behavioral variability. Generally speaking, it would be of great interest to expand the complexity of the systems in which innovation transmission is modeled.

There must be some limits placed on the use of social information for its rate of transmission to be held in check. Why would an animal *not* adopt an advantageous novel behavior from another member of its species? All learning can be costly due to time, risk, or energy expended in acquiring information (Sherratt 2011; Abbott and Sherratt 2013; Kikuchi and Sherratt 2015). Furthermore, in a randomly varying environment, associations learned at one moment in time are only adaptive if they indicate the best action at subsequent times (Dunlap and Stephens 2009), so investments in learning may not always be recouped. Other potential costs to social learning include increased competition from conspecifics (Seppänen et al. 2007), as well as increased chances of acquiring pathogens (Cantor et al. 2021).Indeed, the evolution of sociality and pathogen virulence may co-determine animal social networks (Prado et al. 2009).

Social information is just one type of information available to organisms. Organisms also obtain information through genetic biases, early developmental effects, and personal experience (Laland 2004; Dall et al. 2005). A critical alternative to social information is personal information. Which one an animal uses can be context-dependent. When bats have unreliable personal information about prey cues, they are more likely to exploit social information (Jones et al. 2013; Dunlap et al. 2016). Some recent models examine the evolution of personal versus social information use (Wakano and Aoki 2007; Borofsky and Feldman 2022). Borofsky and Feldman (2022) studied the population dynamics of two prey species attacked by predators of constant population size. In their model, a predator’s tendency to use social information coevolved with its tendency to exhibit conformity or anticonformity, both of which aided predators in making adaptive foraging decisions. Resource competition favored anticonformity, maintaining variability in predator behavior (Borofsky and Feldman 2022).

Although models of social versus personal information ask an evolutionary question, they also have implications for ecology. Adaptive foraging can stabilize the dynamics of ecological networks (Valdovinos et al. 2010). A general question is how social (and personal) information affect the dynamics of multispecies communities. The results of the present study make it tempting to hypothesize that highly innovative species may depend upon continuous innovation to exploit new populations of prey if their innovations drive local resources extinct, or to abundances so low that predator populations cannot sustain themselves. Among birds, innovative species tend to have lower extinction risk and more stable or increasing populations than less innovative species (Ducatez et al. 2020). However, this is attributed to innovators responding less aversely to habitat destruction, so the results of (Ducatez et al. 2020) do not speak directly to our study of predator-prey relationships. A promising future direction is examining the ecological dynamics predator-prey systems where information use is evolutionarily labile.

Although we focused on innovations among predators, prey may reciprocally change in response to predator innovations, rendering predator innovations less advantageous. Prey responses are well-documented in the context of predator evolutionary innovations (e.g. Hanifin et al. 2008), but could occur in response to predator behavioral innovations too (Whitehead et al. 2019; Cantor et al. 2021).Furthermore, countermeasures by prey are socially transmitted, such as alarm calling (Magrath et al. 2015) and habitat use (Fortin et al. 2005). Cultural arms races between predators and prey have the potential to create a constantly shifting landscape of fear over which predators and prey contest their existence (Brown et al. 1999; Laundré et al. 2001).

One conclusion of this study is that social learning can maintain behavioral variability in certain predator-prey systems. Another is that social learning can modify the trophic distribution of predator and prey across time by creating population cycles. In general, emerging theory suggests that social information can have significant impacts on population persistence (Schmidt et al. 2015; Schmidt 2017), competitive outcomes (Gilpin et al. 2016; Gil et al. 2018, 2019; Wakano et al. 2018), and predatory behavior (Borofsky and Feldman 2022). Exploring further scenarios such as those we have discussed will help us broadly understand the role information plays in ecological processes and maintaining phenotypic diversity within populations.

## Supporting information

All supplements

## Acknowledgments

We would like to thank members of the Holt lab group, Isabel Damas-Moreira, Marcus Feldman, Klaus Reinhold, and Rose Thorogood for helpful comments and discussion.

## Funding

German Research Foundation (DFG): SFB TRR 212 (NC3) – 316099922 (DWK)

United States Department of Agriculture (USDA): USDA-NSF-NIH Ecology and Evolution of Infectious Diseases program – 2017-67013-26870 (MWS)

University of Florida Foundation (MWS)

We acknowledge support for the publication costs by the Open Access Publication Fund of Bielefeld University.

## Author contributions

Conceptualization: DWK, MWS

Methodology: DWK, MWS

Investigation: DWK, MWS

Visualization: DWK

Writing—original draft: DWK

Writing—review & editing: DWK, MWS

## Data and materials availability

All code is available in the supplementary materials.

---

**Supplementary Figure 1.**
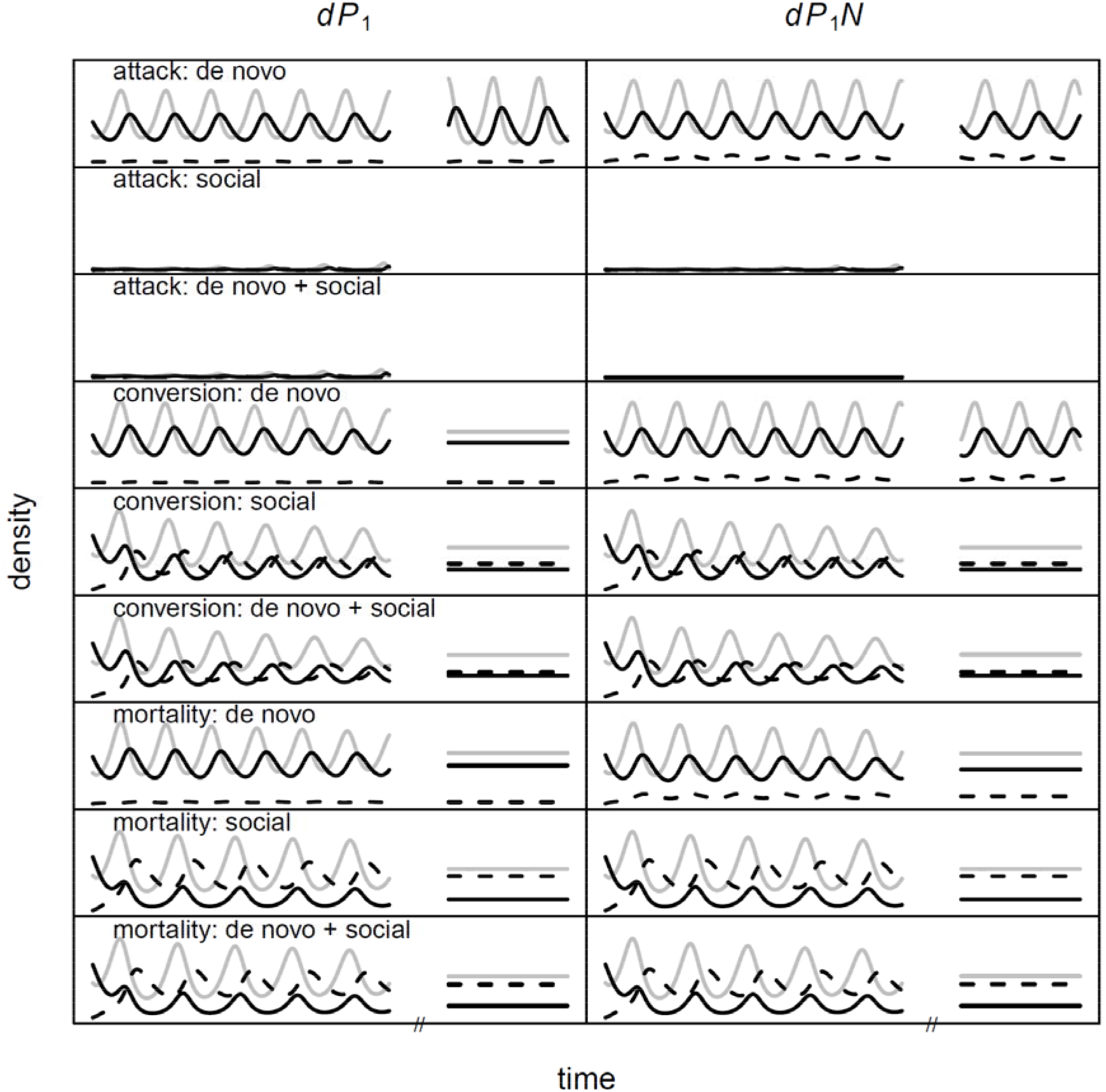
Time series comparison of the effects of *de novo* innovation versus acquisition of innovation via social learning on Lotka-Volterra predator-prey population dynamics. In all cases, when both were possible, the behavior of the system more closely resembles that with social learning alone, rather than *de novo* innovation alone. When not a trait subject to innovation (so values were equal between naïve and innovative predators), *a* = 0.15, *b* = 0.4, *m* = 0.2, *s* = 0.16, *d* = 0.01, *r* = 0.4. For innovations, *a*_2_ = 0.2, *b*_2_ = 0.7, and *m*_2_ = 0.12.

**Supplementary Figure 2.**
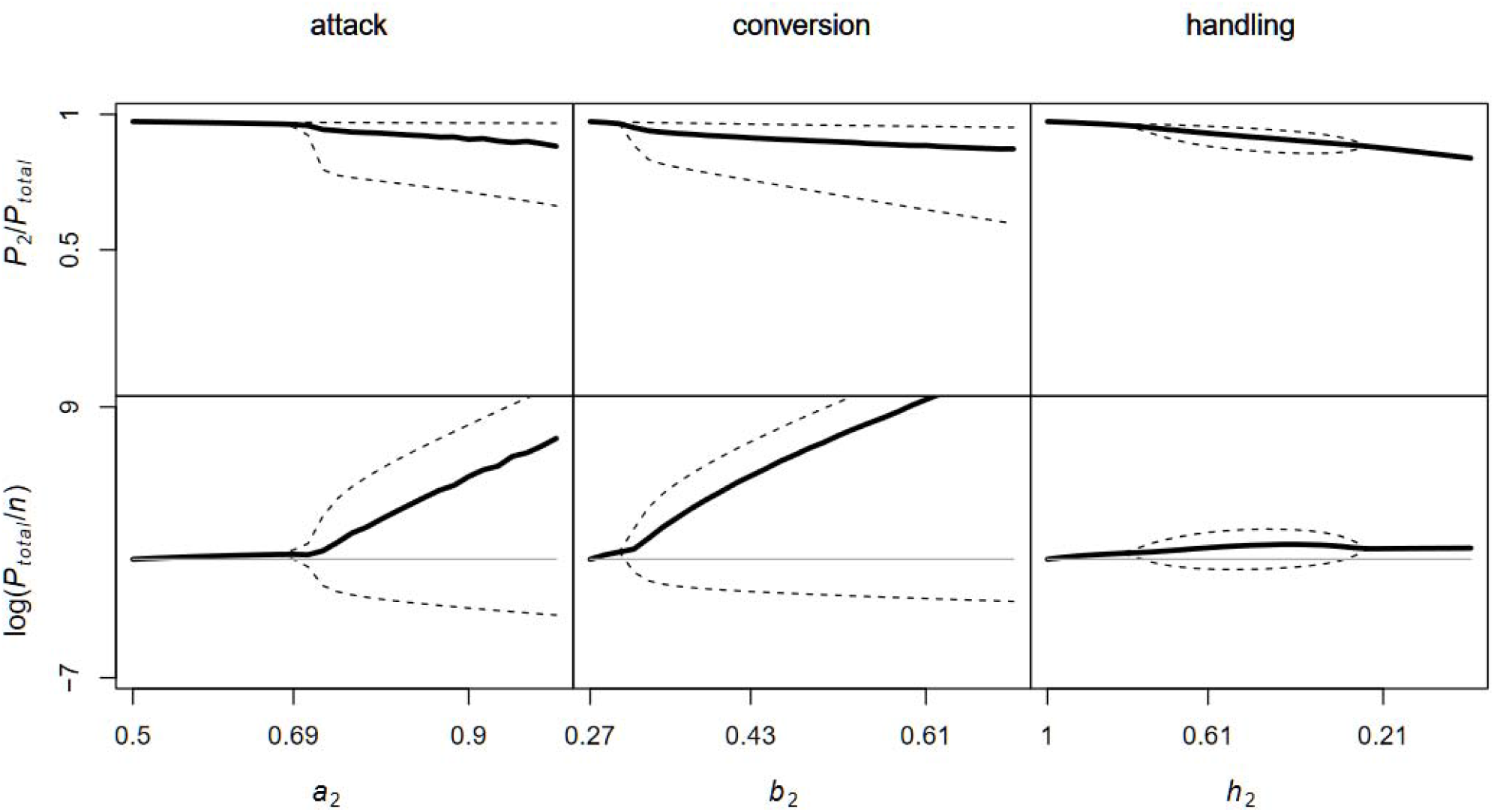
Bifurcation analyses of the system in response to each kind of innovation with the *sNP*_1_*P*_2_ transmission function. Except for the varying parameter on the abscissa, naïve and innovative predators shared the same parameter values: *a* = 0.5, *b* = 0.5, *m* = 0.25, *h* = 0.85, *r* = 2, *s =* 0.2, *k* = 10.

**Supplementary Figure 3.**
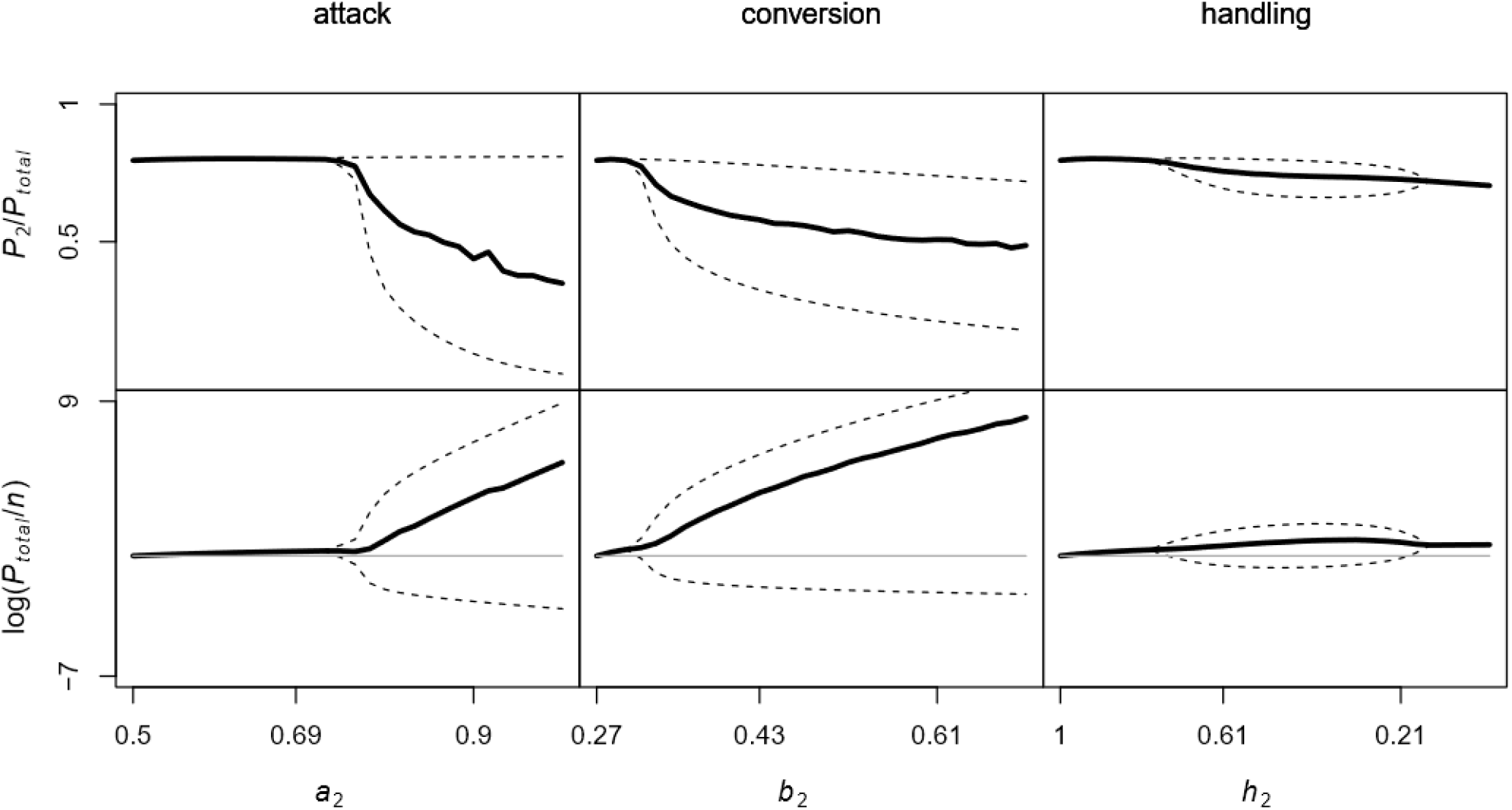
Bifurcation analyses of the system in response to each kind of innovation with the 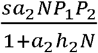 transmission function. Except for the varying parameter on the abscissa, naïve and innovative predators shared the same parameter values: *a* = 0.5, *b* = 0.5, *m* = 0.25, *h* = 0.85, *r* = 2, *s =* 0.2, *k* = 10.

**Supplementary Figure 4.**
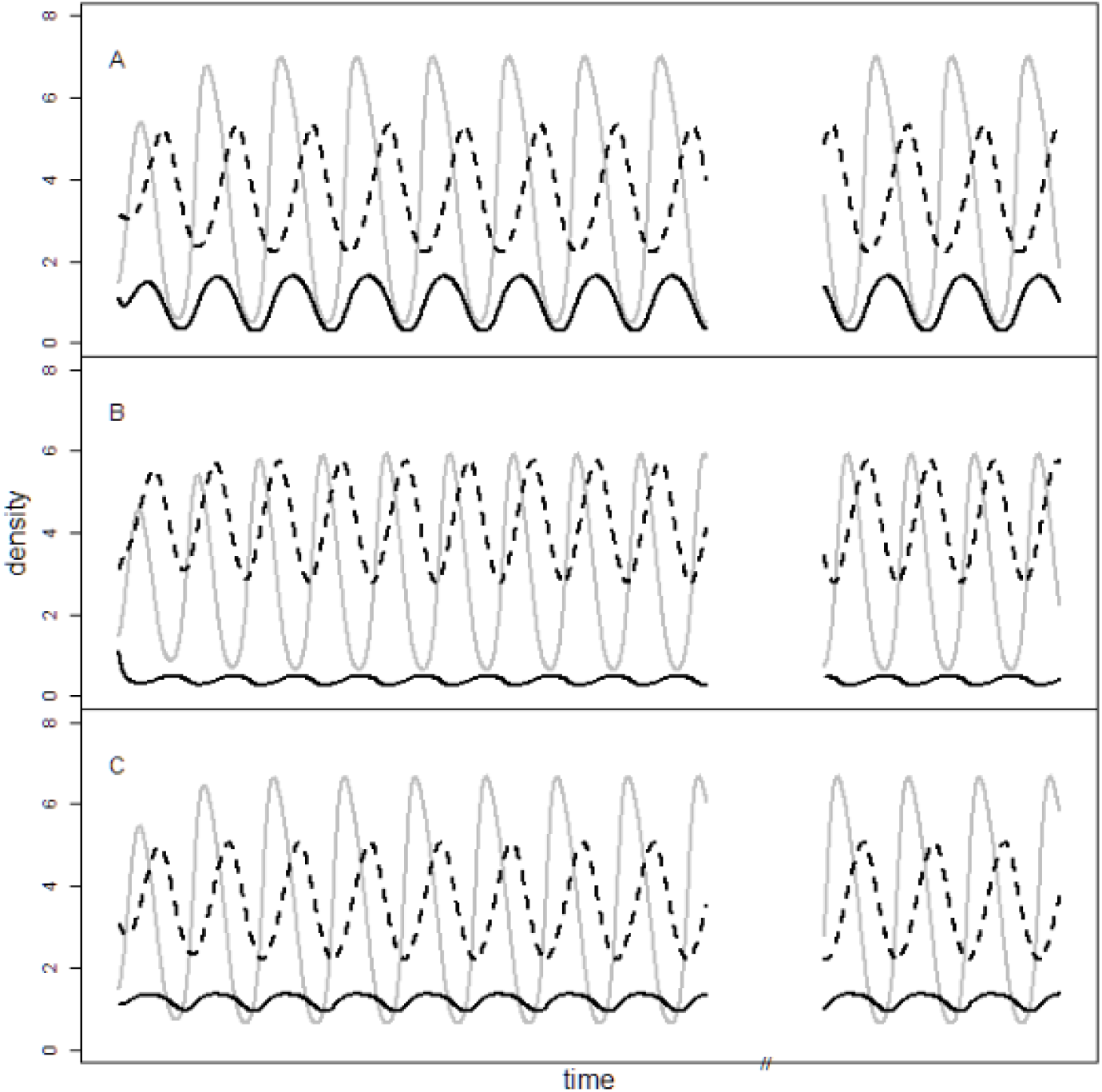
Time series showing the population dynamics of prey (gray), naïve predators (black), and innovative predators (dashed) in Rosenzweig-MacArthur system. A. The *sP*_1_*P*_2_ transmission function. Here, the peaks of naïve and innovative predator cycling are almost completely out of phase, creating large fluctuations in the proportion of innovators in the predator population. B. The *sNP*_1_*P*_2_ transmission function. With this function, following a predator population crash, the innovative predator population recovers *first*, followed by the naïve predator population. This means a larger proportion of the predator population remains in the innovator class. C. The 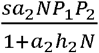 transmission function. In this case, the saturating transmission of the innovation tends to synchronize the cycles in populations of naïve and innovative predators. This keeps the proportion of innovators from growing extremely high. Parameter values: *a* = 0.5, *b* = 0.5, *m* = 0.25, *h*_1_ = 1, *h*_2_ = 0.5, *r* = 2, *s =* 0.2, *k* = 10.

**Supplementary Figure 5.**
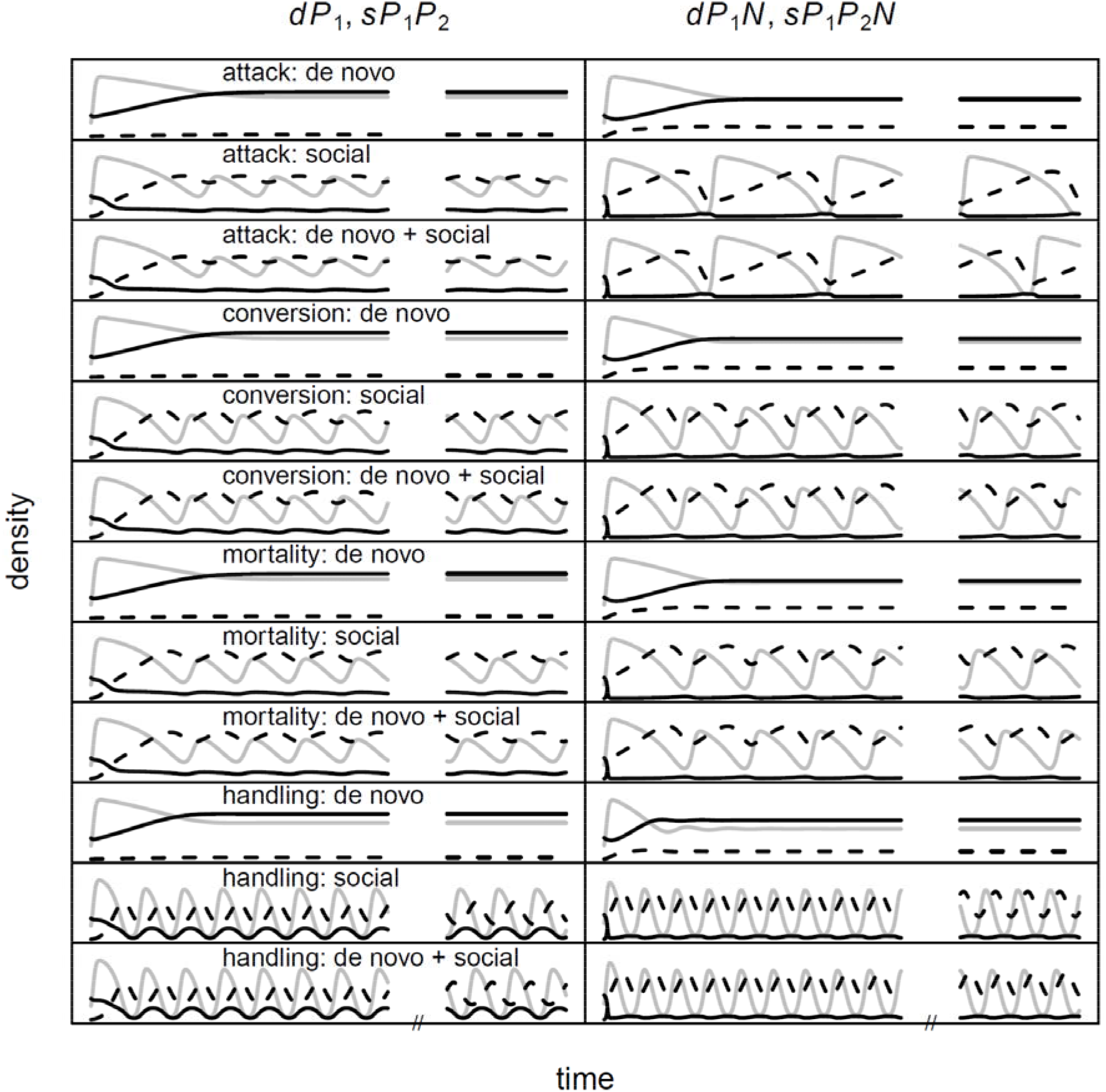
Time series comparison of the effects of *de novo* innovation versus acquisition of innovation via social learning on Rosenzweig-MacArthur predator-prey population dynamics. In all cases, when both were possible, the behavior of the system more closely resembles that with social learning alone, rather than *de novo* innovation alone. When not a trait subject to innovation (so values were equal between naïve and innovative predators), *a* = 0.5, *b* = 0.27, *m* = 0.2, *h* = 1, *s* = 0.2, *d* = 0.01, *r* = 2, *k* = 10. For innovations, *a*_2_ = 0.72, *b*_2_ = 0.31, *m*_2_ = 0.175, *h*_2_ = 0.5.

**Supplementary Figure 6.**
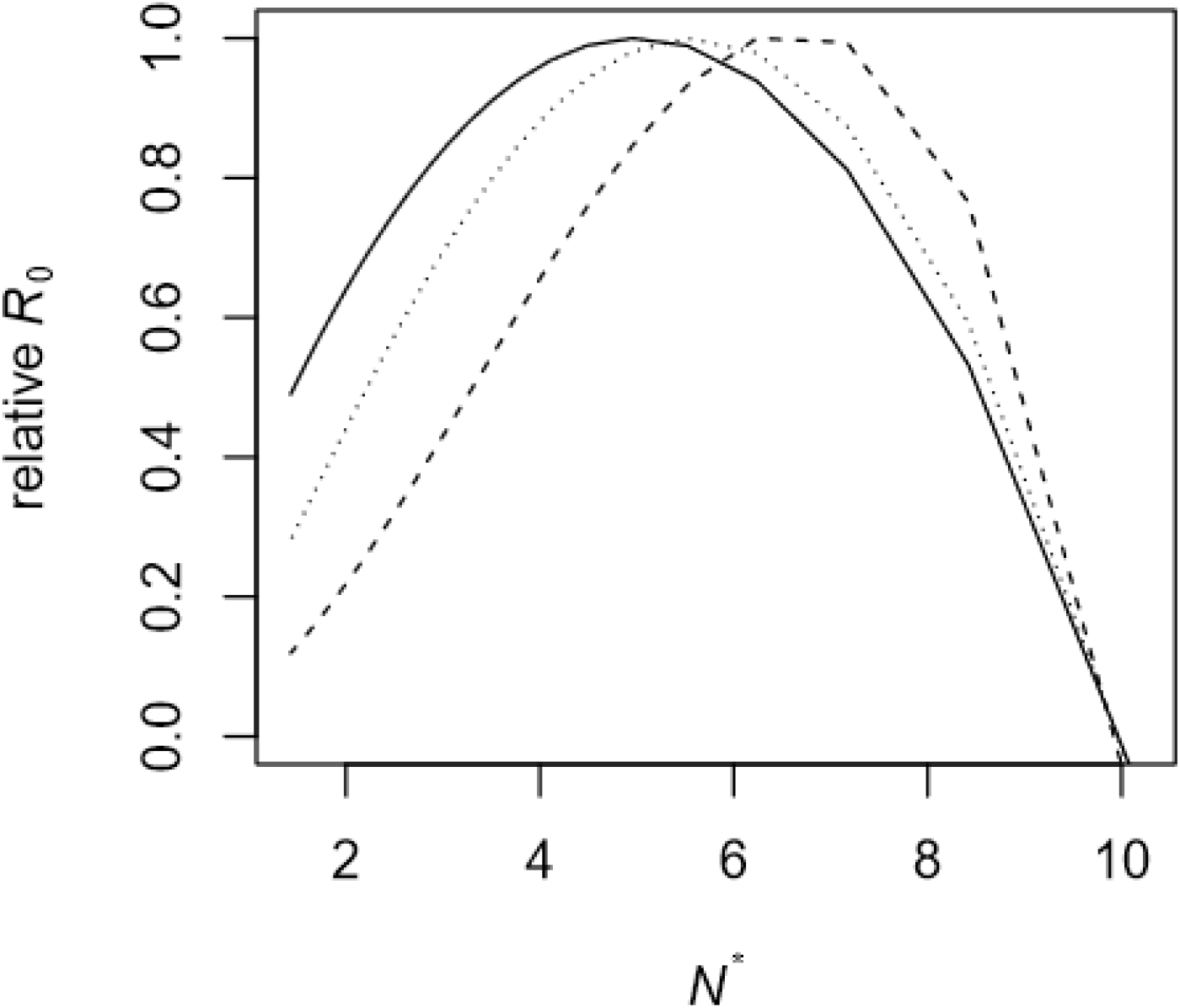
Relative *R*_0_ (i.e. scaled to 1) under the *sP*_1_ *P*_2_ transmission function (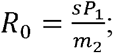; solid), *sNP*_1_ *P*_2_ transmission function (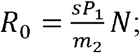; dashed), and 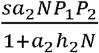 transmission function (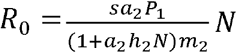; dotted). Innovations that spread via the latter two transmission functions will be more likely to invade at higher prey population abundances (and lower predator population abundances).

