## Supplementary material for "Socially transmitted innovations in dynamic predator-prey systems": All supplements: Fig2.R

```
library(deSolve)

#### Predator-prey dynamics when predators use social learning ####

## Figure 2 - timeseries from Lotka-Volterra model where populations are dynamic
##            and there is an innovation in attack rate spreading among predators.

#### Lotka-Volterra model ####
MODEL <- function(time, state, parameters) {
  with(as.list(c(state, parameters)), {       
    dn <-  n*(r-a1*p1-a2*p2)
    dp1 <-  a1*b1*p1*n+a2*b2*p2*n-s*p1*p2-m1*p1
    dp2 <-  s*p1*p2-m2*p2
    return(list(c(dn, dp1, dp2)))
  })
}

## Specify parameter values
Parms <- list(r = 0.4, 
              a1 = 0.15,
              a2 = 0.15, 
              b1 = 0.4, 
              b2 = 0.7,
              s = 0.16,
              m1 = 0.2,
              m2 = 0.2)

## Specify Initial Conditions 
## Here I am starting the system a little off equilibrium (n_star, p1_star, p2_star) 
## in case the equilibrium point is unstable.
n_star = with(Parms, m2*(a2*m1 - a1*m2 + r*s)/(a1*a2*(b1 - b2)*m2 + a2*b2*r*s))
p1_star = with(Parms, m2/s)
p2_star = with(Parms, (r*s-a1*m2)/(a2*s))
init = c(n = n_star, p1 = p1_star+0.1, p2 = p2_star+0.1)

sim.time <- seq(0, 4000, by = 0.1)

## Feed model, initial conditions and parameter values into solver
out <- ode(y=init, times=sim.time, func=MODEL, parms=Parms)
out = as.data.frame(out)

maa <- dim(out)[1]
pdf('Fig1.pdf',4,3)
par(mfrow = c(2,1), mar = c(0,0,0,0), oma = c(2,2,0.5,0.5))
plot(out$time[0:1500], out$n[0:1500], xlim = c(0,240),
     pch=16,  type = 'l', lwd = 2, col = 'gray',
     ylim=c(0, max(out$n, out$p1, out$p2)+1),
     ylab="Abundance", xlab="Time", yaxt = 'n', xaxt = 'n')
lines(out$time[0:1500], out$p1[0:1500], lwd = 2, lty = 1)
lines(out$time[0:1500], out$p2[0:1500], lwd = 2, lty = 2)
lines(out$time[1800:2400], out$n[(maa-600):maa], lwd = 2, col = 'gray',)
lines(out$time[1800:2400], out$p1[(maa-600):maa], lwd = 2, lty = 1)
lines(out$time[1800:2400], out$p2[(maa-600):maa], lwd = 2, lty = 2)
text(240/30,(max(out$n, out$p1, out$p2)+0.5)/20*19 ,'A', cex = 1.2)

#panel B

Parms <- list(r = 0.4, 
              a1 = 0.15,
              a2 = 0.15, 
              b1 = 0.4, 
              b2 = 0.7,
              s = 0.25,
              m1 = 0.2,
              m2 = 0.2)

## Specify Initial Conditions 
## Here I am starting the system a little off equilibrium (n_star, p1_star, p2_star) 
##in case the equilibrium point is unstable
n_star = with(Parms, m2*(a2*m1 - a1*m2 + r*s)/(a1*a2*(b1 - b2)*m2 + a2*b2*r*s))
p1_star = with(Parms, m2/s)
p2_star = with(Parms, (r*s-a1*m2)/(a2*s))
init = c(n = n_star, p1 = p1_star+0.1, p2 = p2_star+0.1)

sim.time <- seq(0, 4000, by = 0.1)

out <- ode(y=init, times=sim.time, func=MODEL, parms=Parms)
out = as.data.frame(out)

## plot this model run

plot(out$time[0:1500], out$n[0:1500], xlim = c(0,240),
     pch=16,  type = 'l', lwd = 2, col = 'gray',
     ylim=c(0, max(out$n, out$p1, out$p2)+1),
     ylab="Abundance", xlab="Time", yaxt = 'n', xaxt = 'n')
lines(out$time[0:1500], out$p1[0:1500], lwd = 2, lty = 1)
lines(out$time[0:1500], out$p2[0:1500], lwd = 2, lty = 2)
lines(out$time[1800:2400], out$n[(maa-600):maa], lwd = 2, col = 'gray',)
lines(out$time[1800:2400], out$p1[(maa-600):maa], lwd = 2, lty = 1)
lines(out$time[1800:2400], out$p2[(maa-600):maa], lwd = 2, lty = 2)
text(165,-.3,'//', cex = 1, xpd = NA)
text(240/30,(max(out$n, out$p1, out$p2)+0.5)/20*19 ,'B', cex = 1.2)
mtext('time', side = 1, line = 0.4, outer = T)
mtext('density', side = 2, line = 0.4, outer = T)
dev.off()
```
