## Supplementary material for "Socially transmitted innovations in dynamic predator-prey systems": All supplements: Fig3.R

```
## Figure 2 - equilibrium proportion innovators in the SI model without vital rates
##            compared to the dynamic model where populations vary

## Vary r so that prey growth rate changes
varying.parm <- seq(0.7, 6, length.out = 30)
vparm = 8 #must set this to corresponding parameter in Parms

## Specify parameter values (focal parameter takes the first value in varying.parm)
Parms <- list(a1 = 0.167, #default 0.167 for b1 < b2, m1 > m2
              a2 = 0.167,
              b1 = 0.5, #default 0.5 for b1 < b2, m1 > m2
              b2 = 0.55, 
              m1 = 0.5,#default 0.55 for b1 < b2
              m2 = 0.5, #default 0.5 for m1 > m2
              s = 0.16, #default 0.2 for b1 < b2, 0.16 for m1 > m2
              r = varying.parm[1])   #default 2 for b1 < b2, m1 > m2  

## Find equilibrium conditions for dynamic model
beg <- function(parama){
  a1 = parama[[1]]; a2 = parama[[2]]; b1 = parama[[3]]; b2 = parama[[4]]
  m1 = parama[[5]]; m2 = parama[[6]]; s = parama[[7]]; r = parama[[8]]
  n_star = m1*(a2*m1-a1*m2+r*s)/(a1*a2*m2*(b1-b2)+a2*b2*r*s)
  p1_star = m2/s
  p2_star = (r*s - a1*m2)/(a2*s)
  init = c(n = n_star, p1 = p1_star, p2 = p2_star)
  init
}

#plot equilibrium population size as a function of parameter value
eq <- beg(Parms)
for(i in 2:length(varying.parm)){
  Parms[[vparm]] <- varying.parm[i] 
  eq <- rbind(eq,beg(Parms)) 
}
Parms[[vparm]] <- varying.parm[1] 

yup <- cbind(varying.parm,eq)
pdf('figure2.pdf', 3,3)
par(mfrow = c(1,1), mar = c(4,5,1,1))
plot(yup[,1], yup[,4]/(yup[,3]+yup[,4]), type = 'l', ylim = c(0,1),
     xlab = expression(paste(italic(r))), ylab = 'proportion innovators')
#equilibrium from SI model (see Mathematica notebook for derivation)
abline(h = (Parms[[6]]/Parms[[7]])/(1+Parms[[6]]/Parms[[7]]), lty = 2) 
dev.off()
#legend('right', c('SI','LV P-P'),lty=c(2,1))
```
