## Supplementary material for "Socially transmitted innovations in dynamic predator-prey systems": All supplements: Figure4_SimpleNullclines.R

```
library(deSolve)

## Figure 3 - Panel A: A Rosenzweig-MacArthur system at stable equilibrium
##            Panel B: A similar system cycling about an unstable equilibrium

x <- seq(0,11,0.1)
a = 0.52
b = 0.24
m = 0.2
h = 1
r = 1
k = 10

prey <- function(n,k,r,h,a) ((k-n)*r*(1+a*h*n))/(a*k)
pred <- function(a,b,h,m) m/(a*(b-h*m))
eqpred <- function(n,a,b,h,m,r,k) b*r*(a*b*k-m-a*h*k*m)/(a^2 * k * (b - h*m)^2)

pdf('fig3.pdf', 3.2, 1.8)
par(mfrow = c(1,2), mar = c(0,0,0,0), oma = c(1.5,1.5,0.3,0.3))
plot(x,prey(x,k,r,h,a), type = 'l', lwd = 2.5, xaxt = 'n', yaxt = 'n',
     ylim = c(0,5),
     xaxs = 'i', yaxs = 'i', col = 'gray')
abline(v = pred(a,b,h,m), lwd = 2.5, lty = 1)
points(pred(a,b,h,m), eqpred(n,a,b,h,m,r,k), pch = 16, cex = 1.5)
mtext('predator', side = 2, line = 0.5, outer = T)
text(1,4.5,'A',cex = 1.2)

#cyclical plot
#a <- 0.5
b <- 0.305
#h <- 1

plot(x,prey(x,k,r,h,a), type = 'l', lwd = 2.5, xaxt = 'n', yaxt = 'n',
     ylim = c(0,5),xaxs = 'i', yaxs = 'i', col = 'gray')
abline(v = pred(a,b,h,m), lwd = 2.5, lty = 1)
points(pred(a,b,h,m), eqpred(n,a,b,h,m,r,k), cex = 1.5)

t2 <- function(a, h, p, n) a*p*n/(1 + a*h*n) 
cycmod <- function(time, state, parameters) {
  with(as.list(c(state, parameters)), {   
    dn <- n*r*(k-n)/k - t2(a,h,p,n) 
    dp <- b*t2(a,h,p,n) - m*p 
    return(list(c(dn, dp)))
  })
}
Parms <- list(a,b,h,m,r,k)
sim.time <- seq(0, 1000, by = 0.1)
out <- ode(y = c(n = 1,p = 1), times = sim.time, func=cycmod, parms=Parms)
ma <- dim(out)[1]
lines(out[(ma-680):(ma-320),2],out[(ma-680):(ma-320),3], lty = 3, lwd = 1.5)
arrows(out[ma-680,2], out[ma-680,3], out[ma-679,2], out[ma-679,3], length = 0.09)
arrows(out[ma-540,2], out[ma-540,3], out[ma-539,2], out[ma-539,3], length = 0.09)
arrows(out[ma-450,2], out[ma-450,3], out[ma-449,2], out[ma-449,3], length = 0.09)
arrows(out[ma-370,2], out[ma-370,3], out[ma-369,2], out[ma-369,3], length = 0.09)
text(1,4.5,'B', cex = 1.2)
mtext('prey', side = 1, line = 0.4, outer = T)
dev.off()
```
