## Supplementary material for "Socially transmitted innovations in dynamic predator-prey systems": All supplements: Figure5_SuppFigs2and3_pretty.R

```
library(deSolve)

t2 <- function(a, h, p, n) a*p*n/(1 + a*h*n) #type 2 functional response

m3 <- function(x){ #finds max, min, and mean for time series data
  a <- max(x)
  b <- min(x)
  z <- mean(x)
  c(a,b,z)
}

## Figure 4 - Bifurcation diagrams for the mass action s*P1*P2 transmission
##            function. 
## Supplementary Figures 1 & 2 - Exploration of two alternative transmission 
##                               functions with bifurcation diagrams. You will
##                               need to change the transmission functions
##                               accordingly, and comment out the mortality
##                               innovation in the variable parameters.

### The Models ###
## to be solved using dsolve ##

#mass action social learning depends on P1 and P2 and optionally N
RosenzweigMacArthur <- function(time, state, parameters) {
  with(as.list(c(state, parameters)), {   
    dn <- n*r*(k-n)/k - t2(a1,h1,p1,n) - t2(a2,h2,p2,n)
    dp1 <- b1*t2(a1,h1,p1,n) + b2*t2(a2,h2,p2,n) - s*p1*p2 - m1*p1 #change to appropriate transmission function
    dp2 <- s*p1*p2  - m2*p2 #change to appropriate transmission function
    return(list(c(dn, dp1, dp2)))
  })
}

## Specify Initial Conditions (for all R-M models)
## Finds equilibrium for the n, p1 alone (basic consumer-resource) model.
## Also gives the equilibrium point for the model
## when all individuals are naive and there
## is no social learning, a stable point of comparison
## under the default parameter values.
beg <- function(parama){
  a1 = parama[[1]]; a2 = parama[[2]]; b1 = parama[[3]]; b2 = parama[[4]];
  m1 = parama[[5]]; m2 = parama[[6]]; h1 = parama[[7]]; h2 = parama[[8]];
  s = parama[[9]]; r = parama[[10]]; k = parama[[11]]; 
  n_star = m1/(a1*(b1-h1*m1))
  p1_star = b1*r*(a1*b1*k-m1-a1*h1*k*m1)/(a1^2 * k*(b1-h1*m1)^2)
  p2_star = 0
  init = c(n = n_star, p1 = p1_star, p2 = p2_star)
  init
}

## What values of the parameter we want to examine
## should we plot in the bifurcation diagram?

#Give values of varying parameters as a list to loop through
#You will need to change the default parameters to e.g. a1 = 0.5, a2 = 0.55
#if evaluating sensitivity to a non-innovation trait like
#social learning speed or prey growth rate.
variableparameters <- list(seq(0.5,1,length.out=30),
                           seq(0.27,0.7,length.out=30),
                           seq(0.05,0.2,length.out=30), 
                           #don't include mortality for transmission functions involving N;
                           #this doesn't make sense in any case I can think of
                           seq(0.05,1,length.out=30)
)

vp <- c(2,
        4,
        6,
        8) #index of which parameters are varying in Parms - ALSO UPDATE (6 = m2)

ts <- 10000  #total time
asst <- 5000 #burn-in time


#pdf(paste("differentinnovations_bifurc.varying_3way_",n[vparm],'.a1_',v[1],'.a2_',v[2],
#          '.b1_',v[3],'.b2_',v[4],'.m1_',v[5],'.m2_',v[6],'.s_',v[7],'.r_',v[8],
#          ".pdf",sep=''),6,4)
if(length(vp) > 3){
  par(mfcol = c(2,4), oma = c(5,5,4,1), mar = c(0,0,0,0))
} else{
  par(mfcol = c(2,3), oma = c(5,5,4,1), mar = c(0,0,0,0))
}


for(j in 1:length(variableparameters)){
  ## First, specify default parameter values
  Parms <- list(a1 = 0.5, 
                a2 = 0.5,
                b1 = 0.27,  
                b2 = 0.27, 
                m1 = 0.2, 
                m2 = 0.2, 
                h1 = 1, 
                h2 = 1,
                s = 0.2, #default 0.2 
                r = 2,    #default 2 for b1 < b2, m1 > m2
                k = 10   #default 10
  )   
  varying.parm <- variableparameters[[j]]
  vparm <- vp[j] #tells us which index in Parms to vary
  Parms[vparm] <- varying.parm[1]
  
  ## Specify simulation time
  sim.time <- seq(0, ts, by = 0.1)
  
  ## Feed models, initial conditions and parameter values into solver
  out_SLT2_2way <- ode(y=beg(Parms)+0.1, times=sim.time, 
                       func=RosenzweigMacArthur, parms=Parms)
  
  ## Begin bifurcation analyses to see where stability yields to cyclic dynamics
  
  runs <- (nrow(out_SLT2_2way) - asst):nrow(out_SLT2_2way) #time interval to average over
  lastbunch <- data.frame(out_SLT2_2way[runs,])
  lastbunch$ptot <- rowSums(lastbunch[,3:4], na.rm = T)
  lastbunch$p2tot <- lastbunch[,4]/lastbunch$ptot
  lastbunch$pn <- lastbunch$ptot/lastbunch[,2]
  colnames(lastbunch) <- c('time','n','p1','p2','ptot','p2tot','pn')
  
  bifurc_ptot <- c(Parms[[vparm]],m3(lastbunch$ptot))
  bifurc_p2tot <-  c(Parms[[vparm]],m3(lastbunch$p2tot))
  bifurc_pn <-  c(Parms[[vparm]],m3(lastbunch$pn))
  eq <- beg(Parms) #calculate the equilibrium solution for given parameter values
  
  ### For loop
  for(i in 2:length(varying.parm)){
    Parms[[vparm]] <- varying.parm[i] #auto-update the focal parameter
    
    out_SLT2_2way <- ode(y=beg(Parms)+0.01, times=sim.time, 
                         func=RosenzweigMacArthur, parms=Parms)
    lastbunch <- data.frame(out_SLT2_2way[runs,])
    lastbunch$ptot <- rowSums(lastbunch[,3:4], na.rm = T)
    lastbunch$p2tot <- lastbunch[,4]/lastbunch$ptot
    lastbunch$pn <- lastbunch$ptot/lastbunch[,2]
    colnames(lastbunch) <- c('time','n','p1','p2','ptot','p2tot','pn')
    
    bifurc_ptot <- rbind(bifurc_ptot,c(Parms[[vparm]],m3(lastbunch$ptot)))
    bifurc_p2tot <- rbind(bifurc_p2tot,c(Parms[[vparm]],m3(lastbunch$p2tot)))
    bifurc_pn <- rbind(bifurc_pn,c(Parms[[vparm]],m3(lastbunch$pn)))
    eq <- rbind(eq,beg(Parms)) #calculate the equilibrium solution for given parameter values
  }
  
  bifurc_ptot <- cbind(bifurc_ptot,eq[,2])
  bifurc_p2tot <- cbind(bifurc_p2tot, NA)
  bifurc_pn  <- cbind(bifurc_pn, eq[,2]/eq[,1])
  cats <- rep(c('ptot','p2tot','pn'), each = nrow(bifurc_ptot))
  op <- cbind(data.frame(rbind(bifurc_ptot,bifurc_p2tot,bifurc_pn)),cats)
  n <- names(Parms)
  nombres <- c(n[vparm],
               'max_2way','min_2way','mean_2way',
               'eq','var')
  colnames(op) <- nombres
  row.names(op) <- NULL
  
  #option to record data
  #v <- unlist(Parms)
  #fname <- paste('RosenzweigMacArthur_sa2NP1P2_1plusa2h2N_varying_',n[vparm],'.a1_',v[1],'.a2_',v[2],
  #               '.b1_',v[3],'.b2_',v[4],'.m1_',v[5],'.m2_',v[6],'.h1',v[7],'.h2',v[8],'.s_',v[9],'.r_',v[10],'.k',v[11],
  #               '.csv', sep = '')
  #write.csv(op,fname,row.names=F)
  
  ### Graphics ###
  ### If necessary, load data first instead of running long loop ###
  ### Really a separate script, but appended here for working convenience ###
  
  ### bifurcation diagram for p1, p2
  
  i = j #atavism
  
  #panel y-axis labels
  
  y1 <- ifelse(i == 1, expression(paste(italic(P[2]),'/',italic(P[total]),sep = ' ')),'')
  y2 <- ifelse(i == 1, expression(italic(P[total])),'')
  y3 <- ifelse(i == 1, expression(paste('log(',italic(P[total]),'/',italic(N),")",
                                        sep = '')),'')
  
  if(i < 3){
    #first panel down
    
    plot(bifurc_p2tot[,1], bifurc_p2tot[,2], type="l", lty = 2,
         ylab=y1, yaxt = 'n',
         ylim=c(0, 1),
         xlab="", xaxt = "n")
    if(i == 1)  {
      axis(2, at = c(0.5,1), outer = T, 
           labels = as.character(c(0.5, 1)))
      mtext(y1,2, line = 2.5, outer = T, adj = 0.9)
    }
    
    lines(bifurc_p2tot[,1], bifurc_p2tot[,4], lwd=3)
    lines(bifurc_p2tot[,1], bifurc_p2tot[,3], lty = 2) 
    
    ## Second panel down
    pnmax <- 10000
    plot(bifurc_pn[,1], log(bifurc_pn[,2]),  
         ylab=y2, yaxt = 'n', type = 'l' , lty = 2,
         xlab="", xaxt = 'n', ylim=log(c(0.001, pnmax + 0.05)))
    if(i == 1){
      axis(2, at = log(c(0.001,pnmax)), outer = T,
           labels = as.character(round(log(c(0.001, pnmax)))))
      mtext(y3,2, line = 2.5, outer = T, adj = 0.12)
    }
    lines(bifurc_pn[,1], log(bifurc_pn[,4]), lwd=3)
    lines(bifurc_pn[,1], log(bifurc_pn[,3]), lty = 2)
    lines(bifurc_pn[,1], log(bifurc_pn[,5]), col = 'darkgray')
    
    axis(1, at = varying.parm[c(1,12,24)], outer = T,
         labels = as.character(round(varying.parm[c(1,12,24)],2)))
  }
  else{
    #first panel down
    plot(rev(bifurc_p2tot[,1]), bifurc_p2tot[,2], type="l", lty = 2,
         ylab=y1, yaxt = 'n',
         ylim=c(0, 1),
         xlab="", xaxt = "n")
    lines(rev(bifurc_p2tot[,1]), bifurc_p2tot[,4], lwd=3)
    lines(rev(bifurc_p2tot[,1]), bifurc_p2tot[,3], lty = 2) 
    
    ## Second panel down
    pnmax <- 10000
    plot(rev(bifurc_pn[,1]), log(bifurc_pn[,2]),  
         ylab=y2, yaxt = 'n', type = 'l' , lty = 2,
         xlab="", xaxt = 'n',
         ylim=log(c(0.001, pnmax + 0.05)))
    lines(rev(bifurc_pn[,1]), log(bifurc_pn[,4]), lwd=3)
    lines(rev(bifurc_pn[,1]), log(bifurc_pn[,3]), lty = 2)
    lines(rev(bifurc_pn[,1]), log(bifurc_pn[,5]), col = 'darkgray')
    
    axis(1, at = rev(varying.parm[c(1,12,24)]), outer = T,
         labels = as.character(round(varying.parm[c(6,18,30)],2)))
  }
  
  mtext(bquote(italic(.(substr(names(Parms)[vparm],1,1))[2])), 
        at = ifelse(length(vp) > 3, i/4 - .1, i/3 - .18),
        side = 1, line = 3,outer = T)
  
  ## column titles
  if(length(vp) > 3){
    mtext(bquote('attack'), 
          side = 3, line = 2, outer = T, at = 0.12)
    mtext(bquote('conversion'), 
          side = 3, line = 2, outer = T, at = 0.37)
    mtext(bquote('mortality'), 
          side = 3, line = 2, outer = T, at = 0.63)
    mtext(bquote('handling'), 
          side = 3, line = 2, outer = T, at = 0.87)
  }else{
    mtext(bquote('attack'), 
          side = 3, line = 2, outer = T, at = 0.17)
    mtext(bquote('conversion'), 
          side = 3, line = 2, outer = T, at = 0.5)
    mtext(bquote('handling'), 
          side = 3, line = 2, outer = T, at = 0.82)
  }
  
}
#dev.off()
```
