## Supplementary material for "Socially transmitted innovations in dynamic predator-prey systems": All supplements: Figure6.r

```
library(deSolve)

t2 <- function(a, h, p, n) a*p*n/(1 + a*h*n) #type 2 functional response

m3 <- function(x){ #finds max, min, and mean for time series data
  a <- max(x) 
  b <- min(x)
  z <- mean(x)
  c(a,b,z)
}

## Figure 5 - Heatmap of cycle magnitude for sP1P2 transmission function
##            (can be modified to show other transmission functions)
##            Word to the wise - it takes a while to run

### The Models ###
## to be solved using dsolve ##

#mass action social learning depends on P1 and P2
RosenzweigMacArthur <- function(time, state, parameters) { 
  with(as.list(c(state, parameters)), {   
    dn <- n*r*(k-n)/k - t2(a1,h1,p1,n) - t2(a2,h2,p2,n)
    dp1 <- b1*t2(a1,h1,p1,n) + b2*t2(a2,h2,p2,n) - s*p1*p2 - m1*p1 
    dp2 <- s*p1*p2 - m2*p2
    return(list(c(dn, dp1, dp2)))
  })
}

## Specify Initial Conditions (for all models)
## Starts the system a little off equilibrium 
## for the n, p1 alone (basic consumer-resource) model
## in case the equilibrium point is unstable.
## Also gives the equilibrium point for the model
## when all individuals are uninformed and there
## is no social learning, a stable point of comparison
## under the default parameter values.
beg <- function(parama,offset){
  a1 = parama[[1]]; a2 = parama[[2]]; b1 = parama[[3]]; b2 = parama[[4]];
  m1 = parama[[5]]; m2 = parama[[6]]; h1 = parama[[7]]; h2 = parama[[8]];
  s = parama[[9]]; r = parama[[10]]; k = parama[[11]]; 
  n_star = m1/(a1*(b1-h1*m1))
  p1_star = b1*r*(a1*b1*k-m1-a1*h1*k*m1)/(a1^2 * k*(b1-h1*m1)^2)
  p2_star = 0
  init = c(n = n_star+offset, p1 = p1_star+offset, p2 = p2_star)
  init
}

## What values of the parameter we want to examine
## should we plot in the bifurcation diagram?
svals <- seq(0.05,1,length.out=30)
rvals <- seq(0.05,1,length.out=30) 

vp <- c(9,10) #index of which parameters are varying in Parms

## First, specify default parameter values
Parms <- list(a1 = 0.5, 
              a2 = 0.75,
              b1 = 0.27,  
              b2 = 0.27, 
              m1 = 0.2, 
              m2 = 0.2, 
              h1 = 1, 
              h2 = 1,
              s = 0.2, #default 0.2 
              r = 2,    #default 2 for b1 < b2, m1 > m2
              k = 10   #default 10
)   

ts <- 10000 #total time
asst <- 5000#burn-in time  

Parms[vp[1]] <- svals[1]
Parms[vp[2]] <- rvals[1]

off = 0.1 #perturbation from equilibrium for n and p1

## Specify simulation time
sim.time <- seq(0, ts, by = 0.1)

## Feed models, initial conditions and parameter values into solver
out_SLT2_2way <- ode(y=beg(Parms,0)+off, times=sim.time, 
                     func=RosenzweigMacArthur, parms=Parms)

## Begin bifurcation analyses to see where stability yields to cyclic
## dynamics

runs <- (nrow(out_SLT2_2way) - asst):nrow(out_SLT2_2way) #time interval to average over
lastbunch <- data.frame(out_SLT2_2way[runs,])
lastbunch$ptot <- rowSums(lastbunch[,3:4], na.rm = T)
lastbunch$p2tot <- lastbunch[,4]/lastbunch$ptot
lastbunch$pn <- lastbunch$ptot/lastbunch[,2]
colnames(lastbunch) <- c('time','n','p1','p2','ptot','p2tot','pn')

bifurc_ptot <- c(Parms[[vp[1]]],Parms[[vp[2]]],m3(lastbunch$ptot))
bifurc_p2tot <-  c(Parms[[vp[1]]],Parms[[vp[2]]],m3(lastbunch$p2tot))
bifurc_pn <-  c(Parms[[vp[1]]],Parms[[vp[2]]],m3(lastbunch$pn))
eq <- beg(Parms,0) #calculate the equilibrium solution for given parameter values

iters <- expand.grid(1:30,1:30)

for(i in 2:nrow(iters)){
  #auto-update focal parameters
  Parms[vp[1]] <- svals[iters[i,1]]
  Parms[vp[2]] <- rvals[iters[i,2]]
  
  out_SLT2_2way <- ode(y=beg(Parms,0)+off, times=sim.time, 
                       func=RosenzweigMacArthur, parms=Parms)
  lastbunch <- data.frame(out_SLT2_2way[runs,])
  lastbunch$ptot <- rowSums(lastbunch[,3:4], na.rm = T)
  lastbunch$p2tot <- lastbunch[,4]/lastbunch$ptot
  lastbunch$pn <- lastbunch$ptot/lastbunch[,2]
  colnames(lastbunch) <- c('time','n','p1','p2','ptot','p2tot','pn')
  
  bifurc_ptot <- rbind(bifurc_ptot,c(Parms[[vp[1]]],Parms[[vp[2]]],m3(lastbunch$ptot)))
  bifurc_p2tot <- rbind(bifurc_p2tot,c(Parms[[vp[1]]],Parms[[vp[2]]],m3(lastbunch$p2tot)))
  bifurc_pn <- rbind(bifurc_pn,c(Parms[[vp[1]]],Parms[[vp[2]]],m3(lastbunch$pn)))
  eq <- rbind(eq,beg(Parms,0)) #calculate the equilibrium solution for given parameter values
}

bifurc_ptot <- cbind(bifurc_ptot,eq[,2])
bifurc_p2tot <- cbind(bifurc_p2tot, NA)
bifurc_pn  <- cbind(bifurc_pn, eq[,2]/eq[,1])
cats <- rep(c('ptot','p2tot','pn'), each = nrow(bifurc_ptot))
op <- cbind(data.frame(rbind(bifurc_ptot,bifurc_p2tot,bifurc_pn)),cats)
n <- names(Parms)
nombres <- c(n[vp[1]],n[vp[2]],
             'max_2way','min_2way','mean_2way',
             'eq','var')
colnames(op) <- nombres
row.names(op) <- NULL 

#option to record data

#v <- unlist(Parms)
#fname <- paste('SLT2mods_s_r_grid_2way','.a1_',v[1],'.a2_',v[2],
#               '.b1_',v[3],'.b2_',v[4],'.m1_',v[5],'.m2_',v[6],'.s_',v[7],'.r_',v[8],
#               '.csv', sep = '')
#write.csv(op,fname,row.names=F)

# s vs r heatmap graphic

sr <- op
sr <- sr[sr$r <= 1,]
sr$rang <- with(sr,max_2way - min_2way)

datmatsr <- with(sr[which(sr$var == 'ptot'),], 
                 matrix(round(rang,4),30,30))

colfunc <- colorRampPalette(c("lightgray", "black"))
colly <- c('white',colfunc(30))
pdf('sr_stability1x_2way_Figure5.pdf',3.9,3)
par(mar = c(0,0,0,0), oma = c(4,3,1,6.5), mfrow = c(1,1),
    cex.axis = 0.7, mgp = c(0,0.3,0), tcl = -0.4)
image(datmatsr, zlim = range(datmatsr),
      yaxt = 'n', xaxt = 'n', col = colly)
box('plot')
axis(side = 1, at = c(0,0.48,1), labels = c(0.05,0.5,1))
axis(side = 2, at = c(0,1), labels = c(min(sr$r),round(max(sr$r))))
mtext(bquote(italic(r)), side = 2,  line = 1.5)
mtext(bquote(italic(s)), side = 1,  line = 1.5)

legend(1.2,1,legend = rev(c('stable',rep('',25),'cycles',rep('',3),'larger')),
       pch = 15, pt.cex = 2.7, y.intersp = 0.25,
       col = rev(colly), 
       xpd=NA, horiz = F)
dev.off()
```
