## Supplementary material for "Socially transmitted innovations in dynamic predator-prey systems": All supplements: SuppFig1.r

```
#Figure master script for supplementary figures describing
#de novo innovation versus social learning

par(mfcol = c(9,2), mar = c(0,0,0,0), oma = c(3,3,3,0.5))

library(deSolve)

#### Predator-prey dynamics when predators 
#### use social learning vs de novo innovation

## timeseries from Lotka-Volterra model where populations are dynamic
## and there is an innovation in attack rate.

init = c(n = 2, p1 = 3, p2 = 0.1)

sim.time <- seq(0, 5000, by = 0.1)

## Specify parameter values
Parms <- list(r = 0.4, 
              a1 = 0.15,
              a2 = 0.2, 
              b1 = 0.4, 
              b2 = 0.4,
              s = 0.17,
              d = 0.01,
              m1 = 0.2,
              m2 = 0.2)

#### Lotka-Volterra model ####
MODEL_denovo <- function(time, state, parameters) {
  with(as.list(c(state, parameters)), {       
    dn <-  n*(r-a1*p1-a2*p2)
    dp1 <-  a1*b1*p1*n+a2*b2*p2*n-d*p1-m1*p1
    dp2 <-  d*p1-m2*p2
    return(list(c(dn, dp1, dp2)))
  })
}

## Feed model, initial conditions and parameter values into solver
out <- ode(y=init, times=sim.time, func=MODEL_denovo, parms=Parms)
out = as.data.frame(out)

maa <- dim(out)[1]
#par(mfrow = c(3,1), mar = c(0,0,0,0), oma = c(2,2,2,0.5))
plot(out$time[0:1500], out$n[0:1500], xlim = c(0,240),
     pch=16,  type = 'l', lwd = 2, col = 'gray',
     ylim=c(0, max(out$n, out$p1, out$p2)+1),
     ylab="Abundance", xlab="Time", yaxt = 'n', xaxt = 'n')
lines(out$time[0:1500], out$p1[0:1500], lwd = 2, lty = 1)
lines(out$time[0:1500], out$p2[0:1500], lwd = 2, lty = 2)
lines(out$time[1800:2400], out$n[(maa-600):maa], lwd = 2, col = 'gray',)
lines(out$time[1800:2400], out$p1[(maa-600):maa], lwd = 2, lty = 1)
lines(out$time[1800:2400], out$p2[(maa-600):maa], lwd = 2, lty = 2)
text(240/60,(max(out$n, out$p1, out$p2)+0.5)/20*19.5, 
     pos = 4, 'attack: de novo', cex = 1.2)

#panel B

MODEL_social <- function(time, state, parameters) {
  with(as.list(c(state, parameters)), {       
    dn <-  n*(r-a1*p1-a2*p2)
    dp1 <-  a1*b1*p1*n+a2*b2*p2*n-s*p1*p2-m1*p1
    dp2 <-  s*p1*p2-m2*p2
    return(list(c(dn, dp1, dp2)))
  })
}

out_social <- ode(y=init, times=sim.time, func=MODEL_social, parms=Parms)
out_social = as.data.frame(out_social)

## plot this model run

plot(out_social$time[0:1500], out_social$n[0:1500], xlim = c(0,240),
     pch=16,  type = 'l', lwd = 2, col = 'gray',
     ylim=c(0, max(out_social$n, out_social$p1, out_social$p2)+1),
     ylab="Abundance", xlab="Time", yaxt = 'n', xaxt = 'n')
lines(out_social$time[0:1500], out_social$p1[0:1500], lwd = 2, lty = 1)
lines(out_social$time[0:1500], out_social$p2[0:1500], lwd = 2, lty = 2)
lines(out_social$time[1800:2400], out_social$n[(maa-600):maa], lwd = 2, col = 'gray',)
lines(out_social$time[1800:2400], out_social$p1[(maa-600):maa], lwd = 2, lty = 1)
lines(out_social$time[1800:2400], out_social$p2[(maa-600):maa], lwd = 2, lty = 2)

text(240/60,(max(out_social$n, out_social$p1, 
                 out_social$p2))/20*18 ,pos = 4,'attack: social', cex = 1.2)


#panel C
es <- Parms[[6]]
de <- Parms[[7]]
Parms[[6]] <- es*es/(es+de)
Parms[[7]] <- de*de/(es+de)

MODEL_denovo_social <- function(time, state, parameters) {
  with(as.list(c(state, parameters)), {       
    dn <-  n*(r-a1*p1-a2*p2)
    dp1 <-  a1*b1*p1*n+a2*b2*p2*n-s*p1*p2-d*p1-m1*p1
    dp2 <-  d*p1+s*p1*p2-m2*p2
    return(list(c(dn, dp1, dp2)))
  })
}

out_denovo_social <- ode(y=init, times=sim.time, func=MODEL_denovo_social, parms=Parms)
out_denovo_social = as.data.frame(out_denovo_social)

## plot this model run

plot(out_denovo_social$time[0:1500], out_denovo_social$n[0:1500], xlim = c(0,240),
     pch=16,  type = 'l', lwd = 2, col = 'gray',
     ylim=c(0, max(out_denovo_social$n, out_denovo_social$p1, out_denovo_social$p2)+1),
     ylab=bquote(italic('de novo')), xlab="Time", yaxt = 'n', xaxt = 'n')
lines(out_denovo_social$time[0:1500], out_denovo_social$p1[0:1500], lwd = 2, lty = 1)
lines(out_denovo_social$time[0:1500], out_denovo_social$p2[0:1500], lwd = 2, lty = 2)
lines(out_denovo_social$time[1800:2400], out_denovo_social$n[(maa-600):maa], lwd = 2, col = 'gray',)
lines(out_denovo_social$time[1800:2400], out_denovo_social$p1[(maa-600):maa], lwd = 2, lty = 1)
lines(out_denovo_social$time[1800:2400], out_denovo_social$p2[(maa-600):maa], lwd = 2, lty = 2)
text(240/60,(max(out_denovo_social$n, out_denovo_social$p1, 
                 out_denovo_social$p2))/20*18 ,pos = 4,'attack: de novo + social', cex = 1.2)


#############################################################################
#   Conversion rate
## Specify parameter values
Parms <- list(r = 0.4, 
              a1 = 0.15,
              a2 = 0.15, 
              b1 = 0.4, 
              b2 = 0.7,
              s = 0.17,
              d = 0.01,
              m1 = 0.2,
              m2 = 0.2)

#### Lotka-Volterra model ####
MODEL_denovo <- function(time, state, parameters) {
  with(as.list(c(state, parameters)), {       
    dn <-  n*(r-a1*p1-a2*p2)
    dp1 <-  a1*b1*p1*n+a2*b2*p2*n-d*p1-m1*p1
    dp2 <-  d*p1-m2*p2
    return(list(c(dn, dp1, dp2)))
  })
}

## Feed model, initial conditions and parameter values into solver
out <- ode(y=init, times=sim.time, func=MODEL_denovo, parms=Parms)
out = as.data.frame(out)

maa <- dim(out)[1]
#par(mfrow = c(3,1), mar = c(0,0,0,0), oma = c(2,2,2,0.5))
plot(out$time[0:1500], out$n[0:1500], xlim = c(0,240),
     pch=16,  type = 'l', lwd = 2, col = 'gray',
     ylim=c(0, max(out$n, out$p1, out$p2)+1),
     ylab="Abundance", xlab="Time", yaxt = 'n', xaxt = 'n')
lines(out$time[0:1500], out$p1[0:1500], lwd = 2, lty = 1)
lines(out$time[0:1500], out$p2[0:1500], lwd = 2, lty = 2)
lines(out$time[1800:2400], out$n[(maa-600):maa], lwd = 2, col = 'gray',)
lines(out$time[1800:2400], out$p1[(maa-600):maa], lwd = 2, lty = 1)
lines(out$time[1800:2400], out$p2[(maa-600):maa], lwd = 2, lty = 2)
text(240/60,(max(out$n, out$p1, out$p2)+0.5)/20*19.5, 
     pos = 4, 'conversion: de novo', cex = 1.2)

#panel B

MODEL_social <- function(time, state, parameters) {
  with(as.list(c(state, parameters)), {       
    dn <-  n*(r-a1*p1-a2*p2)
    dp1 <-  a1*b1*p1*n+a2*b2*p2*n-s*p1*p2-m1*p1
    dp2 <-  s*p1*p2-m2*p2
    return(list(c(dn, dp1, dp2)))
  })
}

out_social <- ode(y=init, times=sim.time, func=MODEL_social, parms=Parms)
out_social = as.data.frame(out_social)

## plot this model run

plot(out_social$time[0:1500], out_social$n[0:1500], xlim = c(0,240),
     pch=16,  type = 'l', lwd = 2, col = 'gray',
     ylim=c(0, max(out_social$n, out_social$p1, out_social$p2)+1),
     ylab="Abundance", xlab="Time", yaxt = 'n', xaxt = 'n')
lines(out_social$time[0:1500], out_social$p1[0:1500], lwd = 2, lty = 1)
lines(out_social$time[0:1500], out_social$p2[0:1500], lwd = 2, lty = 2)
lines(out_social$time[1800:2400], out_social$n[(maa-600):maa], lwd = 2, col = 'gray',)
lines(out_social$time[1800:2400], out_social$p1[(maa-600):maa], lwd = 2, lty = 1)
lines(out_social$time[1800:2400], out_social$p2[(maa-600):maa], lwd = 2, lty = 2)

text(240/60,(max(out_social$n, out_social$p1, 
                 out_social$p2)+0.5)/20*19.5 ,pos = 4,'conversion: social', cex = 1.2)


#panel C
es <- Parms[[6]]
de <- Parms[[7]]
Parms[[6]] <- es*es/(es+de)
Parms[[7]] <- de*de/(es+de)

MODEL_denovo_social <- function(time, state, parameters) {
  with(as.list(c(state, parameters)), {       
    dn <-  n*(r-a1*p1-a2*p2)
    dp1 <-  a1*b1*p1*n+a2*b2*p2*n-s*p1*p2-d*p1-m1*p1
    dp2 <-  d*p1+s*p1*p2-m2*p2
    return(list(c(dn, dp1, dp2)))
  })
}

out_denovo_social <- ode(y=init, times=sim.time, func=MODEL_denovo_social, parms=Parms)
out_denovo_social = as.data.frame(out_denovo_social)

## plot this model run

plot(out_denovo_social$time[0:1500], out_denovo_social$n[0:1500], xlim = c(0,240),
     pch=16,  type = 'l', lwd = 2, col = 'gray',
     ylim=c(0, max(out_denovo_social$n, out_denovo_social$p1, out_denovo_social$p2)+1),
     ylab=bquote(italic('de novo')), xlab="Time", yaxt = 'n', xaxt = 'n')
lines(out_denovo_social$time[0:1500], out_denovo_social$p1[0:1500], lwd = 2, lty = 1)
lines(out_denovo_social$time[0:1500], out_denovo_social$p2[0:1500], lwd = 2, lty = 2)
lines(out_denovo_social$time[1800:2400], out_denovo_social$n[(maa-600):maa], lwd = 2, col = 'gray',)
lines(out_denovo_social$time[1800:2400], out_denovo_social$p1[(maa-600):maa], lwd = 2, lty = 1)
lines(out_denovo_social$time[1800:2400], out_denovo_social$p2[(maa-600):maa], lwd = 2, lty = 2)
text(240/60,(max(out_denovo_social$n, out_denovo_social$p1, 
                 out_denovo_social$p2)+0.5)/20*19.5 ,pos = 4,'conversion: de novo + social', cex = 1.2)

#############################################################################
#   Mortality rate
## Specify parameter values
Parms <- list(r = 0.4, 
              a1 = 0.15,
              a2 = 0.15, 
              b1 = 0.4, 
              b2 = 0.4,
              s = 0.17,
              d = 0.01,
              m1 = 0.2,
              m2 = 0.12)

#### Lotka-Volterra model ####
MODEL_denovo <- function(time, state, parameters) {
  with(as.list(c(state, parameters)), {       
    dn <-  n*(r-a1*p1-a2*p2)
    dp1 <-  a1*b1*p1*n+a2*b2*p2*n-d*p1-m1*p1
    dp2 <-  d*p1-m2*p2
    return(list(c(dn, dp1, dp2)))
  })
}

## Feed model, initial conditions and parameter values into solver
out <- ode(y=init, times=sim.time, func=MODEL_denovo, parms=Parms)
out = as.data.frame(out)

maa <- dim(out)[1]
#par(mfrow = c(3,1), mar = c(0,0,0,0), oma = c(2,2,2,0.5))
plot(out$time[0:1500], out$n[0:1500], xlim = c(0,240),
     pch=16,  type = 'l', lwd = 2, col = 'gray',
     ylim=c(0, max(out$n, out$p1, out$p2)+1),
     ylab="Abundance", xlab="Time", yaxt = 'n', xaxt = 'n')
lines(out$time[0:1500], out$p1[0:1500], lwd = 2, lty = 1)
lines(out$time[0:1500], out$p2[0:1500], lwd = 2, lty = 2)
lines(out$time[1800:2400], out$n[(maa-600):maa], lwd = 2, col = 'gray',)
lines(out$time[1800:2400], out$p1[(maa-600):maa], lwd = 2, lty = 1)
lines(out$time[1800:2400], out$p2[(maa-600):maa], lwd = 2, lty = 2)
text(240/60,(max(out$n, out$p1, out$p2)+0.5)/20*19.5, 
     pos = 4, 'mortality: de novo', cex = 1.2)

#panel B

MODEL_social <- function(time, state, parameters) {
  with(as.list(c(state, parameters)), {       
    dn <-  n*(r-a1*p1-a2*p2)
    dp1 <-  a1*b1*p1*n+a2*b2*p2*n-s*p1*p2-m1*p1
    dp2 <-  s*p1*p2-m2*p2
    return(list(c(dn, dp1, dp2)))
  })
}

out_social <- ode(y=init, times=sim.time, func=MODEL_social, parms=Parms)
out_social = as.data.frame(out_social)

## plot this model run

plot(out_social$time[0:1500], out_social$n[0:1500], xlim = c(0,240),
     pch=16,  type = 'l', lwd = 2, col = 'gray',
     ylim=c(0, max(out_social$n, out_social$p1, out_social$p2)+1),
     ylab="Abundance", xlab="Time", yaxt = 'n', xaxt = 'n')
lines(out_social$time[0:1500], out_social$p1[0:1500], lwd = 2, lty = 1)
lines(out_social$time[0:1500], out_social$p2[0:1500], lwd = 2, lty = 2)
lines(out_social$time[1800:2400], out_social$n[(maa-600):maa], lwd = 2, col = 'gray',)
lines(out_social$time[1800:2400], out_social$p1[(maa-600):maa], lwd = 2, lty = 1)
lines(out_social$time[1800:2400], out_social$p2[(maa-600):maa], lwd = 2, lty = 2)

text(240/60,(max(out_social$n, out_social$p1, 
                 out_social$p2)+0.5)/20*19.5 ,pos = 4,'mortality: social', cex = 1.2)

#panel C
es <- Parms[[6]]
de <- Parms[[7]]
Parms[[6]] <- es*es/(es+de)
Parms[[7]] <- de*de/(es+de)

MODEL_denovo_social <- function(time, state, parameters) {
  with(as.list(c(state, parameters)), {       
    dn <-  n*(r-a1*p1-a2*p2)
    dp1 <-  a1*b1*p1*n+a2*b2*p2*n-s*p1*p2-d*p1-m1*p1
    dp2 <-  d*p1+s*p1*p2-m2*p2
    return(list(c(dn, dp1, dp2)))
  })
}

out_denovo_social <- ode(y=init, times=sim.time, func=MODEL_denovo_social, parms=Parms)
out_denovo_social = as.data.frame(out_denovo_social)

## plot this model run

plot(out_denovo_social$time[0:1500], out_denovo_social$n[0:1500], xlim = c(0,240),
     pch=16,  type = 'l', lwd = 2, col = 'gray',
     ylim=c(0, max(out_denovo_social$n, out_denovo_social$p1, out_denovo_social$p2)+1),
     ylab=bquote(italic('de novo')), xlab="Time", yaxt = 'n', xaxt = 'n')
lines(out_denovo_social$time[0:1500], out_denovo_social$p1[0:1500], lwd = 2, lty = 1)
lines(out_denovo_social$time[0:1500], out_denovo_social$p2[0:1500], lwd = 2, lty = 2)
lines(out_denovo_social$time[1800:2400], out_denovo_social$n[(maa-600):maa], lwd = 2, col = 'gray',)
lines(out_denovo_social$time[1800:2400], out_denovo_social$p1[(maa-600):maa], lwd = 2, lty = 1)
lines(out_denovo_social$time[1800:2400], out_denovo_social$p2[(maa-600):maa], lwd = 2, lty = 2)
text(240/60,(max(out_denovo_social$n, out_denovo_social$p1, 
                 out_denovo_social$p2)+0.5)/20*19.5 ,pos = 4,'mortality: de novo + social', cex = 1.2)
text(165,-.3,'//', cex = 1, xpd = NA)


############################################################################################################
############################################################################################################
############################################################################################################

# dP1N attack rate

Parms <- list(r = 0.4, 
              a1 = 0.15,
              a2 = 0.2, 
              b1 = 0.4, 
              b2 = 0.4,
              s = 0.17,
              d = 0.01,
              m1 = 0.2,
              m2 = 0.2)

MODEL_denovo <- function(time, state, parameters) {
  with(as.list(c(state, parameters)), {       
    dn <-   n*(r - a1*p1 - a2*p2)
    dp1 <-  a1*b1*p1*n + a2*b2*p2*n - d*p1*n - m1*p1
    dp2 <-  d*p1*n - m2*p2
    return(list(c(dn, dp1, dp2)))
  })
}

out <- ode(y=init, times=sim.time, func=MODEL_denovo, parms=Parms, 
           rtol = 1e-12, atol = 1e-12)
out = as.data.frame(out)

maa <- dim(out)[1]
plot(out$time[0:1500], out$n[0:1500], xlim = c(0,240),
     pch=16,  type = 'l', lwd = 2, col = 'gray',
     ylim=c(0, max(out$n, out$p1, out$p2)+1),
     ylab="Abundance", xlab="Time", yaxt = 'n', xaxt = 'n')
lines(out$time[0:1500], out$p1[0:1500], lwd = 2, lty = 1)
lines(out$time[0:1500], out$p2[0:1500], lwd = 2, lty = 2)
lines(out$time[1800:2400], out$n[(maa-600):maa], lwd = 2, col = 'gray',)
lines(out$time[1800:2400], out$p1[(maa-600):maa], lwd = 2, lty = 1)
lines(out$time[1800:2400], out$p2[(maa-600):maa], lwd = 2, lty = 2)


MODEL_social <- function(time, state, parameters) {
  with(as.list(c(state, parameters)), {       
    dn <-  n*(r-a1*p1-a2*p2)
    dp1 <-  a1*b1*p1*n+a2*b2*p2*n-s*p1*p2-m1*p1
    dp2 <-  s*p1*p2-m2*p2
    return(list(c(dn, dp1, dp2)))
  })
}

out_social <- ode(y=init, times=sim.time, func=MODEL_social, parms=Parms,
                  rtol = 1e-10, atol = 1e-10)
out_social = as.data.frame(out_social)

## plot this model run

plot(out_social$time[0:1500], out_social$n[0:1500], xlim = c(0,240),
     pch=16,  type = 'l', lwd = 2, col = 'gray',
     ylim=c(0, max(out_social$n, out_social$p1, out_social$p2)+1),
     ylab="Abundance", xlab="Time", yaxt = 'n', xaxt = 'n')
lines(out_social$time[0:1500], out_social$p1[0:1500], lwd = 2, lty = 1)
lines(out_social$time[0:1500], out_social$p2[0:1500], lwd = 2, lty = 2)
lines(out_social$time[1800:2400], out_social$n[(maa-600):maa], lwd = 2, col = 'gray',)
lines(out_social$time[1800:2400], out_social$p1[(maa-600):maa], lwd = 2, lty = 1)
lines(out_social$time[1800:2400], out_social$p2[(maa-600):maa], lwd = 2, lty = 2)

es <- Parms[[6]]
de <- Parms[[7]]
Parms[[6]] <- es*es/(es+de)
Parms[[7]] <- de*de/(es+de)

MODEL_denovo_social <- function(time, state, parameters) {
  with(as.list(c(state, parameters)), {       
    dn <-  n*(r-a1*p1-a2*p2)
    dp1 <-  a1*b1*p1*n+a2*b2*p2*n-s*p1*p2-d*p1*n-m1*p1
    dp2 <-  d*p1*n+s*p1*p2-m2*p2
    return(list(c(dn, dp1, dp2)))
  })
}

out_denovo_social <- ode(y=init, times=sim.time, 
                         func=MODEL_denovo_social, parms=Parms,
                         rtol = 1e-10, atol = 1e-10)
out_denovo_social = as.data.frame(out_denovo_social)

## plot this model run

plot(out_denovo_social$time[0:1500], out_denovo_social$n[0:1500], xlim = c(0,240),
     pch=16,  type = 'l', lwd = 2, col = 'gray',
     ylim=c(0, max(out_denovo_social$n, out_denovo_social$p1, out_denovo_social$p2)+1),
     ylab="Abundance", xlab="Time", yaxt = 'n', xaxt = 'n')
lines(out_denovo_social$time[0:1500], out_denovo_social$p1[0:1500], lwd = 2, lty = 1)
lines(out_denovo_social$time[0:1500], out_denovo_social$p2[0:1500], lwd = 2, lty = 2)
lines(out_denovo_social$time[1800:2400], out_denovo_social$n[(maa-600):maa], lwd = 2, col = 'gray',)
lines(out_denovo_social$time[1800:2400], out_denovo_social$p1[(maa-600):maa], lwd = 2, lty = 1)
lines(out_denovo_social$time[1800:2400], out_denovo_social$p2[(maa-600):maa], lwd = 2, lty = 2)

#****************************************************************************************#
# Conversion rate

Parms <- list(r = 0.4, 
              a1 = 0.15,
              a2 = 0.15, 
              b1 = 0.4, 
              b2 = 0.7,
              s = 0.17,
              d = 0.01,
              m1 = 0.2,
              m2 = 0.2)

MODEL_denovo <- function(time, state, parameters) {
  with(as.list(c(state, parameters)), {       
    dn <-   n*(r - a1*p1 - a2*p2)
    dp1 <-  a1*b1*p1*n + a2*b2*p2*n - d*p1*n - m1*p1
    dp2 <-  d*p1*n - m2*p2
    return(list(c(dn, dp1, dp2)))
  })
}

out <- ode(y=init, times=sim.time, func=MODEL_denovo, parms=Parms, 
           rtol = 1e-12, atol = 1e-12)
out = as.data.frame(out)

maa <- dim(out)[1]
plot(out$time[0:1500], out$n[0:1500], xlim = c(0,240),
     pch=16,  type = 'l', lwd = 2, col = 'gray',
     ylim=c(0, max(out$n, out$p1, out$p2)+1),
     ylab="Abundance", xlab="Time", yaxt = 'n', xaxt = 'n')
lines(out$time[0:1500], out$p1[0:1500], lwd = 2, lty = 1)
lines(out$time[0:1500], out$p2[0:1500], lwd = 2, lty = 2)
lines(out$time[1800:2400], out$n[(maa-600):maa], lwd = 2, col = 'gray',)
lines(out$time[1800:2400], out$p1[(maa-600):maa], lwd = 2, lty = 1)
lines(out$time[1800:2400], out$p2[(maa-600):maa], lwd = 2, lty = 2)


MODEL_social <- function(time, state, parameters) {
  with(as.list(c(state, parameters)), {       
    dn <-  n*(r-a1*p1-a2*p2)
    dp1 <-  a1*b1*p1*n+a2*b2*p2*n-s*p1*p2-m1*p1
    dp2 <-  s*p1*p2-m2*p2
    return(list(c(dn, dp1, dp2)))
  })
}

out_social <- ode(y=init, times=sim.time, func=MODEL_social, parms=Parms,
                  rtol = 1e-10, atol = 1e-10)
out_social = as.data.frame(out_social)

## plot this model run

plot(out_social$time[0:1500], out_social$n[0:1500], xlim = c(0,240),
     pch=16,  type = 'l', lwd = 2, col = 'gray',
     ylim=c(0, max(out_social$n, out_social$p1, out_social$p2)+1),
     ylab="Abundance", xlab="Time", yaxt = 'n', xaxt = 'n')
lines(out_social$time[0:1500], out_social$p1[0:1500], lwd = 2, lty = 1)
lines(out_social$time[0:1500], out_social$p2[0:1500], lwd = 2, lty = 2)
lines(out_social$time[1800:2400], out_social$n[(maa-600):maa], lwd = 2, col = 'gray',)
lines(out_social$time[1800:2400], out_social$p1[(maa-600):maa], lwd = 2, lty = 1)
lines(out_social$time[1800:2400], out_social$p2[(maa-600):maa], lwd = 2, lty = 2)

es <- Parms[[6]]
de <- Parms[[7]]
Parms[[6]] <- es*es/(es+de)
Parms[[7]] <- de*de/(es+de)

MODEL_denovo_social <- function(time, state, parameters) {
  with(as.list(c(state, parameters)), {       
    dn <-  n*(r-a1*p1-a2*p2)
    dp1 <-  a1*b1*p1*n+a2*b2*p2*n-s*p1*p2-d*p1*n-m1*p1
    dp2 <-  d*p1*n+s*p1*p2-m2*p2
    return(list(c(dn, dp1, dp2)))
  })
}

out_denovo_social <- ode(y=init, times=sim.time, 
                         func=MODEL_denovo_social, parms=Parms,
                         rtol = 1e-10, atol = 1e-10)
out_denovo_social = as.data.frame(out_denovo_social)

## plot this model run

plot(out_denovo_social$time[0:1500], out_denovo_social$n[0:1500], xlim = c(0,240),
     pch=16,  type = 'l', lwd = 2, col = 'gray',
     ylim=c(0, max(out_denovo_social$n, out_denovo_social$p1, out_denovo_social$p2)+1),
     ylab="Abundance", xlab="Time", yaxt = 'n', xaxt = 'n')
lines(out_denovo_social$time[0:1500], out_denovo_social$p1[0:1500], lwd = 2, lty = 1)
lines(out_denovo_social$time[0:1500], out_denovo_social$p2[0:1500], lwd = 2, lty = 2)
lines(out_denovo_social$time[1800:2400], out_denovo_social$n[(maa-600):maa], lwd = 2, col = 'gray',)
lines(out_denovo_social$time[1800:2400], out_denovo_social$p1[(maa-600):maa], lwd = 2, lty = 1)
lines(out_denovo_social$time[1800:2400], out_denovo_social$p2[(maa-600):maa], lwd = 2, lty = 2)

#****************************************************************************************#
# mortality rate

Parms <- list(r = 0.4, 
              a1 = 0.15,
              a2 = 0.15, 
              b1 = 0.4, 
              b2 = 0.4,
              s = 0.17,
              d = 0.01,
              m1 = 0.2,
              m2 = 0.12)

MODEL_denovo <- function(time, state, parameters) {
  with(as.list(c(state, parameters)), {       
    dn <-   n*(r - a1*p1 - a2*p2)
    dp1 <-  a1*b1*p1*n + a2*b2*p2*n - d*p1*n - m1*p1
    dp2 <-  d*p1*n - m2*p2
    return(list(c(dn, dp1, dp2)))
  })
}

out <- ode(y=init, times=sim.time, func=MODEL_denovo, parms=Parms, 
           rtol = 1e-12, atol = 1e-12)
out = as.data.frame(out)

maa <- dim(out)[1]
plot(out$time[0:1500], out$n[0:1500], xlim = c(0,240),
     pch=16,  type = 'l', lwd = 2, col = 'gray',
     ylim=c(0, max(out$n, out$p1, out$p2)+1),
     ylab="Abundance", xlab="Time", yaxt = 'n', xaxt = 'n')
lines(out$time[0:1500], out$p1[0:1500], lwd = 2, lty = 1)
lines(out$time[0:1500], out$p2[0:1500], lwd = 2, lty = 2)
lines(out$time[1800:2400], out$n[(maa-600):maa], lwd = 2, col = 'gray',)
lines(out$time[1800:2400], out$p1[(maa-600):maa], lwd = 2, lty = 1)
lines(out$time[1800:2400], out$p2[(maa-600):maa], lwd = 2, lty = 2)


MODEL_social <- function(time, state, parameters) {
  with(as.list(c(state, parameters)), {       
    dn <-  n*(r-a1*p1-a2*p2)
    dp1 <-  a1*b1*p1*n+a2*b2*p2*n-s*p1*p2-m1*p1
    dp2 <-  s*p1*p2-m2*p2
    return(list(c(dn, dp1, dp2)))
  })
}

out_social <- ode(y=init, times=sim.time, func=MODEL_social, parms=Parms,
                  rtol = 1e-10, atol = 1e-10)
out_social = as.data.frame(out_social)

## plot this model run

plot(out_social$time[0:1500], out_social$n[0:1500], xlim = c(0,240),
     pch=16,  type = 'l', lwd = 2, col = 'gray',
     ylim=c(0, max(out_social$n, out_social$p1, out_social$p2)+1),
     ylab="Abundance", xlab="Time", yaxt = 'n', xaxt = 'n')
lines(out_social$time[0:1500], out_social$p1[0:1500], lwd = 2, lty = 1)
lines(out_social$time[0:1500], out_social$p2[0:1500], lwd = 2, lty = 2)
lines(out_social$time[1800:2400], out_social$n[(maa-600):maa], lwd = 2, col = 'gray',)
lines(out_social$time[1800:2400], out_social$p1[(maa-600):maa], lwd = 2, lty = 1)
lines(out_social$time[1800:2400], out_social$p2[(maa-600):maa], lwd = 2, lty = 2)

es <- Parms[[6]]
de <- Parms[[7]]
Parms[[6]] <- es*es/(es+de)
Parms[[7]] <- de*de/(es+de)

MODEL_denovo_social <- function(time, state, parameters) {
  with(as.list(c(state, parameters)), {       
    dn <-  n*(r-a1*p1-a2*p2)
    dp1 <-  a1*b1*p1*n+a2*b2*p2*n-s*p1*p2-d*p1*n-m1*p1
    dp2 <-  d*p1*n+s*p1*p2-m2*p2
    return(list(c(dn, dp1, dp2)))
  })
}

out_denovo_social <- ode(y=init, times=sim.time, 
                         func=MODEL_denovo_social, parms=Parms,
                         rtol = 1e-10, atol = 1e-10)
out_denovo_social = as.data.frame(out_denovo_social)

## plot this model run

plot(out_denovo_social$time[0:1500], out_denovo_social$n[0:1500], xlim = c(0,240),
     pch=16,  type = 'l', lwd = 2, col = 'gray',
     ylim=c(0, max(out_denovo_social$n, out_denovo_social$p1, out_denovo_social$p2)+1),
     ylab="Abundance", xlab="Time", yaxt = 'n', xaxt = 'n')
lines(out_denovo_social$time[0:1500], out_denovo_social$p1[0:1500], lwd = 2, lty = 1)
lines(out_denovo_social$time[0:1500], out_denovo_social$p2[0:1500], lwd = 2, lty = 2)
lines(out_denovo_social$time[1800:2400], out_denovo_social$n[(maa-600):maa], lwd = 2, col = 'gray',)
lines(out_denovo_social$time[1800:2400], out_denovo_social$p1[(maa-600):maa], lwd = 2, lty = 1)
lines(out_denovo_social$time[1800:2400], out_denovo_social$p2[(maa-600):maa], lwd = 2, lty = 2)
text(165,-.3,'//', cex = 1, xpd = NA)

mtext('density', side = 2, line = 1.6, outer = T)
mtext('time', side = 1, line = 1.9, outer = T)
mtext(bquote(italic(dP[1])), side = 3, line = 1, outer = T, adj = 0.25)
mtext(bquote(italic(dP[1]*N)), side = 3, line = 1, outer = T, adj = 0.75)
```
