## Supplementary material for "Socially transmitted innovations in dynamic predator-prey systems": All supplements: SuppFig5.r

```
#Figure master script for supplementary figures describing
#de novo innovation versus social learning

par(mfcol = c(12,2), mar = c(0,0,0,0), oma = c(3,3,3,0.5))

library(deSolve)

#### Predator-prey dynamics when predators 
#### use social learning vs de novo innovation

## timeseries from Lotka-Volterra model where populations are dynamic
## and there is an innovation in attack rate.

init = c(n = 2, p1 = 3, p2 = 0.1)

sim.time <- seq(0, 500, by = 0.1)

t2 <- function(a, h, p, n) a*p*n/(1 + a*h*n) 

## Specify parameter values
Parms <- list(a1 = 0.5, #(0.5)
              a2 = 0.72, #(0.72)
              b1 = 0.27,  #(0.27)
              b2 = 0.27, #(0.31)
              m1 = 0.2,  #(0.2)
              m2 = 0.2, #(0.175)
              h1 = 1, 
              h2 = 1,
              s = 0.2, #default 0.16 
              d = 0.01,
              r = 2,   #default 2 for b1 < b2, m1 > m2
              k = 10   #default 10
)

#### Rosenzweig-MacArthur model ####
MODEL_denovo <- function(time, state, parameters) {
  with(as.list(c(state, parameters)), {       
    dn <-  n*r*(k-n)/k - t2(a1,h1,p1,n) - t2(a2,h2,p2,n)
    dp1 <-  b1*t2(a1,h1,p1,n) + b2*t2(a2,h2,p2,n)-d*p1-m1*p1
    dp2 <-  d*p1-m2*p2
    return(list(c(dn, dp1, dp2)))
  })
}

## Feed model, initial conditions and parameter values into solver
out <- ode(y=init, times=sim.time, func=MODEL_denovo, parms=Parms)
out = as.data.frame(out)

maa <- dim(out)[1]
plot(out$time[0:1500], out$n[0:1500], xlim = c(0,240),
     pch=16,  type = 'l', lwd = 2, col = 'gray',
     ylim=c(0, max(out$n, out$p1, out$p2)+1),
     ylab="Abundance", xlab="Time", yaxt = 'n', xaxt = 'n')
lines(out$time[0:1500], out$p1[0:1500], lwd = 2, lty = 1)
lines(out$time[0:1500], out$p2[0:1500], lwd = 2, lty = 2)
lines(out$time[1800:2400], out$n[(maa-600):maa], lwd = 2, col = 'gray',)
lines(out$time[1800:2400], out$p1[(maa-600):maa], lwd = 2, lty = 1)
lines(out$time[1800:2400], out$p2[(maa-600):maa], lwd = 2, lty = 2)
text(240/60,(max(out$n, out$p1, out$p2)+0.5)/20*19.5, 
     pos = 4, 'attack: de novo', cex = 1.2)

#panel B

MODEL_social <- function(time, state, parameters) {
  with(as.list(c(state, parameters)), {   
    dn <- n*r*(k-n)/k - t2(a1,h1,p1,n) - t2(a2,h2,p2,n)
    dp1 <- b1*t2(a1,h1,p1,n) + b2*t2(a2,h2,p2,n) - s*p1*p2 - m1*p1 #change to appropriate transmission function
    dp2 <- s*p1*p2  - m2*p2 #change to appropriate transmission function
    return(list(c(dn, dp1, dp2)))
  })
}

out_social <- ode(y=init, times=sim.time, func=MODEL_social, parms=Parms)
out_social = as.data.frame(out_social)

## plot this model run

plot(out_social$time[0:1500], out_social$n[0:1500], xlim = c(0,240),
     pch=16,  type = 'l', lwd = 2, col = 'gray',
     ylim=c(0, max(out_social$n, out_social$p1, out_social$p2)+1),
     ylab="Abundance", xlab="Time", yaxt = 'n', xaxt = 'n')
lines(out_social$time[0:1500], out_social$p1[0:1500], lwd = 2, lty = 1)
lines(out_social$time[0:1500], out_social$p2[0:1500], lwd = 2, lty = 2)
lines(out_social$time[1800:2400], out_social$n[(maa-600):maa], lwd = 2, col = 'gray',)
lines(out_social$time[1800:2400], out_social$p1[(maa-600):maa], lwd = 2, lty = 1)
lines(out_social$time[1800:2400], out_social$p2[(maa-600):maa], lwd = 2, lty = 2)

text(240/60,(max(out_social$n, out_social$p1, 
                 out_social$p2)+0.5)/20*19 ,pos = 4,'attack: social', cex = 1.2)


#panel C
es <- Parms[[9]]
de <- Parms[[10]]
Parms[[9]] <- es*es/(es+de)
Parms[[10]] <- de*de/(es+de)

MODEL_denovo_social <- function(time, state, parameters) {
  with(as.list(c(state, parameters)), {   
    dn <- n*r*(k-n)/k - t2(a1,h1,p1,n) - t2(a2,h2,p2,n)
    dp1 <- b1*t2(a1,h1,p1,n) + b2*t2(a2,h2,p2,n) - d*p1 - s*p1*p2 - m1*p1 #change to appropriate transmission function
    dp2 <- d*p1 + s*p1*p2  - m2*p2 #change to appropriate transmission function
    return(list(c(dn, dp1, dp2)))
  })
}

out_denovo_social <- ode(y=init, times=sim.time, func=MODEL_denovo_social, parms=Parms)
out_denovo_social = as.data.frame(out_denovo_social)

## plot this model run

plot(out_denovo_social$time[0:1500], out_denovo_social$n[0:1500], xlim = c(0,240),
     pch=16,  type = 'l', lwd = 2, col = 'gray',
     ylim=c(0, max(out_denovo_social$n, out_denovo_social$p1, out_denovo_social$p2)+1),
     ylab=bquote(italic('de novo')), xlab="Time", yaxt = 'n', xaxt = 'n')
lines(out_denovo_social$time[0:1500], out_denovo_social$p1[0:1500], lwd = 2, lty = 1)
lines(out_denovo_social$time[0:1500], out_denovo_social$p2[0:1500], lwd = 2, lty = 2)
lines(out_denovo_social$time[1800:2400], out_denovo_social$n[(maa-600):maa], lwd = 2, col = 'gray',)
lines(out_denovo_social$time[1800:2400], out_denovo_social$p1[(maa-600):maa], lwd = 2, lty = 1)
lines(out_denovo_social$time[1800:2400], out_denovo_social$p2[(maa-600):maa], lwd = 2, lty = 2)
text(240/60,(max(out_denovo_social$n, out_denovo_social$p1, 
                 out_denovo_social$p2)+0.5)/20*19 ,pos = 4,'attack: de novo + social', cex = 1.2)


#############################################################################
#   Conversion rate
## Specify parameter values
Parms <- list(a1 = 0.5, #(0.5)
              a2 = 0.5, #(0.72)
              b1 = 0.27,  #(0.27)
              b2 = 0.31, #(0.31)
              m1 = 0.2,  #(0.2)
              m2 = 0.2, #(0.175)
              h1 = 1, 
              h2 = 1,
              s = 0.2, #default 0.16 
              d = 0.01,
              r = 2,   #default 2 for b1 < b2, m1 > m2
              k = 10   #default 10
)

#### Rosenzweig-MacArthur model ####
MODEL_denovo <- function(time, state, parameters) {
  with(as.list(c(state, parameters)), {       
    dn <-  n*r*(k-n)/k - t2(a1,h1,p1,n) - t2(a2,h2,p2,n)
    dp1 <-  b1*t2(a1,h1,p1,n) + b2*t2(a2,h2,p2,n)-d*p1-m1*p1
    dp2 <-  d*p1-m2*p2
    return(list(c(dn, dp1, dp2)))
  })
}

## Feed model, initial conditions and parameter values into solver
out <- ode(y=init, times=sim.time, func=MODEL_denovo, parms=Parms)
out = as.data.frame(out)

maa <- dim(out)[1]
plot(out$time[0:1500], out$n[0:1500], xlim = c(0,240),
     pch=16,  type = 'l', lwd = 2, col = 'gray',
     ylim=c(0, max(out$n, out$p1, out$p2)+1),
     ylab="Abundance", xlab="Time", yaxt = 'n', xaxt = 'n')
lines(out$time[0:1500], out$p1[0:1500], lwd = 2, lty = 1)
lines(out$time[0:1500], out$p2[0:1500], lwd = 2, lty = 2)
lines(out$time[1800:2400], out$n[(maa-600):maa], lwd = 2, col = 'gray',)
lines(out$time[1800:2400], out$p1[(maa-600):maa], lwd = 2, lty = 1)
lines(out$time[1800:2400], out$p2[(maa-600):maa], lwd = 2, lty = 2)
text(240/60,(max(out$n, out$p1, out$p2)+0.5)/20*19.5, 
     pos = 4, 'conversion: de novo', cex = 1.2)

#panel B

MODEL_social <- function(time, state, parameters) {
  with(as.list(c(state, parameters)), {   
    dn <- n*r*(k-n)/k - t2(a1,h1,p1,n) - t2(a2,h2,p2,n)
    dp1 <- b1*t2(a1,h1,p1,n) + b2*t2(a2,h2,p2,n) - s*p1*p2 - m1*p1 #change to appropriate transmission function
    dp2 <- s*p1*p2  - m2*p2 #change to appropriate transmission function
    return(list(c(dn, dp1, dp2)))
  })
}

out_social <- ode(y=init, times=sim.time, func=MODEL_social, parms=Parms)
out_social = as.data.frame(out_social)

## plot this model run

plot(out_social$time[0:1500], out_social$n[0:1500], xlim = c(0,240),
     pch=16,  type = 'l', lwd = 2, col = 'gray',
     ylim=c(0, max(out_social$n, out_social$p1, out_social$p2)+1),
     ylab="Abundance", xlab="Time", yaxt = 'n', xaxt = 'n')
lines(out_social$time[0:1500], out_social$p1[0:1500], lwd = 2, lty = 1)
lines(out_social$time[0:1500], out_social$p2[0:1500], lwd = 2, lty = 2)
lines(out_social$time[1800:2400], out_social$n[(maa-600):maa], lwd = 2, col = 'gray',)
lines(out_social$time[1800:2400], out_social$p1[(maa-600):maa], lwd = 2, lty = 1)
lines(out_social$time[1800:2400], out_social$p2[(maa-600):maa], lwd = 2, lty = 2)

text(240/60,(max(out_social$n, out_social$p1, 
                 out_social$p2)+0.5)/20*19 ,pos = 4,'conversion: social', cex = 1.2)


#panel C
es <- Parms[[9]]
de <- Parms[[10]]
Parms[[9]] <- es*es/(es+de)
Parms[[10]] <- de*de/(es+de)

MODEL_denovo_social <- function(time, state, parameters) {
  with(as.list(c(state, parameters)), {   
    dn <- n*r*(k-n)/k - t2(a1,h1,p1,n) - t2(a2,h2,p2,n)
    dp1 <- b1*t2(a1,h1,p1,n) + b2*t2(a2,h2,p2,n) - d*p1 - s*p1*p2 - m1*p1 #change to appropriate transmission function
    dp2 <- d*p1 + s*p1*p2  - m2*p2 #change to appropriate transmission function
    return(list(c(dn, dp1, dp2)))
  })
}

out_denovo_social <- ode(y=init, times=sim.time, func=MODEL_denovo_social, parms=Parms)
out_denovo_social = as.data.frame(out_denovo_social)

## plot this model run

plot(out_denovo_social$time[0:1500], out_denovo_social$n[0:1500], xlim = c(0,240),
     pch=16,  type = 'l', lwd = 2, col = 'gray',
     ylim=c(0, max(out_denovo_social$n, out_denovo_social$p1, out_denovo_social$p2)+1),
     ylab=bquote(italic('de novo')), xlab="Time", yaxt = 'n', xaxt = 'n')
lines(out_denovo_social$time[0:1500], out_denovo_social$p1[0:1500], lwd = 2, lty = 1)
lines(out_denovo_social$time[0:1500], out_denovo_social$p2[0:1500], lwd = 2, lty = 2)
lines(out_denovo_social$time[1800:2400], out_denovo_social$n[(maa-600):maa], lwd = 2, col = 'gray',)
lines(out_denovo_social$time[1800:2400], out_denovo_social$p1[(maa-600):maa], lwd = 2, lty = 1)
lines(out_denovo_social$time[1800:2400], out_denovo_social$p2[(maa-600):maa], lwd = 2, lty = 2)
text(240/60,(max(out_denovo_social$n, out_denovo_social$p1, 
                 out_denovo_social$p2)+0.5)/20*19 ,pos = 4,'conversion: de novo + social', cex = 1.2)

#############################################################################
#   Mortality rate
## Specify parameter values

Parms <- list(a1 = 0.5, #(0.5)
              a2 = 0.5, #(0.72)
              b1 = 0.27,  #(0.27)
              b2 = 0.27, #(0.31)
              m1 = 0.2,  #(0.2)
              m2 = 0.175, #(0.175)
              h1 = 1, 
              h2 = 1,
              s = 0.2, #default 0.16 
              d = 0.01,
              r = 2,   #default 2 for b1 < b2, m1 > m2
              k = 10   #default 10
)

#### Rosenzweig-MacArthur model ####
MODEL_denovo <- function(time, state, parameters) {
  with(as.list(c(state, parameters)), {       
    dn <-  n*r*(k-n)/k - t2(a1,h1,p1,n) - t2(a2,h2,p2,n)
    dp1 <-  b1*t2(a1,h1,p1,n) + b2*t2(a2,h2,p2,n)-d*p1-m1*p1
    dp2 <-  d*p1-m2*p2
    return(list(c(dn, dp1, dp2)))
  })
}

## Feed model, initial conditions and parameter values into solver
out <- ode(y=init, times=sim.time, func=MODEL_denovo, parms=Parms)
out = as.data.frame(out)

maa <- dim(out)[1]
plot(out$time[0:1500], out$n[0:1500], xlim = c(0,240),
     pch=16,  type = 'l', lwd = 2, col = 'gray',
     ylim=c(0, max(out$n, out$p1, out$p2)+1),
     ylab="Abundance", xlab="Time", yaxt = 'n', xaxt = 'n')
lines(out$time[0:1500], out$p1[0:1500], lwd = 2, lty = 1)
lines(out$time[0:1500], out$p2[0:1500], lwd = 2, lty = 2)
lines(out$time[1800:2400], out$n[(maa-600):maa], lwd = 2, col = 'gray',)
lines(out$time[1800:2400], out$p1[(maa-600):maa], lwd = 2, lty = 1)
lines(out$time[1800:2400], out$p2[(maa-600):maa], lwd = 2, lty = 2)
text(240/60,(max(out$n, out$p1, out$p2)+0.5)/20*19.5, 
     pos = 4, 'mortality: de novo', cex = 1.2)

#panel B

MODEL_social <- function(time, state, parameters) {
  with(as.list(c(state, parameters)), {   
    dn <- n*r*(k-n)/k - t2(a1,h1,p1,n) - t2(a2,h2,p2,n)
    dp1 <- b1*t2(a1,h1,p1,n) + b2*t2(a2,h2,p2,n) - s*p1*p2 - m1*p1 #change to appropriate transmission function
    dp2 <- s*p1*p2  - m2*p2 #change to appropriate transmission function
    return(list(c(dn, dp1, dp2)))
  })
}

out_social <- ode(y=init, times=sim.time, func=MODEL_social, parms=Parms)
out_social = as.data.frame(out_social)

## plot this model run

plot(out_social$time[0:1500], out_social$n[0:1500], xlim = c(0,240),
     pch=16,  type = 'l', lwd = 2, col = 'gray',
     ylim=c(0, max(out_social$n, out_social$p1, out_social$p2)+1),
     ylab="Abundance", xlab="Time", yaxt = 'n', xaxt = 'n')
lines(out_social$time[0:1500], out_social$p1[0:1500], lwd = 2, lty = 1)
lines(out_social$time[0:1500], out_social$p2[0:1500], lwd = 2, lty = 2)
lines(out_social$time[1800:2400], out_social$n[(maa-600):maa], lwd = 2, col = 'gray',)
lines(out_social$time[1800:2400], out_social$p1[(maa-600):maa], lwd = 2, lty = 1)
lines(out_social$time[1800:2400], out_social$p2[(maa-600):maa], lwd = 2, lty = 2)

text(240/60,(max(out_social$n, out_social$p1, 
                 out_social$p2)+0.5)/20*19 ,pos = 4,'mortality: social', cex = 1.2)


#panel C
es <- Parms[[9]]
de <- Parms[[10]]
Parms[[9]] <- es*es/(es+de)
Parms[[10]] <- de*de/(es+de)

MODEL_denovo_social <- function(time, state, parameters) {
  with(as.list(c(state, parameters)), {   
    dn <- n*r*(k-n)/k - t2(a1,h1,p1,n) - t2(a2,h2,p2,n)
    dp1 <- b1*t2(a1,h1,p1,n) + b2*t2(a2,h2,p2,n) - d*p1 - s*p1*p2 - m1*p1 #change to appropriate transmission function
    dp2 <- d*p1 + s*p1*p2  - m2*p2 #change to appropriate transmission function
    return(list(c(dn, dp1, dp2)))
  })
}

out_denovo_social <- ode(y=init, times=sim.time, func=MODEL_denovo_social, parms=Parms)
out_denovo_social = as.data.frame(out_denovo_social)

## plot this model run

plot(out_denovo_social$time[0:1500], out_denovo_social$n[0:1500], xlim = c(0,240),
     pch=16,  type = 'l', lwd = 2, col = 'gray',
     ylim=c(0, max(out_denovo_social$n, out_denovo_social$p1, out_denovo_social$p2)+1),
     ylab=bquote(italic('de novo')), xlab="Time", yaxt = 'n', xaxt = 'n')
lines(out_denovo_social$time[0:1500], out_denovo_social$p1[0:1500], lwd = 2, lty = 1)
lines(out_denovo_social$time[0:1500], out_denovo_social$p2[0:1500], lwd = 2, lty = 2)
lines(out_denovo_social$time[1800:2400], out_denovo_social$n[(maa-600):maa], lwd = 2, col = 'gray',)
lines(out_denovo_social$time[1800:2400], out_denovo_social$p1[(maa-600):maa], lwd = 2, lty = 1)
lines(out_denovo_social$time[1800:2400], out_denovo_social$p2[(maa-600):maa], lwd = 2, lty = 2)
text(240/60,(max(out_denovo_social$n, out_denovo_social$p1, 
                 out_denovo_social$p2)+0.5)/20*19 ,pos = 4,'mortality: de novo + social', cex = 1.2)

##############################################################################
# handling time

Parms <- list(a1 = 0.5, #(0.5)
              a2 = 0.5, #(0.72)
              b1 = 0.27,  #(0.27)
              b2 = 0.27, #(0.31)
              m1 = 0.2,  #(0.2)
              m2 = 0.2, #(0.175)
              h1 = 1, 
              h2 = 0.5,
              s = 0.2, #default 0.16 
              d = 0.01,
              r = 2,   #default 2 for b1 < b2, m1 > m2
              k = 10   #default 10
)

#### Rosenzweig-MacArthur model ####
MODEL_denovo <- function(time, state, parameters) {
  with(as.list(c(state, parameters)), {       
    dn <-  n*r*(k-n)/k - t2(a1,h1,p1,n) - t2(a2,h2,p2,n)
    dp1 <-  b1*t2(a1,h1,p1,n) + b2*t2(a2,h2,p2,n)-d*p1-m1*p1
    dp2 <-  d*p1-m2*p2
    return(list(c(dn, dp1, dp2)))
  })
}

## Feed model, initial conditions and parameter values into solver
out <- ode(y=init, times=sim.time, func=MODEL_denovo, parms=Parms)
out = as.data.frame(out)

maa <- dim(out)[1]
plot(out$time[0:1500], out$n[0:1500], xlim = c(0,240),
     pch=16,  type = 'l', lwd = 2, col = 'gray',
     ylim=c(0, max(out$n, out$p1, out$p2)+1),
     ylab="Abundance", xlab="Time", yaxt = 'n', xaxt = 'n')
lines(out$time[0:1500], out$p1[0:1500], lwd = 2, lty = 1)
lines(out$time[0:1500], out$p2[0:1500], lwd = 2, lty = 2)
lines(out$time[1800:2400], out$n[(maa-600):maa], lwd = 2, col = 'gray',)
lines(out$time[1800:2400], out$p1[(maa-600):maa], lwd = 2, lty = 1)
lines(out$time[1800:2400], out$p2[(maa-600):maa], lwd = 2, lty = 2)
text(240/60,(max(out$n, out$p1, out$p2)+0.5)/20*19.5, 
     pos = 4, 'handling: de novo', cex = 1.2)

#panel B

MODEL_social <- function(time, state, parameters) {
  with(as.list(c(state, parameters)), {   
    dn <- n*r*(k-n)/k - t2(a1,h1,p1,n) - t2(a2,h2,p2,n)
    dp1 <- b1*t2(a1,h1,p1,n) + b2*t2(a2,h2,p2,n) - s*p1*p2 - m1*p1 #change to appropriate transmission function
    dp2 <- s*p1*p2  - m2*p2 #change to appropriate transmission function
    return(list(c(dn, dp1, dp2)))
  })
}

out_social <- ode(y=init, times=sim.time, func=MODEL_social, parms=Parms)
out_social = as.data.frame(out_social)

## plot this model run

plot(out_social$time[0:1500], out_social$n[0:1500], xlim = c(0,240),
     pch=16,  type = 'l', lwd = 2, col = 'gray',
     ylim=c(0, max(out_social$n, out_social$p1, out_social$p2)+1),
     ylab="Abundance", xlab="Time", yaxt = 'n', xaxt = 'n')
lines(out_social$time[0:1500], out_social$p1[0:1500], lwd = 2, lty = 1)
lines(out_social$time[0:1500], out_social$p2[0:1500], lwd = 2, lty = 2)
lines(out_social$time[1800:2400], out_social$n[(maa-600):maa], lwd = 2, col = 'gray',)
lines(out_social$time[1800:2400], out_social$p1[(maa-600):maa], lwd = 2, lty = 1)
lines(out_social$time[1800:2400], out_social$p2[(maa-600):maa], lwd = 2, lty = 2)

text(240/60,(max(out_social$n, out_social$p1, 
                 out_social$p2)+0.5)/20*19 ,pos = 4,'handling: social', cex = 1.2)


#panel C
es <- Parms[[9]]
de <- Parms[[10]]
Parms[[9]] <- es*es/(es+de)
Parms[[10]] <- de*de/(es+de)

MODEL_denovo_social <- function(time, state, parameters) {
  with(as.list(c(state, parameters)), {   
    dn <- n*r*(k-n)/k - t2(a1,h1,p1,n) - t2(a2,h2,p2,n)
    dp1 <- b1*t2(a1,h1,p1,n) + b2*t2(a2,h2,p2,n) - d*p1 - s*p1*p2 - m1*p1 #change to appropriate transmission function
    dp2 <- d*p1 + s*p1*p2  - m2*p2 #change to appropriate transmission function
    return(list(c(dn, dp1, dp2)))
  })
}

out_denovo_social <- ode(y=init, times=sim.time, func=MODEL_denovo_social, parms=Parms)
out_denovo_social = as.data.frame(out_denovo_social)

## plot this model run

plot(out_denovo_social$time[0:1500], out_denovo_social$n[0:1500], xlim = c(0,240),
     pch=16,  type = 'l', lwd = 2, col = 'gray',
     ylim=c(0, max(out_denovo_social$n, out_denovo_social$p1, out_denovo_social$p2)+1),
     ylab=bquote(italic('de novo')), xlab="Time", yaxt = 'n', xaxt = 'n')
lines(out_denovo_social$time[0:1500], out_denovo_social$p1[0:1500], lwd = 2, lty = 1)
lines(out_denovo_social$time[0:1500], out_denovo_social$p2[0:1500], lwd = 2, lty = 2)
lines(out_denovo_social$time[1800:2400], out_denovo_social$n[(maa-600):maa], lwd = 2, col = 'gray',)
lines(out_denovo_social$time[1800:2400], out_denovo_social$p1[(maa-600):maa], lwd = 2, lty = 1)
lines(out_denovo_social$time[1800:2400], out_denovo_social$p2[(maa-600):maa], lwd = 2, lty = 2)
text(240/60,(max(out_denovo_social$n, out_denovo_social$p1, 
                 out_denovo_social$p2)+0.5)/20*19 ,pos = 4,'handling: de novo + social', cex = 1.2)
text(165,-.3,'//', cex = 1, xpd = NA)

############################################################################################################
############################################################################################################
############################################################################################################

# dP1N attack rate
## Specify parameter values
Parms <- list(a1 = 0.5, #(0.5)
              a2 = 0.72, #(0.72)
              b1 = 0.27,  #(0.27)
              b2 = 0.27, #(0.31)
              m1 = 0.2,  #(0.2)
              m2 = 0.2, #(0.175)
              h1 = 1, 
              h2 = 1,
              s = 0.2, #default 0.16 
              d = 0.01,
              r = 2,   #default 2 for b1 < b2, m1 > m2
              k = 10   #default 10
)

#### Rosenzweig-MacArthur model ####
MODEL_denovo <- function(time, state, parameters) {
  with(as.list(c(state, parameters)), {       
    dn <-  n*r*(k-n)/k - t2(a1,h1,p1,n) - t2(a2,h2,p2,n)
    dp1 <-  b1*t2(a1,h1,p1,n) + b2*t2(a2,h2,p2,n)-d*p1*n-m1*p1
    dp2 <-  d*p1*n-m2*p2
    return(list(c(dn, dp1, dp2)))
  })
}

## Feed model, initial conditions and parameter values into solver
out <- ode(y=init, times=sim.time, func=MODEL_denovo, parms=Parms)
out = as.data.frame(out)

maa <- dim(out)[1]
plot(out$time[0:1500], out$n[0:1500], xlim = c(0,240),
     pch=16,  type = 'l', lwd = 2, col = 'gray',
     ylim=c(0, max(out$n, out$p1, out$p2)+1),
     ylab="Abundance", xlab="Time", yaxt = 'n', xaxt = 'n')
lines(out$time[0:1500], out$p1[0:1500], lwd = 2, lty = 1)
lines(out$time[0:1500], out$p2[0:1500], lwd = 2, lty = 2)
lines(out$time[1800:2400], out$n[(maa-600):maa], lwd = 2, col = 'gray',)
lines(out$time[1800:2400], out$p1[(maa-600):maa], lwd = 2, lty = 1)
lines(out$time[1800:2400], out$p2[(maa-600):maa], lwd = 2, lty = 2)

#panel B

MODEL_social <- function(time, state, parameters) {
  with(as.list(c(state, parameters)), {   
    dn <- n*r*(k-n)/k - t2(a1,h1,p1,n) - t2(a2,h2,p2,n)
    dp1 <- b1*t2(a1,h1,p1,n) + b2*t2(a2,h2,p2,n) - s*n*p1*p2 - m1*p1 #change to appropriate transmission function
    dp2 <- s*n*p1*p2  - m2*p2 #change to appropriate transmission function
    return(list(c(dn, dp1, dp2)))
  })
}

out_social <- ode(y=init, times=sim.time, func=MODEL_social, parms=Parms)
out_social = as.data.frame(out_social)

## plot this model run

plot(out_social$time[0:1500], out_social$n[0:1500], xlim = c(0,240),
     pch=16,  type = 'l', lwd = 2, col = 'gray',
     ylim=c(0, max(out_social$n, out_social$p1, out_social$p2)+1),
     ylab="Abundance", xlab="Time", yaxt = 'n', xaxt = 'n')
lines(out_social$time[0:1500], out_social$p1[0:1500], lwd = 2, lty = 1)
lines(out_social$time[0:1500], out_social$p2[0:1500], lwd = 2, lty = 2)
lines(out_social$time[1800:2400], out_social$n[(maa-600):maa], lwd = 2, col = 'gray',)
lines(out_social$time[1800:2400], out_social$p1[(maa-600):maa], lwd = 2, lty = 1)
lines(out_social$time[1800:2400], out_social$p2[(maa-600):maa], lwd = 2, lty = 2)

#panel C
es <- Parms[[9]]
de <- Parms[[10]]
Parms[[9]] <- es*es/(es+de)
Parms[[10]] <- de*de/(es+de)

MODEL_denovo_social <- function(time, state, parameters) {
  with(as.list(c(state, parameters)), {   
    dn <- n*r*(k-n)/k - t2(a1,h1,p1,n) - t2(a2,h2,p2,n)
    dp1 <- b1*t2(a1,h1,p1,n) + b2*t2(a2,h2,p2,n) - d*p1*n - s*n*p1*p2 - m1*p1 #change to appropriate transmission function
    dp2 <- d*p1*n + s*n*p1*p2  - m2*p2 #change to appropriate transmission function
    return(list(c(dn, dp1, dp2)))
  })
}

out_denovo_social <- ode(y=init, times=sim.time, func=MODEL_denovo_social, parms=Parms)
out_denovo_social = as.data.frame(out_denovo_social)

## plot this model run

plot(out_denovo_social$time[0:1500], out_denovo_social$n[0:1500], xlim = c(0,240),
     pch=16,  type = 'l', lwd = 2, col = 'gray',
     ylim=c(0, max(out_denovo_social$n, out_denovo_social$p1, out_denovo_social$p2)+1),
     ylab=bquote(italic('de novo')), xlab="Time", yaxt = 'n', xaxt = 'n')
lines(out_denovo_social$time[0:1500], out_denovo_social$p1[0:1500], lwd = 2, lty = 1)
lines(out_denovo_social$time[0:1500], out_denovo_social$p2[0:1500], lwd = 2, lty = 2)
lines(out_denovo_social$time[1800:2400], out_denovo_social$n[(maa-600):maa], lwd = 2, col = 'gray',)
lines(out_denovo_social$time[1800:2400], out_denovo_social$p1[(maa-600):maa], lwd = 2, lty = 1)
lines(out_denovo_social$time[1800:2400], out_denovo_social$p2[(maa-600):maa], lwd = 2, lty = 2)

#############################################################################
#   Conversion rate
## Specify parameter values
Parms <- list(a1 = 0.5, #(0.5)
              a2 = 0.5, #(0.72)
              b1 = 0.27,  #(0.27)
              b2 = 0.31, #(0.31)
              m1 = 0.2,  #(0.2)
              m2 = 0.2, #(0.175)
              h1 = 1, 
              h2 = 1,
              s = 0.2, #default 0.16 
              d = 0.01,
              r = 2,   #default 2 for b1 < b2, m1 > m2
              k = 10   #default 10
)

#### Rosenzweig-MacArthur model ####
MODEL_denovo <- function(time, state, parameters) {
  with(as.list(c(state, parameters)), {       
    dn <-  n*r*(k-n)/k - t2(a1,h1,p1,n) - t2(a2,h2,p2,n)
    dp1 <-  b1*t2(a1,h1,p1,n) + b2*t2(a2,h2,p2,n)-d*p1*n-m1*p1
    dp2 <-  d*p1*n-m2*p2
    return(list(c(dn, dp1, dp2)))
  })
}

## Feed model, initial conditions and parameter values into solver
out <- ode(y=init, times=sim.time, func=MODEL_denovo, parms=Parms)
out = as.data.frame(out)

maa <- dim(out)[1]
plot(out$time[0:1500], out$n[0:1500], xlim = c(0,240),
     pch=16,  type = 'l', lwd = 2, col = 'gray',
     ylim=c(0, max(out$n, out$p1, out$p2)+1),
     ylab="Abundance", xlab="Time", yaxt = 'n', xaxt = 'n')
lines(out$time[0:1500], out$p1[0:1500], lwd = 2, lty = 1)
lines(out$time[0:1500], out$p2[0:1500], lwd = 2, lty = 2)
lines(out$time[1800:2400], out$n[(maa-600):maa], lwd = 2, col = 'gray',)
lines(out$time[1800:2400], out$p1[(maa-600):maa], lwd = 2, lty = 1)
lines(out$time[1800:2400], out$p2[(maa-600):maa], lwd = 2, lty = 2)

#panel B

MODEL_social <- function(time, state, parameters) {
  with(as.list(c(state, parameters)), {   
    dn <- n*r*(k-n)/k - t2(a1,h1,p1,n) - t2(a2,h2,p2,n)
    dp1 <- b1*t2(a1,h1,p1,n) + b2*t2(a2,h2,p2,n) - s*n*p1*p2 - m1*p1 #change to appropriate transmission function
    dp2 <- s*n*p1*p2  - m2*p2 #change to appropriate transmission function
    return(list(c(dn, dp1, dp2)))
  })
}

out_social <- ode(y=init, times=sim.time, func=MODEL_social, parms=Parms)
out_social = as.data.frame(out_social)

## plot this model run

plot(out_social$time[0:1500], out_social$n[0:1500], xlim = c(0,240),
     pch=16,  type = 'l', lwd = 2, col = 'gray',
     ylim=c(0, max(out_social$n, out_social$p1, out_social$p2)+1),
     ylab="Abundance", xlab="Time", yaxt = 'n', xaxt = 'n')
lines(out_social$time[0:1500], out_social$p1[0:1500], lwd = 2, lty = 1)
lines(out_social$time[0:1500], out_social$p2[0:1500], lwd = 2, lty = 2)
lines(out_social$time[1800:2400], out_social$n[(maa-600):maa], lwd = 2, col = 'gray',)
lines(out_social$time[1800:2400], out_social$p1[(maa-600):maa], lwd = 2, lty = 1)
lines(out_social$time[1800:2400], out_social$p2[(maa-600):maa], lwd = 2, lty = 2)

#panel C
es <- Parms[[9]]
de <- Parms[[10]]
Parms[[9]] <- es*es/(es+de)
Parms[[10]] <- de*de/(es+de)

MODEL_denovo_social <- function(time, state, parameters) {
  with(as.list(c(state, parameters)), {   
    dn <- n*r*(k-n)/k - t2(a1,h1,p1,n) - t2(a2,h2,p2,n)
    dp1 <- b1*t2(a1,h1,p1,n) + b2*t2(a2,h2,p2,n) - d*p1*n - s*n*p1*p2 - m1*p1 #change to appropriate transmission function
    dp2 <- d*p1*n + s*n*p1*p2  - m2*p2 #change to appropriate transmission function
    return(list(c(dn, dp1, dp2)))
  })
}

out_denovo_social <- ode(y=init, times=sim.time, func=MODEL_denovo_social, parms=Parms)
out_denovo_social = as.data.frame(out_denovo_social)

## plot this model run

plot(out_denovo_social$time[0:1500], out_denovo_social$n[0:1500], xlim = c(0,240),
     pch=16,  type = 'l', lwd = 2, col = 'gray',
     ylim=c(0, max(out_denovo_social$n, out_denovo_social$p1, out_denovo_social$p2)+1),
     ylab=bquote(italic('de novo')), xlab="Time", yaxt = 'n', xaxt = 'n')
lines(out_denovo_social$time[0:1500], out_denovo_social$p1[0:1500], lwd = 2, lty = 1)
lines(out_denovo_social$time[0:1500], out_denovo_social$p2[0:1500], lwd = 2, lty = 2)
lines(out_denovo_social$time[1800:2400], out_denovo_social$n[(maa-600):maa], lwd = 2, col = 'gray',)
lines(out_denovo_social$time[1800:2400], out_denovo_social$p1[(maa-600):maa], lwd = 2, lty = 1)
lines(out_denovo_social$time[1800:2400], out_denovo_social$p2[(maa-600):maa], lwd = 2, lty = 2)

#############################################################################
#   Mortality rate
## Specify parameter values

Parms <- list(a1 = 0.5, #(0.5)
              a2 = 0.5, #(0.72)
              b1 = 0.27,  #(0.27)
              b2 = 0.27, #(0.31)
              m1 = 0.2,  #(0.2)
              m2 = 0.175, #(0.175)
              h1 = 1, 
              h2 = 1,
              s = 0.2, #default 0.16 
              d = 0.01,
              r = 2,   #default 2 for b1 < b2, m1 > m2
              k = 10   #default 10
)

#### Rosenzweig-MacArthur model ####
MODEL_denovo <- function(time, state, parameters) {
  with(as.list(c(state, parameters)), {       
    dn <-  n*r*(k-n)/k - t2(a1,h1,p1,n) - t2(a2,h2,p2,n)
    dp1 <-  b1*t2(a1,h1,p1,n) + b2*t2(a2,h2,p2,n)-d*p1*n-m1*p1
    dp2 <-  d*p1*n-m2*p2
    return(list(c(dn, dp1, dp2)))
  })
}

## Feed model, initial conditions and parameter values into solver
out <- ode(y=init, times=sim.time, func=MODEL_denovo, parms=Parms)
out = as.data.frame(out)

maa <- dim(out)[1]
plot(out$time[0:1500], out$n[0:1500], xlim = c(0,240),
     pch=16,  type = 'l', lwd = 2, col = 'gray',
     ylim=c(0, max(out$n, out$p1, out$p2)+1),
     ylab="Abundance", xlab="Time", yaxt = 'n', xaxt = 'n')
lines(out$time[0:1500], out$p1[0:1500], lwd = 2, lty = 1)
lines(out$time[0:1500], out$p2[0:1500], lwd = 2, lty = 2)
lines(out$time[1800:2400], out$n[(maa-600):maa], lwd = 2, col = 'gray',)
lines(out$time[1800:2400], out$p1[(maa-600):maa], lwd = 2, lty = 1)
lines(out$time[1800:2400], out$p2[(maa-600):maa], lwd = 2, lty = 2)

#panel B

MODEL_social <- function(time, state, parameters) {
  with(as.list(c(state, parameters)), {   
    dn <- n*r*(k-n)/k - t2(a1,h1,p1,n) - t2(a2,h2,p2,n)
    dp1 <- b1*t2(a1,h1,p1,n) + b2*t2(a2,h2,p2,n) - s*n*p1*p2 - m1*p1 #change to appropriate transmission function
    dp2 <- s*n*p1*p2  - m2*p2 #change to appropriate transmission function
    return(list(c(dn, dp1, dp2)))
  })
}

out_social <- ode(y=init, times=sim.time, func=MODEL_social, parms=Parms)
out_social = as.data.frame(out_social)

## plot this model run

plot(out_social$time[0:1500], out_social$n[0:1500], xlim = c(0,240),
     pch=16,  type = 'l', lwd = 2, col = 'gray',
     ylim=c(0, max(out_social$n, out_social$p1, out_social$p2)+1),
     ylab="Abundance", xlab="Time", yaxt = 'n', xaxt = 'n')
lines(out_social$time[0:1500], out_social$p1[0:1500], lwd = 2, lty = 1)
lines(out_social$time[0:1500], out_social$p2[0:1500], lwd = 2, lty = 2)
lines(out_social$time[1800:2400], out_social$n[(maa-600):maa], lwd = 2, col = 'gray',)
lines(out_social$time[1800:2400], out_social$p1[(maa-600):maa], lwd = 2, lty = 1)
lines(out_social$time[1800:2400], out_social$p2[(maa-600):maa], lwd = 2, lty = 2)

#panel C
es <- Parms[[9]]
de <- Parms[[10]]
Parms[[9]] <- es*es/(es+de)
Parms[[10]] <- de*de/(es+de)

MODEL_denovo_social <- function(time, state, parameters) {
  with(as.list(c(state, parameters)), {   
    dn <- n*r*(k-n)/k - t2(a1,h1,p1,n) - t2(a2,h2,p2,n)
    dp1 <- b1*t2(a1,h1,p1,n) + b2*t2(a2,h2,p2,n) - d*p1*n - s*n*p1*p2 - m1*p1 #change to appropriate transmission function
    dp2 <- d*p1*n + s*n*p1*p2  - m2*p2 #change to appropriate transmission function
    return(list(c(dn, dp1, dp2)))
  })
}

out_denovo_social <- ode(y=init, times=sim.time, func=MODEL_denovo_social, parms=Parms)
out_denovo_social = as.data.frame(out_denovo_social)

## plot this model run

plot(out_denovo_social$time[0:1500], out_denovo_social$n[0:1500], xlim = c(0,240),
     pch=16,  type = 'l', lwd = 2, col = 'gray',
     ylim=c(0, max(out_denovo_social$n, out_denovo_social$p1, out_denovo_social$p2)+1),
     ylab=bquote(italic('de novo')), xlab="Time", yaxt = 'n', xaxt = 'n')
lines(out_denovo_social$time[0:1500], out_denovo_social$p1[0:1500], lwd = 2, lty = 1)
lines(out_denovo_social$time[0:1500], out_denovo_social$p2[0:1500], lwd = 2, lty = 2)
lines(out_denovo_social$time[1800:2400], out_denovo_social$n[(maa-600):maa], lwd = 2, col = 'gray',)
lines(out_denovo_social$time[1800:2400], out_denovo_social$p1[(maa-600):maa], lwd = 2, lty = 1)
lines(out_denovo_social$time[1800:2400], out_denovo_social$p2[(maa-600):maa], lwd = 2, lty = 2)

##############################################################################
# handling time

Parms <- list(a1 = 0.5, #(0.5)
              a2 = 0.5, #(0.72)
              b1 = 0.27,  #(0.27)
              b2 = 0.27, #(0.31)
              m1 = 0.2,  #(0.2)
              m2 = 0.2, #(0.175)
              h1 = 1, 
              h2 = 0.5,
              s = 0.2, #default 0.16 
              d = 0.01,
              r = 2,   #default 2 for b1 < b2, m1 > m2
              k = 10   #default 10
)

#### Rosenzweig-MacArthur model ####
MODEL_denovo <- function(time, state, parameters) {
  with(as.list(c(state, parameters)), {       
    dn <-  n*r*(k-n)/k - t2(a1,h1,p1,n) - t2(a2,h2,p2,n)
    dp1 <-  b1*t2(a1,h1,p1,n) + b2*t2(a2,h2,p2,n)-d*p1*n-m1*p1
    dp2 <-  d*p1*n-m2*p2
    return(list(c(dn, dp1, dp2)))
  })
}

## Feed model, initial conditions and parameter values into solver
out <- ode(y=init, times=sim.time, func=MODEL_denovo, parms=Parms)
out = as.data.frame(out)

maa <- dim(out)[1]
plot(out$time[0:1500], out$n[0:1500], xlim = c(0,240),
     pch=16,  type = 'l', lwd = 2, col = 'gray',
     ylim=c(0, max(out$n, out$p1, out$p2)+1),
     ylab="Abundance", xlab="Time", yaxt = 'n', xaxt = 'n')
lines(out$time[0:1500], out$p1[0:1500], lwd = 2, lty = 1)
lines(out$time[0:1500], out$p2[0:1500], lwd = 2, lty = 2)
lines(out$time[1800:2400], out$n[(maa-600):maa], lwd = 2, col = 'gray',)
lines(out$time[1800:2400], out$p1[(maa-600):maa], lwd = 2, lty = 1)
lines(out$time[1800:2400], out$p2[(maa-600):maa], lwd = 2, lty = 2)

#panel B

MODEL_social <- function(time, state, parameters) {
  with(as.list(c(state, parameters)), {   
    dn <- n*r*(k-n)/k - t2(a1,h1,p1,n) - t2(a2,h2,p2,n)
    dp1 <- b1*t2(a1,h1,p1,n) + b2*t2(a2,h2,p2,n) - s*n*p1*p2 - m1*p1 #change to appropriate transmission function
    dp2 <- s*n*p1*p2  - m2*p2 #change to appropriate transmission function
    return(list(c(dn, dp1, dp2)))
  })
}

out_social <- ode(y=init, times=sim.time, func=MODEL_social, parms=Parms)
out_social = as.data.frame(out_social)

## plot this model run

plot(out_social$time[0:1500], out_social$n[0:1500], xlim = c(0,240),
     pch=16,  type = 'l', lwd = 2, col = 'gray',
     ylim=c(0, max(out_social$n, out_social$p1, out_social$p2)+1),
     ylab="Abundance", xlab="Time", yaxt = 'n', xaxt = 'n')
lines(out_social$time[0:1500], out_social$p1[0:1500], lwd = 2, lty = 1)
lines(out_social$time[0:1500], out_social$p2[0:1500], lwd = 2, lty = 2)
lines(out_social$time[1800:2400], out_social$n[(maa-600):maa], lwd = 2, col = 'gray',)
lines(out_social$time[1800:2400], out_social$p1[(maa-600):maa], lwd = 2, lty = 1)
lines(out_social$time[1800:2400], out_social$p2[(maa-600):maa], lwd = 2, lty = 2)

#panel C
es <- Parms[[9]]
de <- Parms[[10]]
Parms[[9]] <- es*es/(es+de)
Parms[[10]] <- de*de/(es+de)

MODEL_denovo_social <- function(time, state, parameters) {
  with(as.list(c(state, parameters)), {   
    dn <- n*r*(k-n)/k - t2(a1,h1,p1,n) - t2(a2,h2,p2,n)
    dp1 <- b1*t2(a1,h1,p1,n) + b2*t2(a2,h2,p2,n) - d*p1*n - s*n*p1*p2 - m1*p1 #change to appropriate transmission function
    dp2 <- d*p1*n + s*n*p1*p2  - m2*p2 #change to appropriate transmission function
    return(list(c(dn, dp1, dp2)))
  })
}

out_denovo_social <- ode(y=init, times=sim.time, func=MODEL_denovo_social, parms=Parms)
out_denovo_social = as.data.frame(out_denovo_social)

## plot this model run

plot(out_denovo_social$time[0:1500], out_denovo_social$n[0:1500], xlim = c(0,240),
     pch=16,  type = 'l', lwd = 2, col = 'gray',
     ylim=c(0, max(out_denovo_social$n, out_denovo_social$p1, out_denovo_social$p2)+1),
     ylab=bquote(italic('de novo')), xlab="Time", yaxt = 'n', xaxt = 'n')
lines(out_denovo_social$time[0:1500], out_denovo_social$p1[0:1500], lwd = 2, lty = 1)
lines(out_denovo_social$time[0:1500], out_denovo_social$p2[0:1500], lwd = 2, lty = 2)
lines(out_denovo_social$time[1800:2400], out_denovo_social$n[(maa-600):maa], lwd = 2, col = 'gray',)
lines(out_denovo_social$time[1800:2400], out_denovo_social$p1[(maa-600):maa], lwd = 2, lty = 1)
lines(out_denovo_social$time[1800:2400], out_denovo_social$p2[(maa-600):maa], lwd = 2, lty = 2)

text(165,-.3,'//', cex = 1, xpd = NA)

mtext('density', side = 2, line = 1.6, outer = T)
mtext('time', side = 1, line = 1.9, outer = T)
mtext(bquote(italic(dP[1])*','~italic(sP[1]*P[2])), side = 3, line = 1, outer = T, adj = 0.25)
mtext(bquote(italic(dP[1]*N)*','~italic(sP[1]*P[2]*N)), side = 3, line = 1, outer = T, adj = 0.75)
```
