## Supplementary material for "Socially transmitted innovations in dynamic predator-prey systems": All supplements: SuppFigure4.r

```
#### Predator-prey dynamics when predators use social learning ####
## Generating a bifurcation diagram

library(deSolve)

t2 <- function(a, h, p, n) a*p*n/(1 + a*h*n) 
#### sP1P2 ####
MODEL <- function(time, state, parameters) {
  with(as.list(c(state, parameters)), {   
    dn <- n*r*(k-n)/k - t2(a1,h1,p1,n) - t2(a2,h2,p2,n)
    dp1 <- b1*t2(a1,h1,p1,n) + b2*t2(a2,h2,p2,n) - s*p1*p2 - m1*p1 #change to appropriate transmission function
    dp2 <- s*p1*p2  - m2*p2 #change to appropriate transmission function
    return(list(c(dn, dp1, dp2)))
  })
}

## Specify parameter values 
Parms <- list(a1 = 0.5, 
              a2 = 0.5,
              b1 = 0.27,  
              b2 = 0.27, 
              m1 = 0.2, 
              m2 = 0.2, 
              h1 = 1, 
              h2 = 0.5,
              s = 0.2, #default 0.2 
              r = 2,   #default 2 for b1 < b2, m1 > m2
              k = 10   #default 10
)   

## Specify Initial Conditions 
n_star = with(Parms, m2*(a2*m1 - a1*m2 + r*s)/(a1*a2*(b1 - b2)*m2 + a2*b2*r*s))
p1_star = with(Parms, m2/s)
p2_star = with(Parms, (r*s-a1*m2)/(a2*s))
init = c(n = n_star, p1 = p1_star+0.1, p2 = p2_star+0.1)

## Specify simulation time
sim.time <- seq(0, 4000, by = 0.1)

## Feed model, initial conditions and parameter values into solver
out <- ode(y=init, times=sim.time, func=MODEL, parms=Parms)
out = as.data.frame(out)

maa <- dim(out)[1]
par(mfrow = c(3,1), mar = c(0,0,0,0), oma = c(2,4,0.5,0.5))
plot(out$time[0:1500], out$n[0:1500], xlim = c(0,240),
     pch=16,  type = 'l', lwd = 2, col = 'gray',
     ylim=c(0, 8),
     ylab="Abundance", xlab="Time", xaxt = 'n')
lines(out$time[0:1500], out$p1[0:1500], lwd = 2, lty = 1)
lines(out$time[0:1500], out$p2[0:1500], lwd = 2, lty = 2)
lines(out$time[1800:2400], out$n[(maa-600):maa], lwd = 2, col = 'gray',)
lines(out$time[1800:2400], out$p1[(maa-600):maa], lwd = 2, lty = 1)
lines(out$time[1800:2400], out$p2[(maa-600):maa], lwd = 2, lty = 2)
#text(165,(max(out$n, out$p1, out$p2)+0.5)/2,'...', cex = 1)
text(0,7 ,'A', cex = 1.2)

##########################################################################################
#### sNP1P2 ####
MODEL <- function(time, state, parameters) {
  with(as.list(c(state, parameters)), {   
    dn <- n*r*(k-n)/k - t2(a1,h1,p1,n) - t2(a2,h2,p2,n)
    dp1 <- b1*t2(a1,h1,p1,n) + b2*t2(a2,h2,p2,n) - s*n*p1*p2 - m1*p1 #change to appropriate transmission function
    dp2 <- s*n*p1*p2  - m2*p2 #change to appropriate transmission function
    return(list(c(dn, dp1, dp2)))
  })
}

## Feed model, initial conditions and parameter values into solver
out <- ode(y=init, times=sim.time, func=MODEL, parms=Parms)
out = as.data.frame(out)

maa <- dim(out)[1]

plot(out$time[0:1500], out$n[0:1500], xlim = c(0,240),
     pch=16,  type = 'l', lwd = 2, col = 'gray',
     ylim=c(0, 8),
     ylab="Abundance", xlab="Time", xaxt = 'n')
lines(out$time[0:1500], out$p1[0:1500], lwd = 2, lty = 1)
lines(out$time[0:1500], out$p2[0:1500], lwd = 2, lty = 2)
lines(out$time[1800:2400], out$n[(maa-600):maa], lwd = 2, col = 'gray',)
lines(out$time[1800:2400], out$p1[(maa-600):maa], lwd = 2, lty = 1)
lines(out$time[1800:2400], out$p2[(maa-600):maa], lwd = 2, lty = 2)
#text(165,(max(out$n, out$p1, out$p2)+0.5)/2,'...', cex = 1)
text(0,7 ,'B', cex = 1.2)

##########################################################################################


#### sa2NP1P2/(1+a2h2N) ####
MODEL <- function(time, state, parameters) {
  with(as.list(c(state, parameters)), {   
    dn <- n*r*(k-n)/k - t2(a1,h1,p1,n) - t2(a2,h2,p2,n)
    dp1 <- b1*t2(a1,h1,p1,n) + b2*t2(a2,h2,p2,n) - s*p1*t2(a2,h2,p2,n) - m1*p1 #change to appropriate transmission function
    dp2 <- s*p1*t2(a2,h2,p2,n)  - m2*p2 #change to appropriate transmission function
    return(list(c(dn, dp1, dp2)))
  })
}

## Feed model, initial conditions and parameter values into solver
out <- ode(y=init, times=sim.time, func=MODEL, parms=Parms)
out = as.data.frame(out)

plot(out$time[0:1500], out$n[0:1500], xlim = c(0,240),
     pch=16,  type = 'l', lwd = 2, col = 'gray',
     ylim=c(0, 8),
     ylab="Abundance", xlab="Time", xaxt = 'n')
lines(out$time[0:1500], out$p1[0:1500], lwd = 2, lty = 1)
lines(out$time[0:1500], out$p2[0:1500], lwd = 2, lty = 2)
lines(out$time[1800:2400], out$n[(maa-600):maa], lwd = 2, col = 'gray',)
lines(out$time[1800:2400], out$p1[(maa-600):maa], lwd = 2, lty = 1)
lines(out$time[1800:2400], out$p2[(maa-600):maa], lwd = 2, lty = 2)
text(165,-.3,'//', cex = 1, xpd = NA)
text(0,7,'C', cex = 1.2)
mtext('time', side = 1, line = 0.4, outer = T)
mtext('density', side = 2, line = 2.4, outer = T)
```
