## Supplementary material for "Socially transmitted innovations in dynamic predator-prey systems": All supplements: SuppFigure6.R

```
## Supplementary Figure 4 - A visualization of R0 values relative to
##                          the prey population size (N)


t2 <- function(a, h, p, n) a*p*n/(1 + a*h*n) 
m3 <- function(x){ 
  a <- max(x)
  b <- min(x)
  z <- mean(x)
  c(a,b,z)
}

beg <- function(parama,offset){
  a1 = parama[[1]]; a2 = parama[[2]]; b1 = parama[[3]]; b2 = parama[[4]];
  m1 = parama[[5]]; m2 = parama[[6]]; h1 = parama[[7]]; h2 = parama[[8]];
  s = parama[[9]]; r = parama[[10]]; k = parama[[11]]; 
  n_star = m1/(a1*(b1-h1*m1))
  p1_star = b1*r*(a1*b1*k-m1-a1*h1*k*m1)/(a1^2 * k*(b1-h1*m1)^2)
  p2_star = 0
  init = c(n = n_star+offset, p1 = p1_star+offset, p2 = p2_star)
  init
}

nnull <- function(parama,x){
  a1 = parama[[1]]; a2 = parama[[2]]; b1 = parama[[3]]; b2 = parama[[4]];
  m1 = parama[[5]]; m2 = parama[[6]]; h1 = parama[[7]]; h2 = parama[[8]];
  s = parama[[9]]; r = parama[[10]]; k = parama[[11]]; 
  ((k - x)*(1 + a1*h1*x)*r)/(a1*k)
}

pnull <- function(parama){
  a1 = parama[[1]]; a2 = parama[[2]]; b1 = parama[[3]]; b2 = parama[[4]];
  m1 = parama[[5]]; m2 = parama[[6]]; h1 = parama[[7]]; h2 = parama[[8]];
  s = parama[[9]]; r = parama[[10]]; k = parama[[11]]; 
  m1/(a1*(b1 - h1*m1))
}

R0_two <- function(parama){
  a1 = parama[[1]]; a2 = parama[[2]]; b1 = parama[[3]]; b2 = parama[[4]];
  m1 = parama[[5]]; m2 = parama[[6]]; h1 = parama[[7]]; h2 = parama[[8]];
  s = parama[[9]]; r = parama[[10]]; k = parama[[11]]; 
  nstar <- m1/(a1*(b1-h1*m1))
  p1star <- b1*r*(a1*b1*k-m1-a1*h1*k*m1)/(a1^2 * k*(b1-h1*m1)^2)
  s*p1star/m2
}

R0_three <- function(parama){
  a1 = parama[[1]]; a2 = parama[[2]]; b1 = parama[[3]]; b2 = parama[[4]];
  m1 = parama[[5]]; m2 = parama[[6]]; h1 = parama[[7]]; h2 = parama[[8]];
  s = parama[[9]]; r = parama[[10]]; k = parama[[11]]; 
  nstar <- m1/(a1*(b1-h1*m1))
  p1star <- b1*r*(a1*b1*k-m1-a1*h1*k*m1)/(a1^2 * k*(b1-h1*m1)^2)
  s*p1star*nstar/m2
}

R0_amy <- function(parama){
  a1 = parama[[1]]; a2 = parama[[2]]; b1 = parama[[3]]; b2 = parama[[4]];
  m1 = parama[[5]]; m2 = parama[[6]]; h1 = parama[[7]]; h2 = parama[[8]];
  s = parama[[9]]; r = parama[[10]]; k = parama[[11]]; 
  nstar <- m1/(a1*(b1-h1*m1))
  p1star <- b1*r*(a1*b1*k-m1-a1*h1*k*m1)/(a1^2 * k*(b1-h1*m1)^2)
  s*a2*p1star*nstar/((1+a2*h2*nstar)*m2)
}

Parms <- list(a1 = 0.5, 
              a2 = 0.5,
              b1 = 0.27,  
              b2 = 0.27, 
              m1 = 0.2, 
              m2 = 0.2, 
              h1 = 1, 
              h2 = 1,
              s = 0.2, #default 0.2 
              r = 2,    #default 2 for b1 < b2, m1 > m2
              k = 10   #default 10
)   

#variable b1 assimilation rate
vs <- seq(0.24,0.5,length.out=30)
x <- rep(NA,30)
y1 <- rep(NA,30)
y2 <- rep(NA,30)
y3 <- rep(NA,30)
for(i in 1:length(vs)){
  Parms[[3]] <- vs[i]
  x[i] <- beg(Parms,0)[1]
  y1[i] <- R0_two(Parms)
  y2[i] <- R0_three(Parms)
  y3[i] <- R0_amy(Parms)
}
par(mfrow=c(1,1), oma = c(0,0,0,0),mar=c(4,5,1,1))
plot(x, y1/max(y1), type = 'l', ylim = c(0,1),  ylab = 'relative R0', xlab = '')
lines(x, y2/max(y2), lty = 2)
lines(x, y3/max(y3), lty = 3)

legend('topleft',c('sP1P2','sNP1P2','sa2NP1P2/(1+a2*h2*N)'),lty=c(1,2,3))

#variable h1 handling time
Parms[[3]] <- 0.27
vs <- seq(0.01,1.15,length.out=30)
x <- rep(NA,30)
y1 <- rep(NA,30)
y2 <- rep(NA,30)
y3 <- rep(NA,30)
for(i in 1:length(vs)){
  Parms[[7]] <- vs[i]
  x[i] <- beg(Parms,0)[1]
  y1[i] <- R0_two(Parms)
  y2[i] <- R0_three(Parms)
  y3[i] <- R0_amy(Parms)
}
plot(x, y1/max(y1), type = 'l', ylim = c(0,1),  ylab = 'relative R0', xlab = 'n*')
lines(x, y2/max(y2), lty = 2)
lines(x, y3/max(y3), lty = 3)

legend('topleft',c('sP1P2','sNP1P2','sa2NP1P2/(1+a2*h2*N)'),lty=c(1,2,3))


#variable a1 attack rate
Parms[[7]] <- 1
vs <- seq(0.28,2,length.out=30)
x <- rep(NA,30)
y1 <- rep(NA,30)
y2 <- rep(NA,30)
y3 <- rep(NA,30)
for(i in 1:length(vs)){
  Parms[[1]] <- vs[i]
  x[i] <- beg(Parms,0)[1]
  y1[i] <- R0_two(Parms)
  y2[i] <- R0_three(Parms)
  y3[i] <- R0_amy(Parms)
}
plot(x, y1/max(y1), type = 'l', ylim = c(0,1),  
     ylab = bquote('relative'~italic(R)[0]), 
                   xlab = expression(italic(N)^"*"))
lines(x, y2/max(y2), lty = 2)
lines(x, y3/max(y3), lty = 3)

#legend('topleft',c('sP1P2','sNP1P2','sa2NP1P2/(1+a2*h2*N)'),lty=c(1,2,3))


  
```
